# AKAP1 regulates mitochondrial and synaptic homeostasis to enable neuroprotection and repair in retinal ganglion cell degeneration

**DOI:** 10.1101/2025.11.24.689866

**Authors:** Tonking Bastola, Keun-Young Kim, Ziyao Shen, Guy A. Perkins, Muna Poudel, Veronica Gomez, Yoonjin Lim, Hyejeong Choi, Jae Yon Won, Soo-Ho Choi, Mark H. Ellisman, Robert N. Weinreb, Won-Kyu Ju

**Affiliations:** Viterbi Family Department of Ophthalmology, Viterbi Family Vision Research Center, Shiley Eye Institute, University of California San Diego, La Jolla, CA 92039, USA; National Center for Microscopy and Imaging Research, Department of Neurosciences, University of California San Diego, La Jolla, CA 92039, USA; Institute of Engineering in Medicine, Shu Chien – Gene Lay Department of Bioengineering, University of California San Diego, La Jolla, CA 92039, USA; Department of Ophthalmology and Visual Science, Eunpyeong St. Mary’s Hospital, College of Medicine, The Catholic University of Korea, Seoul, Republic of Korea; Department of Medicine, University of California San Diego, La Jolla, CA 92039, USA

## Abstract

Glaucoma is a leading cause of irreversible blindness, characterized by progressive retinal ganglion cell (RGC) loss and optic nerve degeneration. Mitochondrial dysfunction plays a central role in this neurodegeneration, yet effective targeted therapies remain limited. Here, we identify the mitochondrial scaffold A-kinase anchoring protein 1 (AKAP1) as a critical regulator of RGC resilience and axon regeneration. AKAP1 expression is diminished in human glaucomatous retinas and experimental glaucoma models, correlating with elevated intraocular pressure, disrupted mitochondrial dynamics, oxidative stress, and synaptic instability. Restoration of AKAP1 via adeno-associated virus serotype 2-mediated gene therapy preserves RGC survival, promotes mitochondrial fusion and cristae integrity, enhances ATP production, and mitigates oxidative and apoptotic stress in mouse models of glaucoma and optic nerve injury. Transcriptomic profiling of AKAP1 knockout retinas reveals widespread dysregulation of mitochondrial and synaptic gene networks. Mechanistically, AKAP1 stabilizes synapses by promoting mitochondrial biogenesis, modulating calcium/calmodulin-dependent kinase II and synapsin phosphorylation, maintaining synaptophysin expression, and suppressing complement component C1q expression, thereby preventing early synaptic loss in glaucomatous neurodegeneration. Moreover, restoring AKAP1 expression facilitates axonal regeneration, preserves the central visual pathway, and maintains visual function. Collectively, these findings establish AKAP1 as a master regulator of mitochondrial and synaptic homeostasis and axonal regeneration and a promising therapeutic target for vision preservation in glaucomatous neurodegeneration.

**One Sentence Summary:** AKAP1 protects retinal ganglion cells and preserves vision by restoring mitochondrial and synaptic health in experimental glaucoma models.

## INTRODUCTION

Glaucoma, a leading cause of blindness worldwide, is marked by progressive optic nerve (ON) degeneration and retinal ganglion cell (RGC) death. The pathogenesis involves elevated intraocular pressure (IOP), oxidative stress, and notably, mitochondrial dysfunction—particularly defects in oxidative phosphorylation (OXPHOS) linked to cytochrome c oxidase subunit I polymorphisms (*1–5*). Lowering IOP slows disease progression, but does not halt it, highlighting the need for alternative neuroprotective strategies. Multiple studies corroborate mitochondrial impairment as a driver of glaucomatous neurodegeneration (*2, 6, 7*), yet the molecular pathways linking elevated IOP, mitochondrial dysfunction, and RGC loss remain incompletely defined.

A-kinase anchoring protein 1 (AKAP1) is a mitochondrial scaffold protein that anchors protein kinase A (PKA) and key signaling molecules at the outer mitochondrial membrane (OMM), orchestrating pathways that regulate mitochondrial dynamics and function (*2, 8, 9*). AKAP1 protects against cerebral ischemic injury by maintaining respiratory chain activity, inhibiting DRP1-mediated mitochondrial fission, reducing superoxide production, and modulating Ca²⁺ signaling (*10*). Our previous work showed that glaucomatous RGCs have reduced AKAP1 expression, heightened calcineurin (CaN) activity, and diminished dynamin-related protein 1 (DRP1) serine S637 (S637) phosphorylation, disrupting OXPHOS and promoting mitochondrial fragmentation and RGC death (*8, 11*). Evidence suggests synaptic deficits in the inner retina precede RGC soma loss (*12–14*). Synapses in the inner plexiform layer (IPL), particularly those of bipolar, amacrine, and RGCs, are susceptible to glaucomatous damage due to high metabolic demands and vulnerability to oxidative stress (*13–16*). Early synaptic “pruning” further disrupts neurotransmission, increasing RGC susceptibility. Mitochondrial dynamics, regulated by AKAP1, are essential for synaptic architecture and function (*17, 18*), yet the role of AKAP1 in synaptic activity during glaucoma remains unexplored.

Here, we investigate the impact of AKAP1 on retinal synaptic function, RGC survival, and axon regeneration in glaucomatous and traumatic optic neuropathy models. Experimental glaucoma paradigms demonstrate reduced AKAP1 and PKA in RGCs, correlating with neurodegeneration. Restoring AKAP1 via adeno-associated virus serotype 2 (AAV2) delivery preserves RGC survival, visual pathway integrity, and function; conversely, AKAP1 knockout (*Akap1^−/−^)* mice show worsened RGC dysfunction. Mechanistically, AKAP1 restoration maintains mitochondrial dynamics, adenosine triphosphate (ATP) production, and resistance to stress and apoptosis, while preserving synaptic activity via enhanced mitochondrial biogenesis. In glaucomatous and injury models, AKAP1 promotes axon regeneration and vision restoration. Our results position AKAP1 as a key regulator of mitochondrial integrity and neuronal resilience, supporting its therapeutic potential for neuroprotection and vision restoration in glaucomatous and traumatic optic neuropathies.

## RESULTS

### AKAP1 expression is reduced in glaucomatous human RGCs

AKAP1 is expressed in RGCs and reduced by elevated IOP in experimental glaucoma models (*11*). To assess the role of AKAP1 in human glaucoma, we analyzed donor retinas via immunohistochemistry. Our previous report confirmed the diagnosis of glaucoma diagnosis in donor retinas based on documented, marked loss of RGCs and axons in the optic nerve head (ONH) (*19*). In controls, AKAP1 and PKA localized to RGC somas, co-staining with neuron-specific β-III tubulin (TUJ1) (Fig. 1A-D). In glaucomatous retinas, AKAP1 was significantly increased in the total retina and ganglion cell layer (GCL), but specifically decreased in RGC somas (Fig. 1A and B; fig. S1). PKA was markedly reduced in both GCL and RGC somas, with a significant overall decrease in glaucomatous retinas compared to controls (Fig. 1C and D; fig. S1). Additionally, AKAP1 upregulation was predominantly observed in glutamine synthase (GS)-positive Müller glia (fig. S2A), indicating a glial response to glaucomatous stress. To further characterize this, we exposed cultured Müller glia to elevated hydrostatic pressure (HP, 30 mmHg) for 24 h (fig. S2B), which significantly increased AKAP1 expression and cell viability without cytotoxicity (fig. S2C-E). These results suggest that AKAP1 may support Müller glia survival under glaucomatous conditions, potentially benefiting neuronal health.

**Fig. 1.**
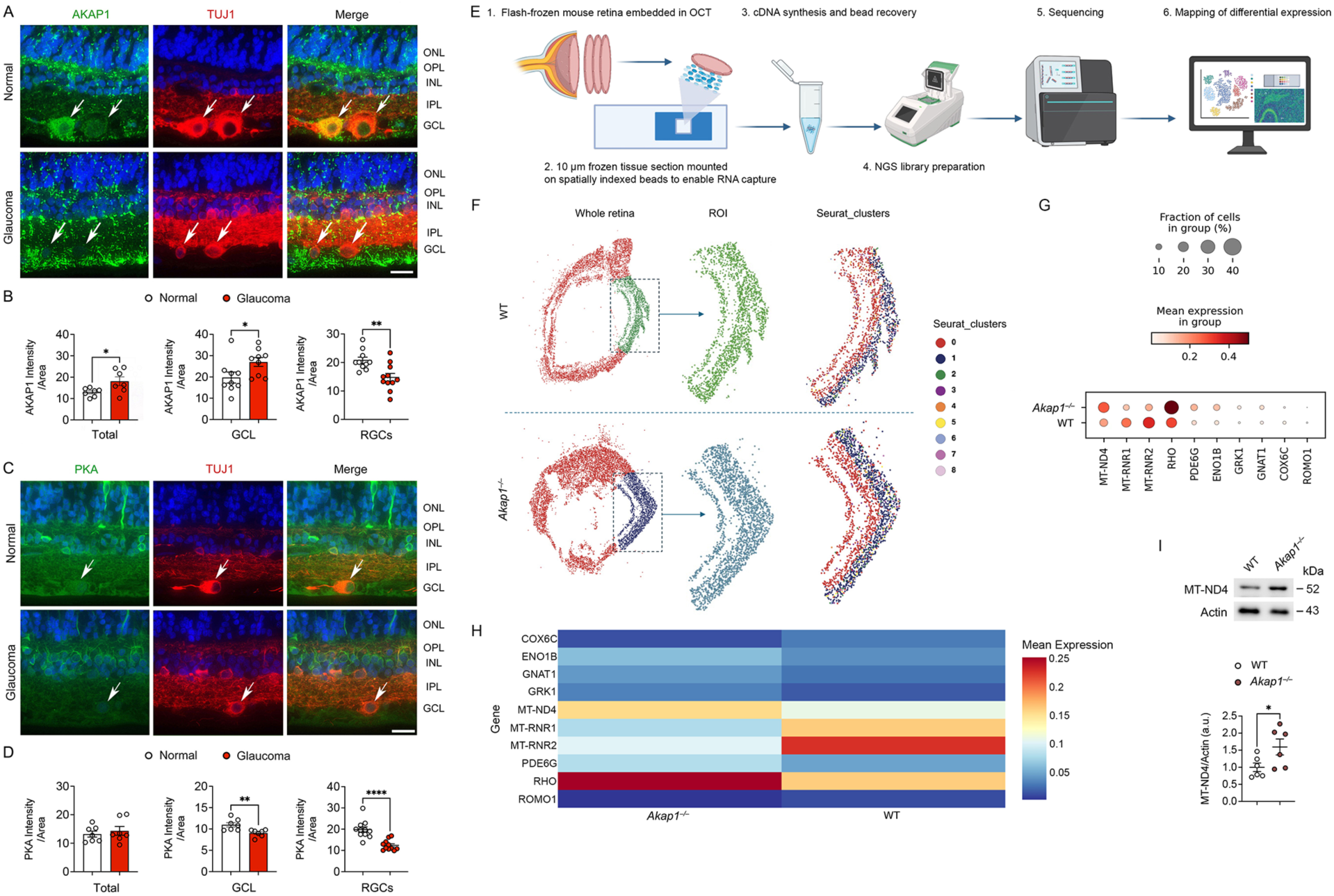
AKAP1 and PKA expression in glaucomatous human retina. (A) Representative retinal image for AKAP1 (green) and TUJ1 (red) immunoreactivities. Arrows indicate AKAP1 immunoreactivity co-labeled with TUJ1 in RGC somas. (B) Quantitative fluorescent intensity of AKAP1 immunoreactivity in the total and GCL layers, as well as in the RGC somas of the glaucomatous human retina (*n* = 7 retinal sections from 7 eyes from donors/group; *n* = 10 to 11 RGCs from 7 eyes from donors/group). (C) Representative retinal image for PKA (green) and TUJ1 (red) immunoreactivities. (D) Quantitative fluorescent intensity of PKA immunoreactivity in the total and GCL layers, as well as in the RGC somas of the glaucomatous human retina (*n* = 7 retinal sections from 7 eyes from donors/group; *n* = 11 RGCs from 7 eyes from donors/group). Arrows indicate PKA immunoreactivity co-labeled with TUJ1 in RGC somas. Spatial transcriptomic profiling and MT-ND4 expression in WT and *Akap1^−/−^* retinas. (E) Schematic diagram for experimental workflow: Retinas from 10-month-old WT and *Akap1^−/−^* mice were flash-frozen in OCT and processed on Curio Seeker 3×3 slides, followed by library preparation, sequencing, and LatchBio analysis. (F) Spatial map of retinal sections showing selected regions of interest (ROIs) and Seurat clusters (0–8). (G) Dot plot of top differentially expressed genes, highlighting upregulated mitochondrial and phototransduction genes (*MT-ND4, RHO, PDE6B, GRK1*) and downregulated mitochondrial ribosomal genes (*MT-RNR1, MT-RNR2, ROMO1*) in *Akap1^−/−^* retinas. (H) Heatmap showing spatial expression of the top 10 DEGs between WT and *Akap1^−/−^* retina. (I) Western blot analysis for the expression level of MT-ND4 protein in WT and *Akap1^−/−^* retina. Error bars represent SEM. Statistical analysis was performed using an unpaired Student’s *t*-test. \**P* < 0.05, \*\**P* < 0.01, and \*\*\*\**P* < 0.0001. Scale bars, 20 μm.

### Transcriptomic analysis of AKAP1-deficient retinas reveals mitochondrial and synaptic dysregulation

To assess transcriptomic consequences of AKAP1 deficiency in the retina, we performed spatial transcriptomic and bulk RNA sequencing analyses on WT and *Akap1^−/−^* mouse retinas. Spatial transcriptomic analysis was performed following the experimental workflow (Fig. 1E) and revealed nine transcriptionally distinct clusters (0–8) across the retinal section of wild-type (WT) and *Akap1^−/−^* mice (Fig. 1F; fig. S2A). Differential gene expression (DEG) analysis by spatial transcriptomic approach demonstrated a distinct molecular signature in the retina of *Akap1^−/−^*mice, with upregulation of mitochondrial and phototransduction genes, including *MT-ND4*, *RHO*, *PDE6B*, and *GRK1*, and downregulation of mitochondrial ribosomal and oxidative stress–related genes, including *MT-RNR1*, *MT-RNR2*, and *ROMO1* (Fig. 1G and H; fig. S2B). These findings indicate altered mitochondrial function and metabolic stress. Consistently, Western blot analysis further confirmed increased MT-ND4 protein levels in *Akap1^−/−^* retinas compared to WT (Fig. I). Additionally, DEG analysis by bulk RNA sequencing identified 440 significantly dysregulated genes (244 up, 196 down; fold change ≥1.25, adjusted *P*<0.05) (fig. S3). Heatmap-based unsupervised clustering revealed clear segregation between WT and *Akap1^−/−^* retina samples (fig. S4A). Gene Ontology analysis showed enrichment for mitochondrial and synaptic genes among the DEGs (fig. S4B), linking AKAP1 loss to dysregulated energy metabolism and neuronal signaling. Notably, the anti-apoptotic gene Bcl2 and mitochondrial regulator Fbxl4 were significantly upregulated, potentially reflecting compensatory neuroprotective responses in the absence of AKAP1 (*20, 21*). In contrast, synaptic and neurotrophic genes such as BDNF, Gabrr2, Nos1, and others were markedly reduced. Lower BDNF levels suggest impaired RGC survival and synaptic maintenance (*22*), while Gabrr2 downregulation indicates altered inhibitory neurotransmission (*23*), a feature shared with glaucomatous degeneration. Genes related to cytoskeletal remodeling, including Actg1 and Sema4C, were upregulated, implying adaptive structural responses (fig. S4B) (*24*). Pathway enrichment using Hallmark and WikiPathways datasets further highlighted deregulation of neurogenesis, synaptic vesicle cycling, mitochondrial dynamics, and neurodegenerative pathways (fig. S4C). Together, these results demonstrate that AKAP1 deficiency profoundly alters retinal transcriptomic networks critical for mitochondrial function and neuronal connectivity.

### Restoring AKAP1 expression protects RGCs and preserves visual function in glaucomatous D2 mice

To test whether restoring AKAP1 can promote neuroprotection in glaucoma, we administered AAV2-AKAP1 or control (AAV2-Null) intravitreally to 5-month-old DBA/2J (D2) mice, a spontaneous glaucoma model (*25, 26*), and evaluated outcomes at 10 months (Fig. 2A). Age-matched D2-*Gpnmb⁺* (D2G-Null) mice served as controls (*27*). D2 mice showed significantly elevated IOP, and both AKAP1 and the RGC marker, RNA-binding protein with multiple splicing (RBPMS), were reduced in glaucomatous retinas (fig. S6A and B); for consistency, only D2 mice with IOPs between 20–35 mmHg were included (Fig. 2B). RBPMS-positive RGC loss was prominent in D2-Null but prevented in D2-AKAP1 retinas, as confirmed by immunostaining and western blot for His-tag and AKAP1 (Fig. 2C-E; fig. S6C and D). AKAP1 and PKA expression in RGCs and inner retina, both of which were diminished in D2-Null, were restored by AAV2-AKAP1 (Fig. 2F-I). Given the elevated AKAP1 in stressed Müller glia in human and *in vitro* (fig. S3), we examined its expression in D2 retina. AKAP1 was reduced in Müller glia of D2-Null and rescued by AAV2-AKAP1 (fig. S7), suggesting glial maintenance may be supported by improved RGC health. RGC survival was significantly enhanced in D2-AKAP1 vs. D2-Null mice (Fig. 2J and K), and restored superior colliculus (SC) innervation was confirmed by cholera toxin subunit B (CTB) labeling (Fig. 2L and M). Electrophysiological analysis showed improved pattern electroretinogram (pERG) and pattern visual evoked potential (pVEP) amplitudes, reflecting enhanced RGC and ON function, in D2-AKAP1, while pVEP latency was unchanged (Fig. 2N and O). Thus, AKAP1 reconstitution enhances RGC survival, retina-brain connectivity, and visual function in glaucomatous degeneration (Fig. 2P).

**Fig. 2.**
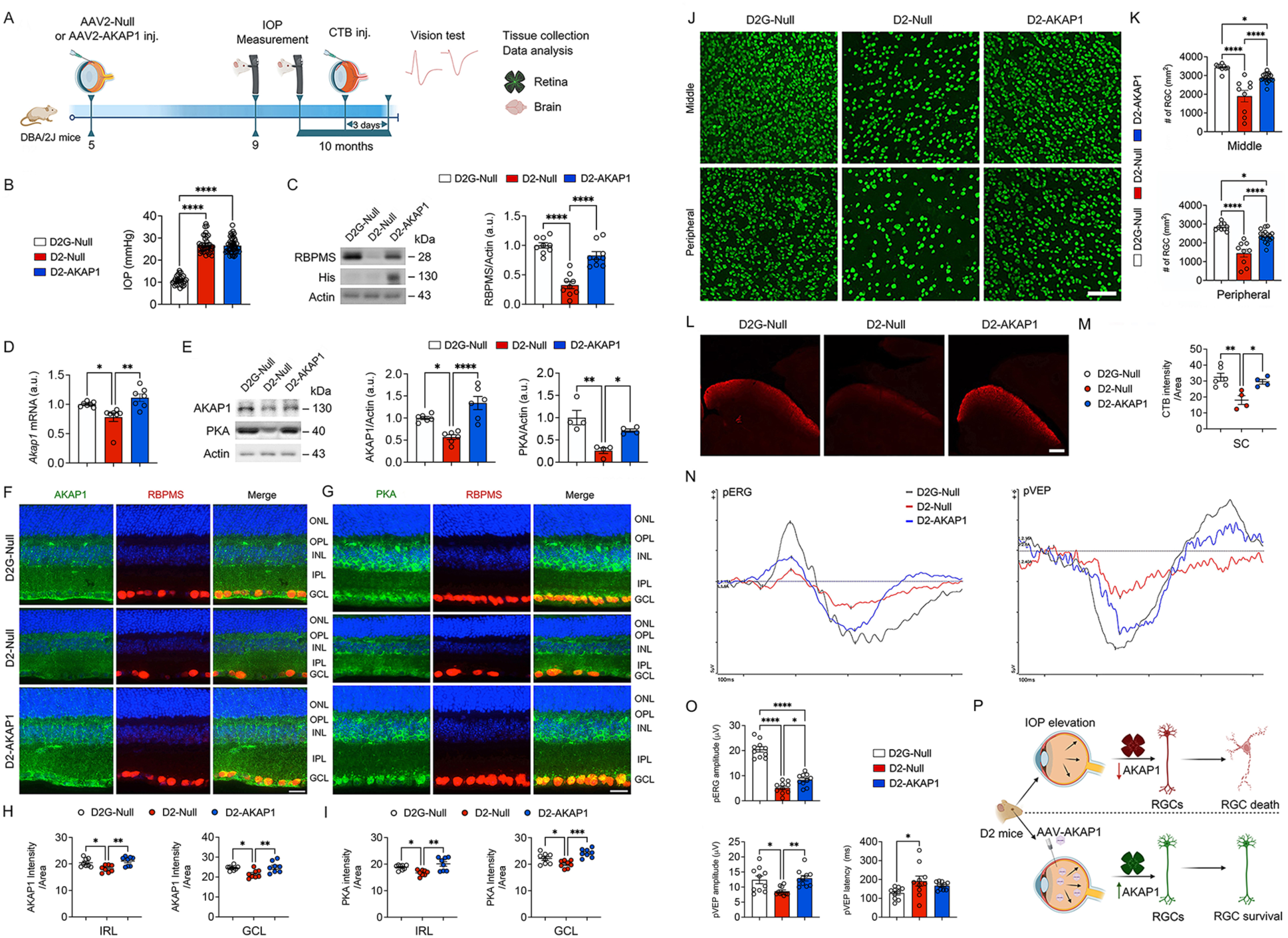
Restoring AKAP1 expression protects RGCs and preserves visual function in glaucomatous D2 mice. (A) Experimental schematic and timeline of AAV injection, IOP measurement, tissue collection, and data analysis in D2G and D2 mice (*n* = 20 mice per group). (B) IOP measurement in 10-month-old D2G-Null, D2-Null, and D2-AKAP1 mice (*n* = 20 mice per group). (C) Representative image of a western blot and densitometry graph for RBPMS and His expression in the retinas (*n* = 9 retinas per group). (D) Quantitative PCR for AKAP1 gene expression in the retinas (*n* = 7 retinas per group). (E) Representative image of a western blot and densitometry graph for AKAP1 and PKA expression in the retinas (*n* = 4 to 6 retinas per group). (F and G) Representative retinal images for AKAP1 (green), PKA (green), and RBPMS (red) immunoreactivities. (H and I) Quantitative fluorescent intensity of AKAP1 immunoreactivity in the inner retinal layer (IRL) and GCL (*n* = 8 retinas per group). (J) Representative retina whole-mounted images for RBPMS (green)-positive RGCs in the middle and peripheral areas of the retina. (K) Quantitative analysis of RGC numbers in the middle and peripheral areas of the retina (*n* = 9 to 18 retina wholemounts per group). (L) Representative images for CTB labeling in the SC. (M) Quantitative analysis of CTB intensity in the SC (*n* = 4 to 6 SC sections from 3 mice per group). (N) Representative recording graphs for pERG and pVEP measurements. (O) Quantitative analysis of pERG amplitude, pVEP amplitude, and pVEP latency (*n* = 10 mice per group). Note that AKAP1 expression restores visual function in glaucomatous D2 mice. (P) Schematic for the effect of sustained AKAP1 expression on glaucomatous retina. Error bars represent SEM. Statistical analysis was performed using one-way ANOVA and Tukey’s multiple comparisons test. \**P* < 0.05, \*\**P* < 0.01, and \*\*\*\**P* < 0.0001. Scale bars, 20 μm (F and G), 50 μm (J), 100 μm (L).

### Restoring AKAP1 expression protects RGCs and preserves visual function in a mouse model of microbead-induced ocular hypertension

To further examine the neuroprotective role of AKAP1, we employed a microbead (MB)-induced ocular hypertension glaucoma model (*28, 29*). Three-month-old C57BL/6J mice received intravitreal AAV2-Null or AAV2-AKAP1 injections three weeks before MB administration (Fig. 3A). IOP was tracked weekly, and mice with IOPs of 20-30 mmHg were included; IOP levels did not differ between Null-MB and AKAP1-MB groups (Fig. 3B). MB-induced IOP elevation led to significant RGC loss in Null-MB compared to Null-control (CNT) mice (Fig. 3C and D), while AAV2-AKAP1 preserved RGCs in MB-injected eyes. Functionally, MB-induced ocular hypertension impaired RGC and ON function, reducing pERG and pVEP amplitudes and prolonging pVEP latency (Fig. 3E and F). Remarkably, AAV2-AKAP1 restored pERG and pVEP amplitudes and normalized pVEP latency, reflecting improved visual function. Together, these data demonstrate that AKAP1 restoration protects RGCs and counteracts visual impairment induced by elevated IOP in experimental glaucoma.

**Fig. 3.**
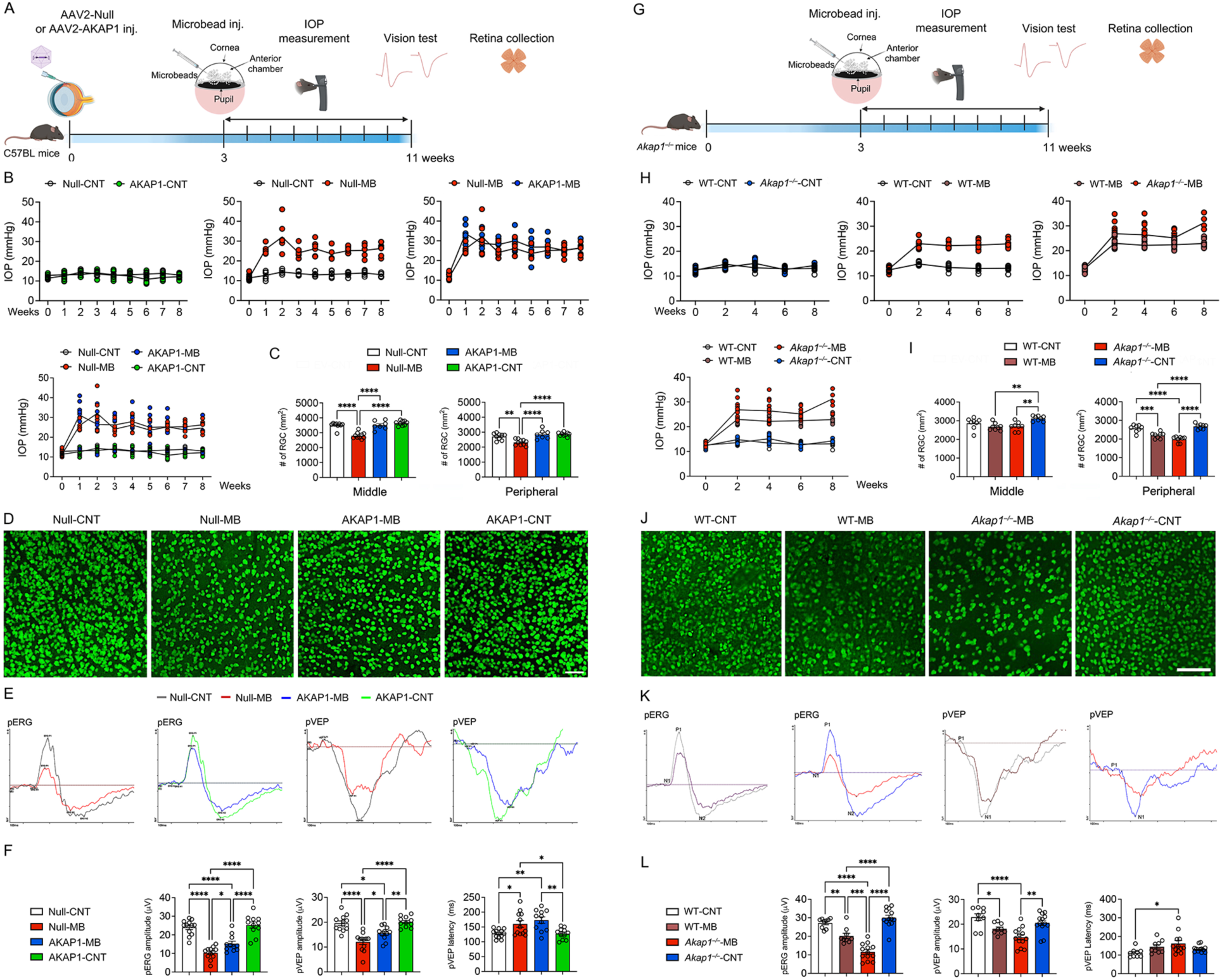
Restoring AKAP1 expression protects RGCs and preserves visual function, but AKAP1 deficiency worsens RGC function in glaucomatous MB mice. (A) Experimental schematic and timeline of AAV injection, IOP measurement, tissue collection, and data analysis in glaucomatous MB mice (*n* = 12-14 mice per group). (B) Monitoring of IOP elevation in control and glaucomatous MB mice (*n* = 12 to 14 mice per group). (C) Quantitative analysis of RGC number in the middle and peripheral areas of the retina (*n* = 7-10 retina wholemounts per group). (D) Representative whole-mounted retina images for RBPMS (green) immunoreactivity in the middle area. (E) Representative recording graphs for pERG and pVEP measurements. (F) Quantitative analysis of pERG amplitude, pVEP amplitude, and pVEP latency (*n* = 12 to 14 mice per group). Note that AKAP1 expression restores visual function in glaucomatous MB mice. (G) Experimental schematic and timeline of MB injection, IOP measurement, tissue collection, and data analysis in glaucomatous *Akap1⁻^/^⁻* MB mice (*n* = 9 mice per group). (H) Monitoring of IOP elevation in *Akap1⁻^/^⁻* MB mice (*n* = 9 to 12 mice per group). (I) Quantitative analysis of RGC number in the middle and peripheral areas of the retina (*n* = 8 retina wholemounts per group). (J) Representative whole-mounted retina images for RBPMS (green) immunoreactivity in the middle area from *Akap1⁻^/^⁻* MB mice. (K) Representative recording graphs for pERG and pVEP measurements. (L) Quantitative analysis of pERG amplitude, pVEP amplitude, and pVEP latency (*n* = 9 to 13 mice per group). Note that AKAP1 deficiency worsens RGC dysfunction in glaucomatous MB mice. Error bars represent SEM. Statistical analysis was performed using one-way ANOVA and Tukey’s multiple comparisons test. \**P* < 0.05, \*\**P* < 0.01, \*\*\**P* < 0.001, and \*\*\*\**P* < 0.0001. Scale bars, 50 μm (D) and 100 μm (J).

### AKAP1 deficiency worsens RGC loss and visual dysfunction in a mouse model of microbead-induced ocular hypertension

To determine if AKAP1 loss exacerbates RGC degeneration under glaucomatous stress, we injected microbeads (MB) into three-month-old *Akap1⁻^/^⁻* mice and monitored IOP for 8 weeks (Fig. 3G). IOP elevation did not differ between WT-MB and *Akap1⁻^/^⁻* –MB groups (Fig. 3H). MB injection reduced RGC counts in WT-MB compared to WT-CNT mice, but RGC numbers were similar between WT-MB and *Akap1⁻^/^⁻* –MB (Fig. 3H-J). Analysis of visual function revealed that WT-MB mice had reduced pERG and pVEP amplitudes, and prolonged pVEP latency (Fig. 3K and L). Notably, AKAP1 deficiency further impaired RGC function, as evidenced by a greater decline in pERG amplitude, though pVEP metrics were not significantly different (Fig. 3K and L). Collectively, these data suggest AKAP1 loss selectively worsens RGC function and may contribute to visual deficits under glaucomatous conditions.

### Restoring AKAP1 expression promotes mitochondrial fusion activity in the retina of glaucomatous D2 mice

Since AKAP1 deficiency triggers mitochondrial fragmentation via CaN activation and decreased phosphorylation of DRP1 (pDRP1) S637 (*8, 11*), we assessed whether AKAP1 restoration preserves mitochondrial dynamics in glaucomatous D2 retinas. CaN level was elevated in D2-Null but normalized with AKAP1 expression (Fig. 4A). *Drp1* mRNA was upregulated in D2-Null and significantly reduced with AKAP1 rescue (Fig. 4B). Western blotting showed reduced total DRP1 protein but increased pDRP1 S637 in D2-AKAP1 versus D2-Null, with unchanged pDRP1 S616 (Fig. 4C). Immunostaining revealed decreased DRP1 in RGCs and the ganglion cell layer following AKAP1 restoration (Fig. 4D and E). Optic atrophy type 1 (*Opa1*) mRNA and total OPA1 protein, which were reduced in D2-Null, were restored by AKAP1, though OPA1 isoform profiles were unaltered (Fig. 4F and G). Mitofusin 2 (MFN2) level, which was reduced in D2-Null, was restored by AKAP1, whereas MFN1 mRNA expression remained unchanged. Protein levels of both MFN1 and MFN2 were unaffected (Fig. 4H and I). Together, these data indicate that AKAP1 restoration promotes mitochondrial fusion by modulating DRP1 pS637 and OPA1, thereby preserving mitochondrial integrity in glaucomatous retinas.

**Fig. 4.**
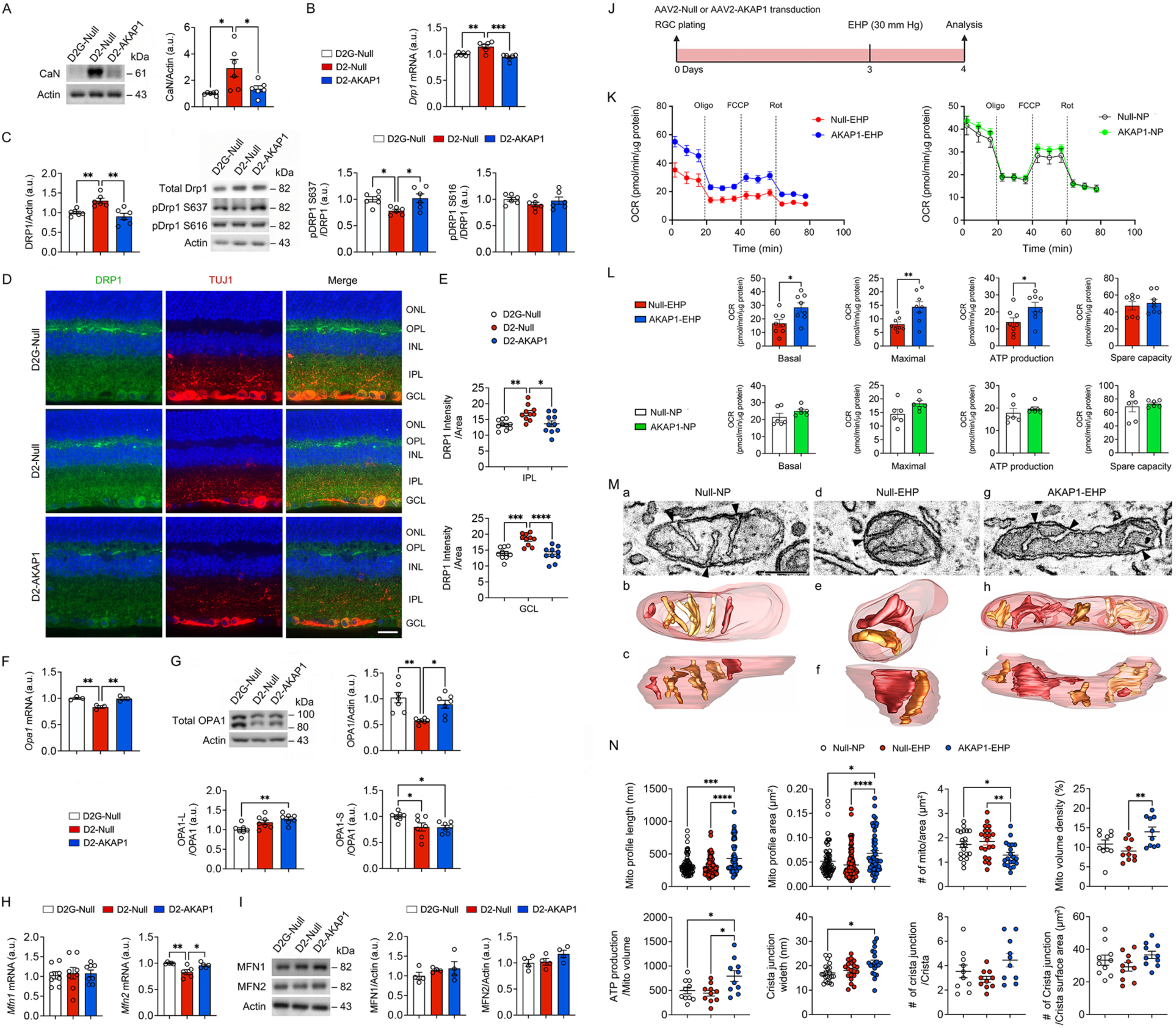
Restoring AKAP1 expression enhances mitochondrial fusion activity and energy production in glaucomatous RGCs. (A) Representative image of a western blot and densitometry graph for CaN expression in the retinas of D2G-Null, D2-Null, and D2-AKAP1 mice (*n* = 6 retinas per group). (B) Quantitative PCR for *Drp1* gene expression in the retinas. (C) Representative image of a western blot and densitometry graph for total DRP1, phospho-DRP1 (S637), and phospho-DRP1 (S616) in the retinas (*n* = 6 to 8 retinas per group). (D) Representative retinal images for DRP1 (green) and TUJ1 (red) immunoreactivities. (E) Quantitative fluorescent intensity of DRP1 immunoreactivity in the GCL of the retinas (*n* = 10 retina sections from 3 mice per group). (F) Quantitative PCR for *Opa1* gene expression in the retinas (*n* = 3 retinas from 3 mice per group). (G) Representative image of a western blot and densitometry graph for total OPA1, OPA1-S form, and OPA1-L form in the retinas (*n* = 7 retinas per group). (H) Quantitative PCR for *Mfn1*/*2* gene expression (*n* = 5 to 8 retinas per group). (I) Representative image of a western blot and densitometry graph for MFN1 and MFN2 in the retinas of (*n* = 4 retinas per group). (J) Experimental schematic and timeline of AAV2 transduction, and data analysis in RGCs under elevated HP conditions *in vitro*. (K) Representative graphs for OCR changes in RGCs under elevated HP conditions. (L) Quantitative analysis of OCR parameters: basal respiration, maximal respiration, ATP production, and spare capacity in RGCs under elevated HP conditions (*n* = 6 to 8 biological replicates per group). (M) Slice through the middle of an EM tomography volume of an EV-NP mitochondrion. Three crista junctions are seen (arrowheads), with 1 showing a larger width (a). Top view of the surface-rendered mitochondrial volume after membrane segmentation (b). Side view (c). All the cristae have a lamellar shape. Slice through the middle of an EM tomography volume of an EV-EHP mitochondrion (d). The crista junction (arrowhead) is wider than 2 of those shown in (a). Top view of the surface-rendered volume (e). The crista at the top shows a twist, further noted in the side view (f). Slice through the middle of an EM tomography volume of an AKAP1-EHP mitochondrion (g). The 3 crista junctions (arrowheads) shown are wider than 2 of those in (a), especially the one at bottom right. The matrix is denser (more condensed) than seen in (a and d), suggesting an increase in metabolism, which is also consistent with wider crista junctions. The top view of the surface-rendered volume shows that the cristae mostly fill the mitochondrial interior, heightening the crista density (h). Side view showing another perspective of the twisting of several cristae (i). (N) Measurements for mitochondrial profile length (*n* = 20 mitochondria per group), profile area (*n* = 20 mitochondria per group), number per unit area (*n* = 10 mitochondria per group), volume density (*n* = 10 mitochondria per group), as well as rate of ATP generation per mitochondrial volume (*n* = 10 mitochondria per group), crista junction width (*n* = 20 mitochondria per group), number of crista junction per crista (*n* = 10 mitochondria per group), and number of crista junction per crista surface area (*n* = 10 mitochondria per group). Bars represent SEM. Statistical analysis was performed using unpaired Student’s *t*-test or one-way ANOVA and Tukey’s multiple comparisons test. \**P* < 0.05, \*\**P* < 0.01, \*\*\**P* < 0.00,1 and \*\*\*\**P* < 0.0001. Scale bars, 20 μm (D) and 200 nm (M). NP, normal pressure; EHP, elevated hydrostatic pressure.

### Augmenting AKAP1 expression promotes mitochondrial respiration, fusion activity and ATP production in RGCs against elevated HP

To further delineate the protective role of AKAP1 under elevated HP, primary RGCs were transduced with AAV2-AKAP1-green fluorescent protein (GFP) for 3 days, showing GFP co-expression in TUJ1-positive RGCs (fig. S8A and B). RGCs received AAV2-Null or AAV2-AKAP1, then were exposed to normal pressure (NP, 15 mmHg) or elevated HP (30 mmHg) for 24 h (Fig. 4J). Under elevated HP, AKAP1-RGCs displayed increased His, AKAP1, and RBPMS protein compared to Null-RGCs (fig. S8C). MTT assays showed reduced viability and mitochondrial activity in Null-RGCs under HP, but both were preserved by AKAP1 overexpression (fig. S8D). LDH release, a cell damage marker, was elevated in Null-RGCs under HP and decreased with AKAP1 (fig. S8D). AKAP1-RGCs exhibited enhanced mitochondrial respiration, evidenced by higher basal, maximal, and ATP-linked oxygen consumption rate (OCR), under elevated HP, while spare capacity and NP conditions remained unchanged (Fig. 4K and L). Electron microscopy (EM) tomography revealed AKAP1-expressing RGCs under HP had larger, elongated mitochondria with increased profile length and area, reduced number, and enhanced cristae architecture, density, and ATP production, indicative of increased fusion and improved bioenergetics (Fig. 4Mb-i and N; videos S1-3). Collectively, these findings demonstrate that AKAP1 promotes mitochondrial fusion, function, and RGC resilience during HP stress.

### Restoring AKAP1 expression ameliorates AMPK activation, oxidative stress, and apoptotic cell death in the retina of glaucomatous D2 mice

Given the association of 5’ AMP-activated protein kinase (AMPK) and p38-MAPK activation with mitochondrial (*8, 30–32*) and oxidative stress (*33, 34*) in glaucomatous RGCs, we tested whether AKAP1 restoration could attenuate these stress pathways and apoptosis in D2-Null retinas. Phosphorylated AMPK (pAMPK) was significantly increased in RGCs of D2-Null compared to D2G-Null mice but was significantly reduced after AKAP1 administration (fig. S9A-C). Similarly, phosphorylated p38 (pp38) levels were elevated in D2-Null retinas and RGCs, but substantially decreased by AKAP1 administration across all retinal layers (fig. S9D-F). Markers of oxidative stress and apoptosis, including SOD2, BAX, and BIM, were upregulated and anti-apoptotic BcL-xL was downregulated in D2-Null retinas; AKAP1 reconstitution reversed these alterations (fig. S9G and H). These results indicate that AKAP1 counters mitochondrial and oxidative stress responses and suppresses apoptosis, thus protecting RGCs in glaucomatous degeneration.

### Augmenting AKAP1 expression promotes RGC survival by promoting mitochondrial respiration, fusion activity and ATP production against oxidative stress

Oxidative stress and mitochondrial dysfunction are central to glaucoma pathogenesis (*2, 35–37*). AKAP1 overexpression protects RGCs from apoptosis under oxidative injury *in vitro* (*2*). To further test this, primary RGCs were transduced with AAV2-Null or AAV2-AKAP1, then treated with PBS or paraquat (PQ, 50 μM) for 24 h (fig. S10A). *Akap1* mRNA was robustly upregulated in AKAP1-RGCs under both conditions (fig. S10B). While PQ reduced AKAP1 and RBPMS protein in Null-RGCs, AKAP1-RGCs maintained higher levels (fig. S10C). PQ decreased cell viability and increased cell damage in Null-RGCs, as assessed by MTT and LDH assays, but AKAP1 overexpression preserved viability and reduced damage (fig. S10D). PQ also suppressed mitochondrial respiration, including basal, maximal, ATP-linked, and spare capacity, whereas AKAP1-RGCs restored all parameters (fig. S10E and F). Mitochondrial superoxide, elevated by PQ in Null-RGCs, was markedly reduced with AKAP1 (fig. S10G and H). We further examined mitochondrial ultrastructure by EM after 24 h of PBS or PQ (fig. S11A and B). PQ induced mitochondrial fragmentation in Null-RGCs, reflected by reduced length and area (fig. S11C). AKAP1-RGCs maintained mitochondrial size and structure under PQ. AKAP1 also reduced mitochondrial number but increased cristae density and ATP production per volume, consistent with enhanced fusion and function, while volume density and crista junction size were unchanged (fig. S11C). These results indicate that AKAP1 mitigates oxidative injury in RGCs by decreasing mitochondrial superoxide, preserving respiration, preventing fragmentation, and maintaining mitochondrial integrity under oxidative stress.

### Restoring AKAP1 expression dephosphorylates CaMKII and synapsin in the retina of glaucomatous D2 mice

Elevated IOP perturbs calcium homeostasis in RGCs, overactivating calcium/calmodulin-dependent kinase II (CaMKII) and glutamate receptors, ultimately driving excitotoxicity and RGC death (*38*). CaMKII, central to synaptic plasticity, phosphorylates synaptic proteins such as synapsin at Thr 286 (*39*), and early glaucomatous changes are closely tied to synaptic dysfunction (*14, 15, 40*), potentially mediated by CaMKII dysregulation. As AKAP1 is implicated in mitochondrial stabilization and dendrite modulation (*17, 18*), we examined whether restoring AKAP1 influences CaMKII phosphorylation in D2 retinas. Phosphorylated CaMKII (pCaMKII) was significantly increased in D2-Null versus D2G-Null but normalized following AKAP1 rescue (Fig. 5A-C). In human glaucomatous retinas, pCaMKII was elevated but not statistically significant (fig. S12A and B). AKAP1 deficiency had no significant effect on overall pCaMKII but did decrease pCaMKII specifically in the IPL (fig. S12C-E). Examining CaMKII’s downstream target synapsin (*39*), we found that pSynapsin was increased in D2-Null and restored to baseline in D2-AKAP1 retinas (Fig. 5D-F). pSynapsin intensity was reduced in human glaucomatous versus control retinas (fig. S12F and G). While AKAP1 loss did not affect whole-retina pSynapsin, it significantly increased pSynapsin in the IPL of *Akap1⁻^/^⁻* retinas (fig. S12H-J). Synapsin protein level was unchanged in *Akap1⁻^/^⁻*retinas compared to WT; however, synapsin intensity in the IPL was reduced in the glaucomatous human retina versus control (fig. S13A-E). Notably, D2-AKAP1 retinas had lower retinal synapsin levels than controls, although group differences in synapsin intensity were not observed (fig. S13F-H). These findings indicate that AKAP1 regulates synaptic function in glaucoma by modulating CaMKII and synapsin phosphorylation.

**Fig. 5.**
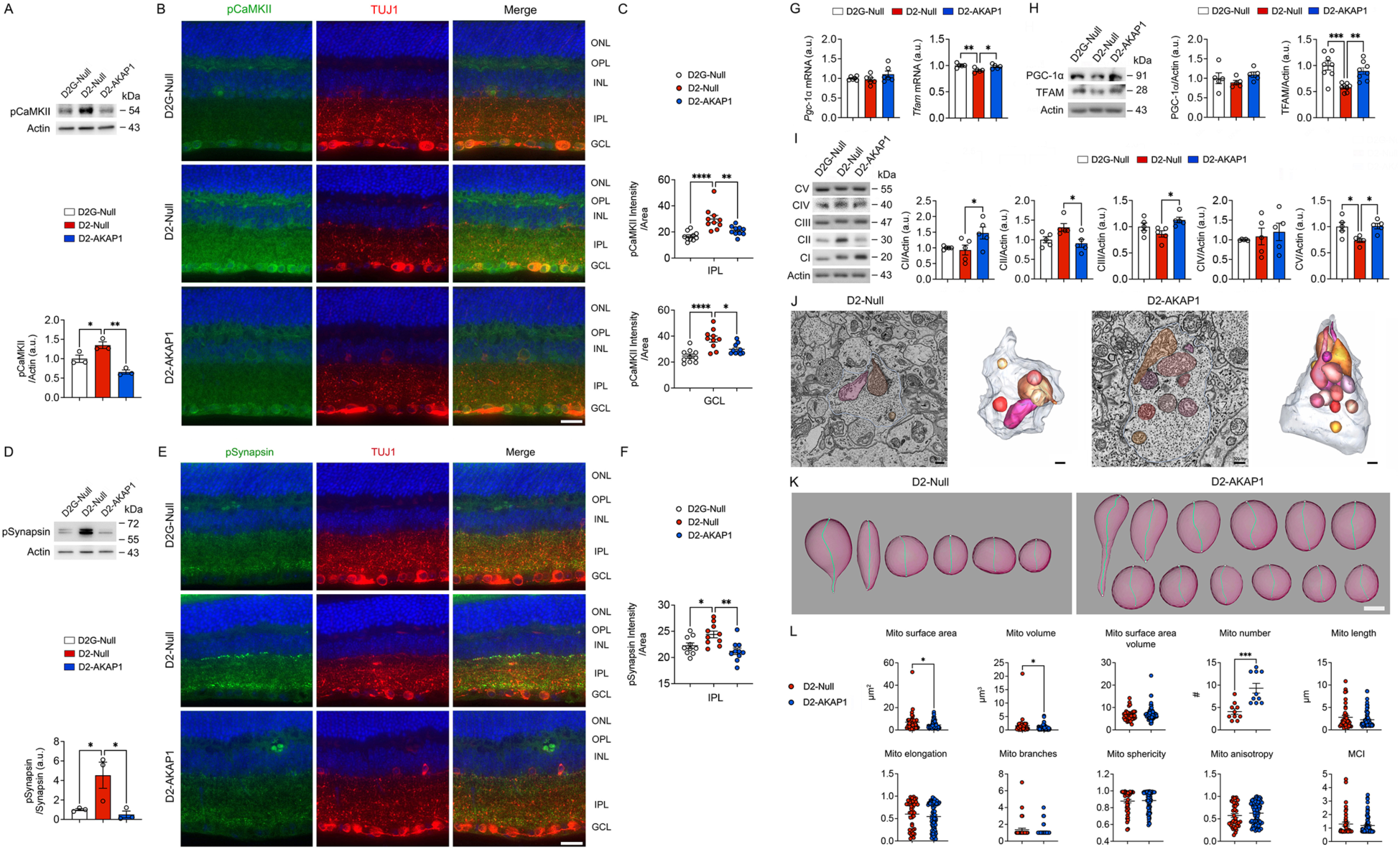
Restoring AKAP1 expression preserves synaptic activity and enhances mitochondrial biogenesis in the presynaptic terminals of glaucomatous D2 mice. (A) Representative image of a western blot and densitometry graph for pCaMKII expression in the retinas of D2G-Null, D2-Null, and D2-AKAP1 mice (*n* = 3 retinas per group). (B) Representative retinal images for pCaMKII (green) and TUJ1 (red) immunoreactivities. (C) Quantitative fluorescent intensity of pCaMKII immunoreactivity in the IPL and GCL of the retinas (*n* = 10 retina sections from 3 mice per group). (D) Representative image of a western blot and densitometry graph for pSynapsin expression in the retinas (*n* = 3 retinas per group). (E) Representative retinal images for pSynapsin (green) and TUJ1 (red) immunoreactivities. (F) Quantitative fluorescent intensity of pSynapsin immunoreactivity in the IPL of the retinas (*n* = 10 retina sections from 3 mice per group). (G) Quantitative PCR for *Pgc-1α* and *Tfam* gene expression in the retinas (*n* = 5 to 6 retinas per group) (H) Representative image of a western blot and densitometry graph for PGC-1α and TFAM in the retinas (*n* = 5 to 8 retinas per group). (I) Representative image of a western blot and densitometry graph for OXPHOS complex in the retinas of D2G-Null, D2-Null, and D2-AKAP1 mice (*n* = 5 retinas per group). (J) Representative SBEM volume showing typical synaptic button structures in D2-Null and D2-AKAP1 retinas; mitochondria (several different colors) highlighted. (K) Expedited and accurate segmentation and analysis of mitochondria. Surface rendering showing round forms of mitochondria. (L) Measurements for mitochondrial surface area (*n* = 20 mitochondria per group), volume (*n* = 20 mitochondria per group), surface area volume (*n* = 20 mitochondria per group), number (*n* = 10 mitochondria per group), length (*n* = 20 mitochondria per group), elongation (*n* = 20 mitochondria per group), branches (*n* = 10 mitochondria per group), sphericity (*n* = 20 mitochondria per group), anisotropy (*n* = 20 mitochondria per group), and MCI (*n* = 20 mitochondria per group) in D2-Null and D2-AKAP1 buttons. Bars represent SEM. Statistical analysis was performed using unpaired Student’s *t*-test or one-way ANOVA and Tukey’s multiple comparisons test. \**P* < 0.05, \*\**P* < 0.01, \*\*\**P* < 0.001, and \*\*\*\**P* < 0.0001. Scale bars, 20 μm (B and E) and 500 nm (J and K).

### Restoring AKAP1 expression triggers mitochondrial biogenesis in presynaptic terminals in the retina of glaucomatous D2 mice

Since AKAP1 modulates mitochondrial biogenesis (*8, 41*), we assessed whether its restoration enhances this process in glaucomatous D2 retinas. D2-AKAP1 retinas exhibited significantly elevated mitochondrial transcription factor A (*Tfam*) mRNA and protein levels versus D2-Null, while peroxisome proliferator-activated receptor gamma coactivator 1-alpha (*Pgc1-α*) expression remained unchanged (Fig. 5G and H). Notably, AKAP1 administration improved OXPHOS, increasing complexes I, III, and V, and preserving complexes II and IV, indicative of enhanced mitochondrial respiration (Fig. 5I). Given that synaptic deficits in the IPL often precede RGC loss in glaucoma (*2, 12–16, 42, 43*), we utilized serial block-face scanning electron microscopy (SBEM) (*19, 44*), for three-dimensional (3D) reconstruction of subcellular organelles (*45*) to assess mitochondrial morphology within presynaptic terminals. Figure 4J displays representative SBEM images (left) and corresponding 3D reconstructions (right) of presynaptic terminals in the IPL from D2-Null and D2-AKAP1 retinas, enabling precise localization of subcellular structures, particularly mitochondria (red) (videos S4 and 5). In Figure 5K, individual mitochondria were rendered in 3D and skeletonized (cyan lines) to quantify structural parameters, including length and branch shape. Both D2-Null and D2-AKAP1 terminals predominantly showed fragmented, rounded mitochondria (Fig. 5J and K; videos S4 and 5). Nevertheless, mitochondrial number per terminal was significantly increased in D2-AKAP1 compared to D2-Null retinas (Fig. 5L), indicating enhanced biogenesis. Despite this increase, mitochondrial surface area, length, elongation, branching, sphericity, anisotropy, and mitochondrial complexity index (MCI) (*46, 47*) remained comparable between groups (Fig. 5L). Together, these data show that AKAP1 restoration augments mitochondrial biogenesis without altering individual mitochondrial morphology, supporting synaptic integrity in the IPL during glaucomatous retinal degeneration.

### Restoring AKAP1 expression preserved synaptic activity by promoting synaptophysin but diminishing C1q expression in the retina of glaucomatous D2 mice

To evaluate synaptic vesicle integrity in the glaucomatous retina, we assessed synaptophysin, a synaptic vesicle marker. Synaptophysin protein and immunoreactivity were reduced in D2-Null retinas compared to D2G-Null, but restored with AKAP1 expression (Fig. 6A-C), with IPL immunoreactivity following the same trend. Human glaucomatous retinas also showed diminished synaptophysin compared to controls (fig. S14A and B), and AKAP1-deficient mice exhibited decreased synaptophysin versus wild-type (fig. S14C-E). We next examined C1q, a complement protein marking dysfunctional synapses for microglial clearance. C1q immunoreactivity was elevated in the IPL and GCL of D2-Null retinas but significantly decreased in D2-AKAP1 retinas (Fig. 6D and E). IBA1, a microglial marker, was similarly increased in D2-Null IPL, but reduced with AKAP1 restoration (fig. S15A and B). Given that C1q is localized in retinal glial cells (*12, 48*), elevated C1q co-localized with IBA1-positive microglia and glial fibrillary acidic protein (GFAP)-positive Müller glia in D2-Null IPL and GCL (fig. S15C and D). In human glaucoma, C1q was not increased overall, but localized with GFAP-positive Müller glia and astrocytes in the GCL (fig. S16A and B). *Akap1⁻^/^⁻* retinas showed increased C1q and GFAP in the IPL and GCL, co-localizing with GFAP-positive Müller glia (fig. S16C and D). Given the synapse loss observed in glaucoma(*49*), AKAP1 delivery restored postsynaptic density protein 95 (PSD95) expression in D2 mice (fig. S17A-C), though no significant differences were seen in human or *Akap1⁻^/^⁻* retinas (fig. S17D-H). Collectively, these findings indicate AKAP1 deficiency promotes retinal synaptic dysfunction, while AKAP1 protects against synaptic loss in glaucomatous neurodegeneration.

**Fig. 6.**
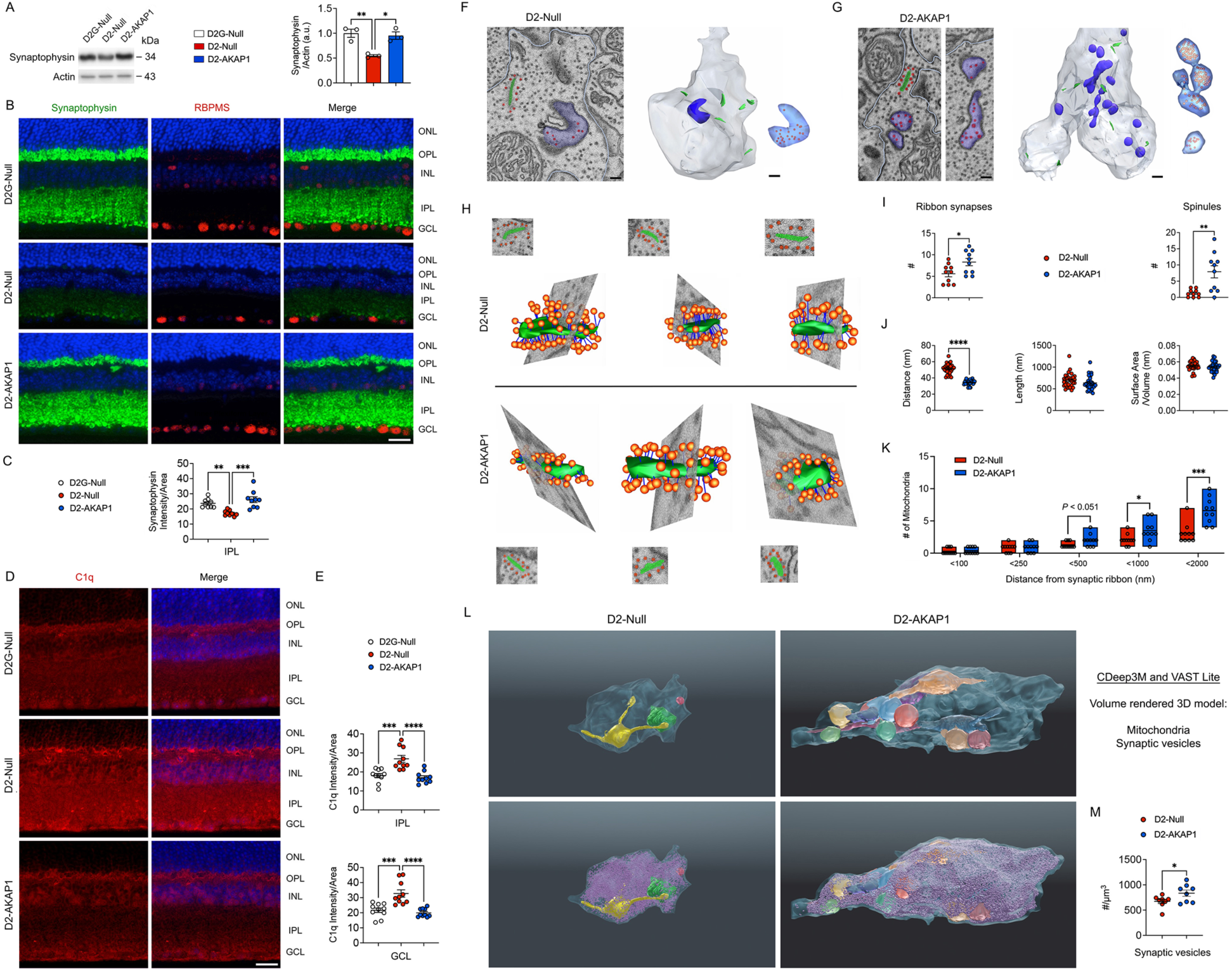
Restoring AKAP1 expression preserved synaptic activity by increasing synaptophysin but decreasing C1q expression in the retina of glaucomatous D2 mice. (A) Representative image of a western blot and densitometry graph for synaptophysin expression in the retinas of D2G-Null, D2-Null, and D2-AKAP1 mice (*n* = 3 retinas per group). (B) Representative retinal images for synaptophysin (green) and RBPMS (red) immunoreactivities. (C) Quantitative fluorescent intensity of synaptophysin immunoreactivity in the IPL of the retinas (*n* = 8 retina sections from 3 mice per group). (D) Representative retinal images for C1q (green) immunoreactivities. (E) Quantitative fluorescent intensity of C1q immunoreactivity in the IPL and GCL of the retinas (*n* = 10 retina sections from 3 mice per group). (F and G) Representative SBEM volume showing typical synaptic bouton structures in D2-Null and D2-AKAP1 retinas; ribbon synapses (green), synaptic vesicles (red), and spinules (blue) highlighted. (H) Representative images for SBEM and 3D IMOD analysis showing typical synaptic ribbons and vesicles in D2-Null and D2-AKAP1 retinas; synaptic ribbons (green) and synaptic vesicles (red) highlighted. (I) Measurements for the number of ribbon synapses and spinules (*n* = 10 synaptic boutons per group). (J) Measurements for the distance between ribbon synapse and vesicle, synaptic ribbon length, and synaptic ribbon surface area per volume (*n* = 30 synaptic ribbons per group). (K) Measurements for the number of mitochondria near ribbon synapses within a defined distance (*n* = 10 synaptic boutons per group). (L) Representative images for volume-rendered 3D models of mitochondria and synaptic vesicles using CDeep3M and Vast Lite. (M) Measurements for the number of synaptic vesicles using CDeep3M and Vast Lite (*n* = 8 synaptic boutons per group). Bars represent SEM. Statistical analysis was performed using unpaired Student’s *t*-test or one-way ANOVA and Tukey’s multiple comparisons test. \**P* < 0.05, \*\**P* < 0.01, \*\*\**P* < 0.001, and \*\*\*\**P* < 0.0001. Scale bars, 20 μm (B and D) and 500 nm (F and G).

### Restoring AKAP1 expression preserved synaptic structure and function in the retina of glaucomatous D2 mice

Using SBEM and 3D reconstruction, we found that AKAP1 administration significantly increased the number of ribbon synapses and spinules in the IPL (Fig. 6F-I; fig. S18). Imaging revealed representative SBEM planes (left) and corresponding 3D reconstructions (right) of ribbon synapses (*50–52*), comprising the synaptic ribbon (green) and rapidly releasing vesicle pool (red), as well as spinules (blue) within synaptic terminals of D2-Null and D2-AKAP1 retinas (Fig. 6F, G, and H; videos S6-9). AKAP1 restoration markedly elevated ribbon synapse and spinule numbers (Fig. 6I). In D2-Null terminals, six ribbon synapses and a single elongated spinule were observed, the latter lacking deep invasion into the presynaptic terminal. In contrast, D2-AKAP1 terminals exhibited nine ribbon synapses and multiple bulbous or varicose spinules, many deeply penetrating the presynaptic cytoplasm and filled with vesicles. This structural remodeling suggests enhanced synaptic connectivity and plasticity in D2-AKAP1 retinas, consistent with the role of AKAP1 in promoting synaptic resilience under glaucomatous stress.

Recent studies highlight the critical impact of spatial proximity between the ribbon and its rapidly releasing vesicle pool on synaptic transmission and neurodegeneration (*53–55*). Using algorithmic analysis, we quantified this distance within presynaptic terminals of D2-AKAP1 versus D2-Null retinas (Fig. 6F-I; fig, S18). We found a significantly decreased ribbon-to-vesicle pool distance in D2-AKAP1 retinas compared to D2-Null (Fig. 6H and K; fig. S18), while ribbon length and surface area per volume remained unchanged between groups (Fig. 6J; fig. S18). Since synaptic mitochondria are pivotal for energy supply and calcium regulation, and local mitochondrial dysfunction is linked to synaptic damage in neurodegeneration (*56–58*). We also measured the proximity of ribbon synapses to neighboring mitochondria. D2-AKAP1 retinas displayed a significant increase in mitochondria located near ribbon synapses within presynaptic terminals (Fig. 6K; fig. S18). These findings suggest AKAP1 restoration enhances synaptic architecture and energetics by optimizing spatial relationships among key synaptic components in glaucoma.

The total number of synaptic vesicles is a key determinant of synaptic transmission capacity, reflecting the synapse’s storage and rapid release ability for neurotransmitters. Accurate quantification is critical but challenging due to the small size and labor-intensive manual segmentation. To address this, we utilized SBEM combined with the machine-learning platform CDeep3M (*19, 59–61*) (fig. S19) to quantify presynaptic terminal volume, mitochondria, and vesicle number. This approach enabled efficient, high-throughput, and unbiased analysis. Notably, D2-AKAP1 retinas exhibited a significantly greater total vesicle count per terminal than D2-Null retinas (Fig. 6L and M; fig. S19; videos S10 and 11). Together, our data show that AKAP1 safeguards synaptic structure and function in the retina during glaucomatous stress. By increasing ribbon synapse and spinule numbers, minimizing ribbon–vesicle pool distance, and promoting mitochondrial proximity, AKAP1 enhances synaptic connectivity, neurotransmission, and energy balance. These protective effects underscore the therapeutic potential of AKAP1 for preventing synaptic loss and neurodegeneration in glaucoma.

### Restoring AKAP1 expression protects RGC axons and enhances mitochondrial fusion activity in the ONH of glaucomatous D2 mice

To further characterize AKAP1-mediated neuroprotection, we evaluated its impact on the glaucomatous ONH and astroglial activation. D2-Null mice exhibited marked ONH axon loss, evidenced by diminished neurofilament 68 (NF68) and increased GFAP immunoreactivity, indicating astrocyte activation, compared to D2G-Null (Fig. 7A). In contrast, D2-AKAP1 mice showed significant restoration of both NF68 and GFAP levels to those observed in D2G-Null (Fig. 7A and B). CTB labeling further confirmed preservation of ON axons in D2-AKAP1 versus D2-Null mice (Fig. 7C and D).

**Fig. 7.**
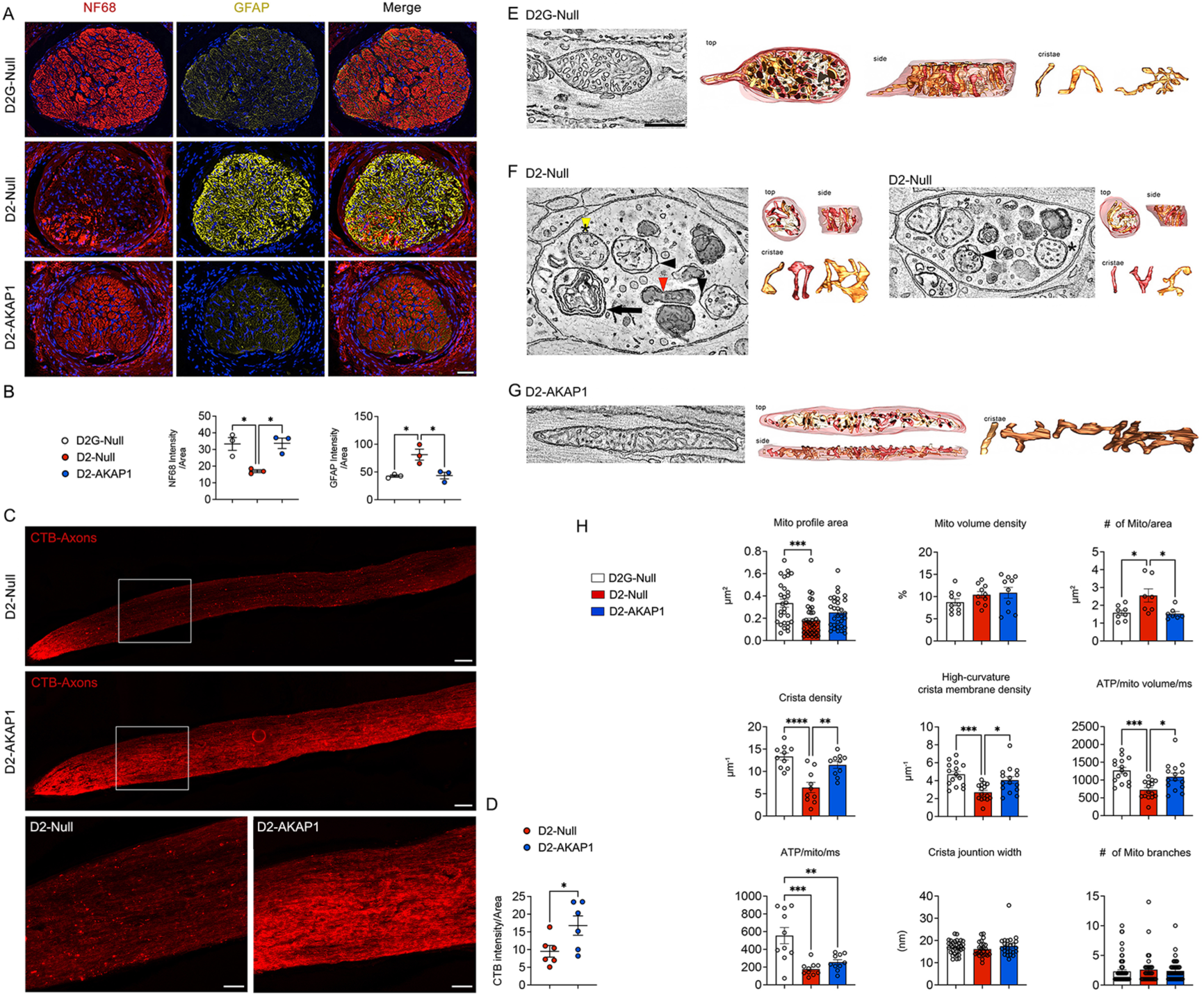
Restoring AKAP1 expression protects ONH axons by promoting mitochondrial fusion activity in the glaucomatous D2 mice. (A) Representative images for NF68 (red) and GFAP (yellow) immunoreactivities in the ONH of D2G-Null, D2-Null, and D2-AKAP1 mice. (B) Quantitative fluorescent intensity of NF68 and GFAP immunoreactivities (*n* = 3 ONH cross sections from 3 mice per group). (C) Representative images for CTB labeling (red) in the ONs. (D) Quantitative fluorescent intensity of CTB labeling in the ONs (*n* = 6 ON longitudinal sections from 6 mice per group). (E) Slice through the middle of an EM tomography volume of a D2G-Null mitochondrion that often shows large and mostly tubular cristae, although some lamellar cristae were also present. These mitochondria typically had high crista density. Top and side views of the surface-rendered mitochondrial volume after membrane segmentation. All the cristae have a lamellar shape. (F) Slice through the middle of an EM tomography volume of D2-Null mitochondria that show various forms of structural perturbations. In axonal varicosities, mitochondria and other organelles were aggregated. Normal-looking mitochondria were present, but were smaller than those found in the D2G-Null ONH and had reduced crista density (yellow arrowhead). Some mitochondria showed partially degraded cristae and had portions of their outer membrane separated from their inner boundary membrane (black arrowheads). Some mitochondria had a dark matrix (red arrowhead). The one below it is possibly another such mitochondrion. An autophagosome was also present (black arrow). This might be a mitophagosome, but the possible internalized mitochondrion has such deformed membranes that it is hard to determine this. Another varicosity showing 3 normal-looking mitochondria (numbered) that were typically small with reduced crista density and one mitochondrion with an abnormal bounding membrane (arrowhead). Other dark organelles had also accumulated in this varicosity. Top and side views of the surface-rendered mitochondrial volume after membrane segmentation. All the cristae have a lamellar shape. (G) D2-AKAP1 nearly completely rescued mitochondria from the damage prevalent in D2-Null ONH. The size of mitochondria was nearly as large as the Gpnmb mitochondria (not statistically different), and the crista density was nearly as great (again, not statistically different). Importantly, the structural damage to mitochondria was not seen, nor were there axonal varicosities with aggregated organelles in the D2-AKAP1 ONH. Top and side views of the surface-rendered mitochondrial volume after membrane segmentation. All the cristae have a lamellar shape. (H) Measurements for mitochondrial profile area (*n* = 20 mitochondria per group), volume density (*n* = 20 mitochondria per group), number per area (*n* = 10 mitochondria per group), crista density (*n* = 10 mitochondria per group), and high curvature crista membrane density, as well as rate of ATP generation per mitochondrial volume (*n* = 10 mitochondria per group), rate of ATP generation per mitochondrion (*n* = 20 mitochondria per group), crista junction width (*n* = 10 mitochondria per group), and number of mitochondrial branches (*n* = 10 mitochondria per group). Bars represent SEM. Statistical analysis was performed using unpaired Student’s *t*-test or one-way ANOVA and Tukey’s multiple comparisons test. \**P* < 0.05, \*\**P* < 0.01, \*\*\**P* < 0.001, and \*\*\*\**P* < 0.0001. Scale bars, 50 μm (A), 100 μm (C, upper/middle panels), 50 μm (C, bottom panel), and 200 nm (E-G).

We next assessed mitochondrial dynamics, cristae architecture, and ATP production in ONH axons via EM tomography (Fig. 7E-G; fig. S20; videos S12-15). Quantitative analysis revealed group differences in mitochondrial profile area, number, cristae density, high-curvature crista membrane density, and ATP production, while crista junction width and mitochondrial branching were unchanged (Fig. 7H). Axonal mitochondria in D2-Null mice displayed reduced profile area compared to D2G-Null, though D2-Null and D2-AKAP1 were similar, indicating decreased mitochondrial size in glaucoma. Mitochondrial volume density was unchanged. Notably, D2-Null mice had increased numbers of fragmented axonal mitochondria, which were significantly reduced in D2-AKAP1 mice, supporting the role of AKAP1 in promoting mitochondrial fusion in glaucomatous ONH axons. Importantly, cristae and high-curvature membrane densities were diminished in D2-Null but restored by AKAP1, suggesting improved mitochondrial structure and function (Fig. 7H).

Using EM tomography to model ATP production per unit mitochondrial volume via high-curvature crista membrane assessment(*62, 63*) (Fig. 7H), we found D2-Null mice exhibited a lower rate of ATP generation in axonal mitochondria compared to D2G-Null. D2-AKAP1 mice showed significant restoration of ATP production in axonal mitochondria over D2-Null (Fig. 7H). Notably, ONH axonal mitochondria displayed predominantly tubular cristae, which featured more high-curvature membrane than synaptic or Müller endfeet mitochondria, supporting greater ATP output(*34*) (Fig. 7H). The ATP generation rate per mitochondrion followed D2-Null < D2-AKAP1 < D2G-Null, driven by D2-Null’s smaller size and reduced high-curvature crista membrane; D2-AKAP1 mitochondria were only slightly smaller and had modestly less high-curvature membrane than D2G-Null, though these differences were not individually significant. The combined effect yielded a significant difference overall. Finally, crista junction width and mitochondrial branching did not differ statistically among groups (Fig. 7H).

### Restoring AKAP1 expression protects RGCs and their axons and preserves visual function in a mouse model of traumatic optic neuropathy

To further evaluate the neuroprotective effects of AKAP1, we used a mouse model of traumatic optic neuropathy (ONC), mimicking RGC axon injury and subsequent soma degeneration (*64*). Three-month-old C57BL/6J mice received intravitreal AAV2-Null or AAV2-AKAP1 injections three weeks prior to ONC (Fig. 8A; fig. S21), followed by assessment of RGC survival and visual function one week later. ONC led to marked reductions in RBPMS and AKAP1 protein levels in Null-ONC retinas versus controls, whereas AKAP1-ONC retinas showed restoration of both proteins (Fig. 8B). RGC counts confirmed a significant loss in Null-ONC, but this was rescued in AKAP1-ONC mice (Fig. 8C and D). Visual function measured via optomotor response, pERG, and pVEP showed Null-ONC mice had impaired spatial frequency, reduced pERG/pVEP amplitudes, and prolonged pVEP latency, while AKAP1 administration restored all metrics, supporting improved RGC function, ON conduction, and visual behavior (Fig. 8E-G). To assess axonal regeneration, CTB labeling and visual function were evaluated three weeks post-ONC (Fig. 8H). CTB intensity in ON and SC was significantly higher in AKAP1-ONC compared to Null-ONC mice, indicating enhanced axonal regrowth (Fig. 8I-L). Functional recovery was further supported by increased spatial frequency, pERG and pVEP amplitudes, and reduced pVEP latency in AKAP1-ONC mice (Fig. 8M-P). Collectively, these results demonstrate that AKAP1 promotes RGC survival, facilitates axonal regeneration, and preserves visual function after ON injury.

**Fig. 8.**
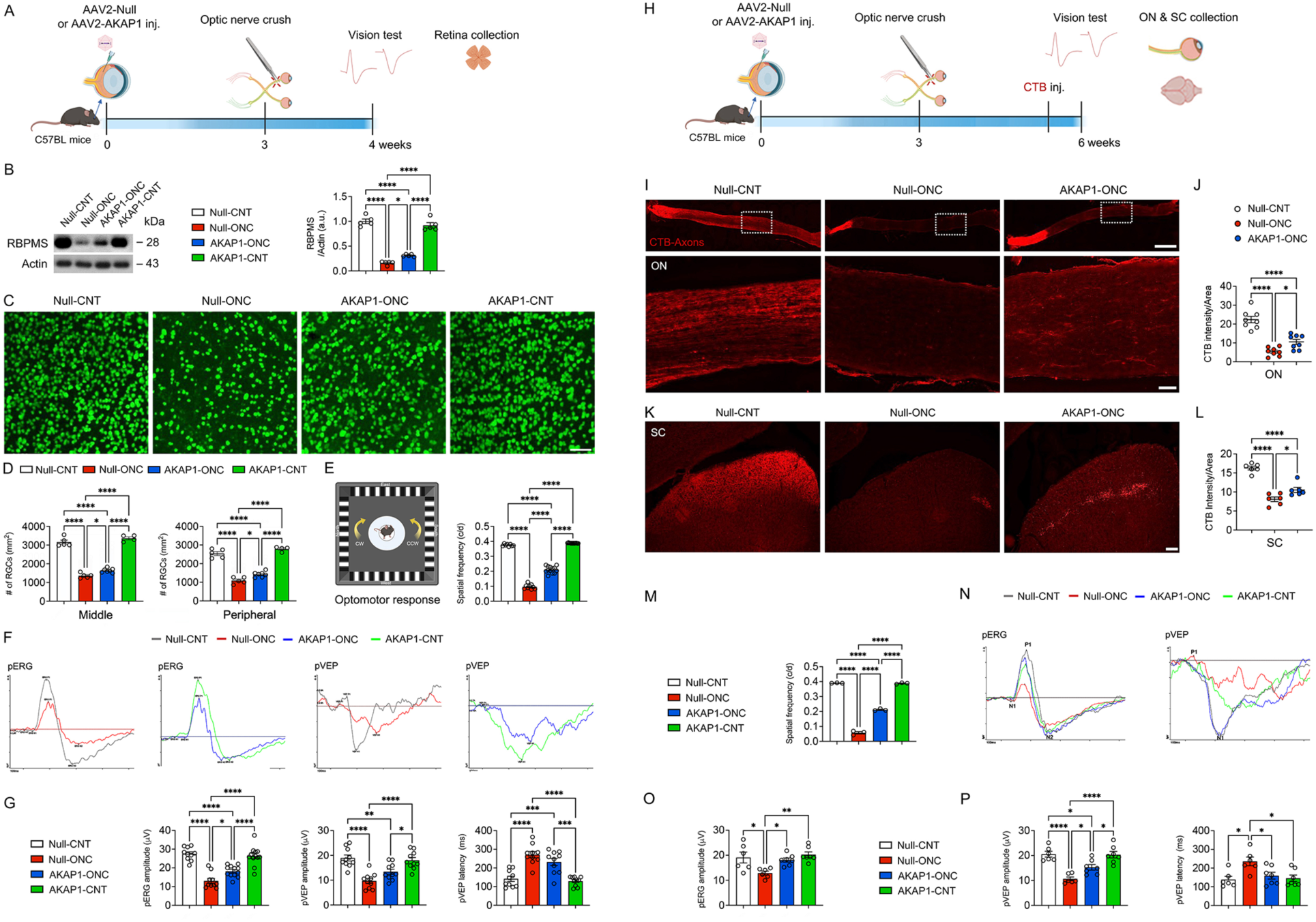
Restoring AKAP1 expression protects RGCs and their axons and preserves visual function, as well as enhances axon regeneration in a mouse model of traumatic optic neuropathy. (A) Experimental schematic and timeline of AAV injection, ONC, tissue collection, and data analysis in traumatic optic neuropathy (1-week ONC, *n* = 9-13 mice per group). (B) Representative image of a western blot and densitometry graph for RBPMS in the retinas (*n* = 5 retinas per group). (C) Representative whole-mounted retina images for RBPMS (green) immunoreactivity in the middle area. (D) Quantitative analysis of RGC number in the middle and peripheral areas of the retina (*n* = 4 to 7 retina wholemounts per group). (E) Representative recording graphs for optomotor response (*n* = 8 to 13 mice per group). (F) Representative recording graphs for pERG and pVEP measurements. (G) Quantitative analysis of pERG amplitude, pVEP amplitude, and pVEP latency (*n* = 10 to 12 mice per group). Note that AKAP1 expression restores visual function in ONC mice. (H) Experimental schematic and timeline of AAV injection, ONC, CTB labeling, tissue collection, and data analysis in traumatic optic neuropathy (3-week ONC, *n* = 6 to 7 mice per group). (I) Representative images for CTB labeling (red) in the ONs. (J) Quantitative fluorescent intensity of CTB labeling in the ONs (*n* = 8 ON longitudinal sections from 4-6 mice per group). (K) Representative images for CTB labeling (red) in the SCs. (L) Quantitative fluorescent intensity of CTB labeling in the SCs (*n* = 6 SC cross sections from 3-5 mice per group). Note that AKAP1 expression enhances ON axon regeneration in ONC mice. (M) Representative recording graphs for optomotor response (*n* = 3 mice per group). (N) Representative recording graphs for pERG and pVEP measurements. (O) Quantitative analysis of pERG amplitude (*n* = 5 mice per group). (P) Quantitative analysis of pVEP amplitude and pVEP latency (*n* = 6 to 7 mice per group). Note that AKAP1 expression restores visual function in ONC mice. Error bars represent SEM. Statistical analysis was performed using one-way ANOVA and Tukey’s multiple comparisons test. \**P* < 0.05, \*\**P* < 0.01, \*\*\**P* < 0.001, and \*\*\*\**P* < 0.0001. Scale bars, 50 μm (C), 500 μm (I, upper panel), 50 μm (I, lower panel), and 100 μm (K).

### Restoring AKAP1 expression results in higher resting calcium levels and promotes calcium buffering capacity against axonal injury

To assess the influence of AKAP1 on calcium signaling and survival following axonal injury, we established an *in vitro* laser ablation model replicating ONC damage. RGC axons were ablated 7.5 μm from the soma, and calcium dynamics were tracked pre– and post-injury (fig. S22A). Phase contrast images confirmed axonal thinning, while fluorescence imaging detected a rapid calcium influx after ablation (fig. S22B and C). Baseline calcium levels were higher in AKAP1-RGCs compared to Null-RGCs (fig. S22D). In AKAP1-RGCs, elevated pre-injury calcium was associated with lower post-injury peak, slower rises and falls, and reduced total calcium displacement, indicating an attenuated injury response. This relationship was absent in Null-RGCs (fig. S23). Although normalized calcium responses showed no statistical differences between groups (fig. S22E), AKAP1-RGCs exhibited less calcium spread to neighboring cells, supporting the modulatory effect of AKAP1 (fig. S22E). Calcium dynamics in axons themselves were comparable (fig. S22F). RGC viability was reassessed 24 h following ablation; AKAP1-RGCs demonstrated significantly higher survival, whereas Null-RGCs frequently fragmented (fig. S22G-H). Pre-injury measurements showed AKAP1-RGCs had wider axons, suggesting enhanced structural reinforcement (fig. S22G-H). These findings highlight the role of AKAP1 in dampening calcium injury responses and promoting RGC survival after axonal damage.

## DISCUSSION

Glaucoma treatment remains primarily focused on lowering IOP through medical therapy, laser procedures, surgery, or a combination of these. However, neuroprotection of RGCs remains a significant challenge due to complex, multifactorial mechanisms underlying RGC death, difficulties in early diagnosis, the absence of clinically proven neuroprotective agents, and obstacles to drug delivery. Gene therapy offers significant potential to overcome several of these challenges by enabling targeted, sustained, and long-term expression of protective proteins. This approach holds promise not only for preserving RGCs and their axons but also for preventing further vision loss in glaucomatous neurodegeneration.

Mitochondrial dysfunction is a central feature of the degenerative RGCs and axonal degeneration in glaucomatous neurodegeneration (*2, 3, 6, 44, 65*). Dysfunctional mitochondria exacerbate oxidative stress, impair ATP production, and trigger cell death pathways, thereby creating a hostile environment for maintaining healthy RGC compartments, including synapses, dendrites, somas, and axons (*2, 6, 44, 65*). AKAP1 modulates key signaling pathways critical for mitochondrial dynamics, bioenergetics, and cell survival (*2, 8–10, 66, 67*). Specifically, AKAP1 regulates mitochondrial fusion and fission by recruiting PKA and controlling DRP1 phosphorylation, thereby maintaining mitochondrial integrity and function(*8–11*). Dysregulation of AKAP1 is associated with impaired mitochondrial dynamics and neuronal dysfunction in the central nervous system, including the retina (*8–11, 68*). Consistent with this, our previous study demonstrated that *Akap1^−/−^*mice exhibit disrupted mitochondrial dynamics and significant RGC loss (*11*). Conversely, *in vitro* studies have shown that AKAP1 overexpression via AAV2-mediated transduction enhances mitochondrial function and protects primary RGCs from oxidative stress-induced apoptosis (*2*). Despite these insights, the *in vivo* protective effects and mechanisms underlying AKAP1 restoration in RGC mitochondria during glaucomatous neurodegeneration remain poorly understood. Here, we report a marked reduction of AKAP1 expression in RGCs from glaucomatous human eyes. Notably, reconstitution of AKAP1 expression promotes RGC survival and preserves visual function across two experimental glaucoma models and a traumatic optic neuropathy model, while also maintaining the integrity of the central visual pathway. These protective effects correlate with enhanced mitochondrial fusion and increased bioenergetic capacity in RGCs challenged by glaucomatous insults such as elevated IOP and oxidative stress. In contrast, loss of AKAP1 exacerbates RGC dysfunction and worsens visual outcomes in experimental glaucoma. Together, these findings underscore the pivotal role of AKAP1-mediated mitochondrial homeostasis in RGC survival and vision preservation, positioning AKAP1 as a promising therapeutic target for glaucomatous neurodegeneration.

Reduced AKAP1 expression in the glaucomatous retina is unlikely to represent a mere epiphenomenon or a secondary consequence of RGC dysfunction. This conclusion is supported by experimental evidence demonstrating that restoration of AKAP1 expression, such as via AAV2-mediated gene therapy, results in marked improvements in RGC survival, mitochondrial function, synaptic integrity, and visual performance. The ability of elevated AKAP1 levels to rescue these deficits strongly implicates AKAP1 loss as a causative factor driving disease progression rather than merely a downstream biomarker of ongoing neurodegeneration. Accordingly, these findings support the interpretation that AKAP1 functions as a critical mechanistic driver of RGC resilience and synaptic health under glaucomatous stress, rather than acting as a bystander in the degenerative process.

A growing body of evidence indicates that synaptic and dendritic alterations occur early in the glaucoma progression, often preceding overt loss of RGC somas (*14–16, 22, 69*). Presynaptic terminals, which demand high metabolic activity, are particularly vulnerable to elevated IOP and oxidative stress. Mitochondrial dysfunction and increased reactive oxygen species production in glaucoma likely contribute to synaptic injury prior to RGC soma degeneration (*2, 14, 40, 70, 71*). However, whether impaired mitochondrial dynamics directly disrupt synaptic activity in the glaucomatous retina remains unclear. While reductions in AKAP1/PKA signaling are linked to decreased synaptic density and compromised neuronal function (*17, 18*), activation of this pathway facilitates synaptic plasticity and promotes neuronal growth (*17, 18*). Notably, PKA anchored by AKAP1 at the OMM plays a critical role in modulating synaptic activity and confers neuroprotection against glaucomatous insults, including excitotoxicity and oxidative stress (*31, 72–74*). Consistent with this, our findings demonstrate that AKAP1/PKA signaling is essential for maintaining mitochondrial dynamics, enhancing mitochondrial biogenesis, and improving mitochondrial respiration in the glaucomatous retina. Together, these data highlight a critical role for AKAP1/PKA-mediated mitochondrial regulation in preserving synaptic integrity and RGC function during glaucomatous neurodegeneration.

CaMKII is a pivotal regulator of synaptic function, critically modulating neurotransmitter release by phosphorylating key synaptic proteins (*75*). Among its substrates, CaMKII phosphorylates synapsins, thereby facilitating synaptic vesicle mobilization and enhancing neurotransmitter release (*39, 76, 77*). In glaucoma, stressors, such as elevated IOP, disrupt calcium homeostasis in retinal cells, including RGCs, leading to aberrant CaMKII activation and excessive phosphorylation. This hyperphosphorylation of CaMKII amplifies glutamatergic signaling via NMDA and AMPA receptors, leading to increased calcium influx and excitotoxic neuronal injury—a primary mechanism driving RGC degeneration. CaMKII is a multifunctional enzyme that becomes autonomously active upon phosphorylation at Thr286 in response to intracellular calcium elevations, thereby regulating synaptic plasticity by targeting multiple synaptic proteins. Notably, synaptic dysfunction arises early in glaucoma, preceding overt RGC loss, and is implicated in disrupted communication between RGCs and their synaptic targets (*49, 78, 79*). Overactivation or dysregulation of CaMKII phosphorylation may destabilize these synapses, thereby accelerating neurodegenerative processes (*38*). Furthermore, the presynaptic proteins synaptophysin and synapsin cooperatively maintain synaptic vesicle clustering through ionic interactions. CaMKII-mediated phosphorylation of synapsin disrupts its interaction with synaptophysin, impairing vesicle clustering and synaptic efficacy (*77*). Collectively, these mechanistic insights highlight CaMKII dysregulation as a central contributor to synaptic impairment and RGC loss in glaucomatous neurodegeneration.

CaMKII activity is typically associated with increased or preserved PSD95 expression (*80*). Loss of CaMKII activity correlates with RGC degeneration, whereas preservation or reactivation of CaMKII protects RGCs in glaucoma models (*38, 81*). Moreover, the reduction of PSD95 puncta correlates with the severity of dendritic remodeling during early glaucomatous neurodegeneration in experimental glaucoma (*49*), suggesting that synaptic pruning precedes dendritic atrophy and cell loss. In this study, AKAP1 administration reduced CaMKII activity while increasing PSD95 expression in the glaucomatous retina. These findings likely represent a maladaptive phase of synaptic remodeling in which dysregulated kinase activity simultaneously reflects compensatory and pathological processes, contributing to synaptic degeneration, impaired neurotransmission, and progressive visual dysfunction. This suggests that AKAP1 functions as a neuroprotective and synapse-stabilizing factor in the retina, particularly during early glaucomatous stress. Given that AKAP1-mediated mitochondrial stabilization enhances dendritic outgrowth but constrains synapse formation (*17, 18*), these effects may be mediated by altered calcium handling or energy distribution. Since mitochondrial dysfunction can influence intracellular calcium transients that regulate CaMKII activation, this provides a plausible mechanism by which AKAP1 may indirectly modulate synaptic signaling. However, the precise role of AKAP1 in synaptic function under glaucomatous conditions remains to be determined. Although current data do not support a direct interaction between AKAP1 and CaMKII or its closely associated synaptic signaling proteins, we propose that AKAP1 influences synaptic integrity indirectly through pathways affecting CaMKII and synapsin phosphorylation and PSD95 expression, thereby contributing to synaptic stability and function during disease progression (*38, 49, 81*).

AKAP1 administration was found to reduce the phosphorylation of synapsins, a family of synaptic vesicle-associated proteins that mediate vesicle tethering to the actin cytoskeleton, while preserving the expression of synaptophysin, a membrane protein integral to presynaptic vesicles (*77, 82, 83*). These molecular effects correlated with an increased number of ribbon synapses and synaptic vesicles, indicating enhanced presynaptic structural integrity. Given the well-established role of PKA-mediated synapsin phosphorylation in regulating synaptic vesicle mobilization and neurotransmitter release (*84–86*), our findings suggest that AKAP1/PKA deficiency may contribute to presynaptic dysfunction via these dysregulations in the glaucomatous retina. Furthermore, presynaptic terminals are particularly susceptible to metabolic stress, and the concurrent maintenance of synaptophysin expression alongside AKAP1-driven mitochondrial protection likely plays a critical role in mitigating synaptic deficits associated with glaucoma. Together, these data highlight the important contribution of AKAP1-mediated mitochondrial and synaptic regulation in preserving presynaptic function during glaucomatous neurodegeneration.

Intriguingly, our findings demonstrate that elevated IOP significantly upregulates complement component C1q in the glaucomatous D2 retina, whereas administration of AKAP1 restores C1q expression to baseline levels. Previous work by Stevens et al. revealed that C1q localizes to retinal synapses in D2 mice early during glaucoma, preceding or coinciding with synapse loss (*12*). In the glaucomatous retina, C1q serves as a molecular tag marking synapses for elimination even at early disease stages, contributing to compartment-specific synapse loss that may drive dendritic atrophy and axonal degeneration as the disease advances (*12*). Our results therefore suggest that AKAP1 mitigates synapse loss in glaucoma by inhibiting activation of the complement cascade, an effect accompanied by reduced microglial activation. Future investigations are warranted to elucidate the precise mechanisms underlying AKAP1-mediated regulation of complement signaling in the glaucomatous retina. Notably, prior studies have shown that AKAP1 overexpression promotes dendritic growth while modulating synaptic density in hippocampal neurons *in vitro* (*9*), further supporting a role for AKAP1 in synaptic maintenance. Collectively, our findings underscore that restoring AKAP1 expression protects synapses in the glaucomatous retina by enhancing mitochondrial function, attenuating stress-induced kinase activation, and preserving both presynaptic and postsynaptic protein integrity, thereby providing a multifaceted neuroprotective effect against RGC degeneration (fig. S24).

Elevated IOP-induced structural and functional impairments of mitochondria in RGC axons in the ONH accelerate glaucomatous neurodegeneration (*2*). Consequently, targeting mitochondrial dynamics and function represents a promising therapeutic strategy to protect RGC axons against glaucomatous insults such as elevated IOP and oxidative stress (*2*). Notably, AKAP1 administration mitigated axonal damage by promoting mitochondrial fusion and preserving bioenergetic capacity, as evidenced by maintained cristae density and ATP production within the glaucomatous glial lamina. Furthermore, AKAP1 conferred neuroprotection by enhancing RGC survival and improving visual function following ONC-induced axonal injury. In vitro studies further revealed that AKAP1 expression modulates intracellular calcium dynamics, augmenting RGC axonal resilience against laser ablation-induced damage. Intriguingly, AKAP1 amplification resulted in elevated basal calcium levels, which correlated with a more controlled calcium response and diminished injury-induced calcium fluctuations. Given the critical role of calcium homeostasis in neuronal survival and the central function of mitochondria in buffering intracellular calcium levels, these findings underscore the capacity of AKAP1 to regulate mitochondrial calcium handling.

Mechanistically, AKAP1 promotes mitochondrial elongation through inhibition of DRP1, thereby enhancing mitochondrial capacity to sequester and release calcium (*10, 67*). Merrill and colleagues demonstrated that AKAP1-mediated neuroprotection is associated with maintained mitochondrial membrane potential and prevention of delayed calcium deregulation during excitotoxic stress (*67*). These observations suggest that AKAP1 preserves mitochondrial function by preventing pathological calcium overload, a key contributor to neuronal injury. Importantly, mitochondria-mediated regulation of calcium signaling is critical for axonal integrity following injury. Collectively, these findings highlight the therapeutic potential of AKAP1 in preserving RGCs and their axons and restoring vision by stabilizing mitochondrial dynamics, maintaining calcium homeostasis, and enhancing mitochondrial calcium buffering capacity in response to axonal damage associated with glaucomatous neurodegeneration (fig. S24).

While our study establishes AKAP1 as a potent neuroprotective factor that preserves RGCs and restores visual function by promoting mitochondrial biogenesis in presynaptic terminals and enhancing mitochondrial fusion and bioenergetics in axons during glaucomatous neurodegeneration, several limitations warrant consideration. First, the precise molecular mechanisms by which AKAP1 regulates critical pathways, including CaMKII signaling, synapsin phosphorylation, and complement proteins such as C1q, remain incompletely defined and require further elucidation. Second, although AAV2-mediated AKAP1 gene therapy demonstrates therapeutic efficacy in murine and *in vitro* models, its translational relevance is constrained by the absence of validation in large animal models, such as minipigs or non-human primates, which better recapitulate human ocular anatomy and glaucoma pathophysiology. Third, the models employed, including DBA/2J, microbead-induced ocular hypertension, and ONC, may not fully capture the heterogeneity and chronic progression characteristic of human glaucoma, potentially limiting generalizability. Fourth, while functional measures of visual behavior and electrophysiology provide valuable endpoints, their correlation with underlying structural and molecular changes requires further strengthening. Fifth, the influence of AKAP1 on calcium homeostasis, though promising, needs more comprehensive evaluation across diverse retinal cellular compartments. Finally, the safety profile, specificity, and delivery efficiency of AAV-based gene therapies remain critical obstacles to clinical translation that must be carefully addressed through long-term studies. Together, these limitations highlight the need for deeper mechanistic investigations, validation in broader and more clinically relevant models, and extended assessments to rigorously evaluate AKAP1-based therapeutic strategies for glaucoma.

In summary, AKAP1 gene therapy preserves vision in glaucoma by enhancing mitochondrial fusion, reducing oxidative stress, stabilizing synapses, and protecting RGCs. This approach provides neuroprotection beyond IOP control through mitochondrial and synaptic regulation. Here, our study highlights that AKAP1 is markedly downregulated in glaucomatous human and mouse retinas, implicating its deficiency in RGC vulnerability. AKAP1 expression enhances mitochondrial fusion and energy production, preserving mitochondrial integrity and function in RGCs. AKAP1 promotes mitochondrial biogenesis and maintains synaptic integrity, protecting retinal synapses from glaucomatous neurodegeneration. AKAP1 facilitates axonal regeneration, preserves the visual pathway, and sustains visual function. These findings establish AKAP1 as a master regulator of mitochondrial and synaptic homeostasis and a promising therapeutic target for RGC protection, axonal regeneration, and vision preservation in glaucomatous neurodegeneration.

## MATERIALS AND METHODS

### Study design

This study aimed to investigate a new concept that *in vivo* delivery of AAV2-AKAP1 protects RGCs in mouse models of glaucoma and traumatic optic neuropathy, contributing to developing a new treatment for glaucoma. In the current study, the retinas from 6 normal donor eyes and 6 eyes from patients with POAG were analyzed to evaluate the expression of AKAP1, PKA, and synaptic markers. All the normal donor and glaucomatous patient eyes were obtained from San Diego Eye Bank with a protocol approved by the University of California, San Diego Human Research Protection Program; the subjects were not diagnosed with other eye diseases, diabetes, or a chronic central nervous system disease. To test the efficacy of intravitreal AAV using the AAV2 vector encoding the AKAP1 protein *in vivo*, we used two different mouse models of glaucoma, including the D2 strain and MB-induced ocular hypertension, which elevate IOP substantially, and an ONC as a model of traumatic optic neuropathy. To test the effect of IOP elevation on AKAP1 deficiency, we used the MB model in *Akap1^−/−^*mice intravitreally injected with AAV2-AKAP1. The AAV-Null, which does not express AKAP1, was used as a negative control. Age– and sex-matched littermates were used as a control in each experiment. No mice were excluded from the data analysis. The number of biological replicates for *in vivo* and *in vitro* experiments is indicated in each figure legend.

### Human tissue samples

Human retina tissue sections were obtained, via San Diego Eye Bank, from five eyes of donors without a history of glaucoma or any other eye disease, and from five eyes of glaucoma patients (*19*). The studies were approved by the University of California, San Diego Human Research Protection Program. Both control and glaucoma patients had no history of other eye disease, diabetes, or chronic central nervous system disease (Table S5).

### Animals

Adult male and female D2 and age-matched D2G mice (The Jackson Laboratory), C57BL/6J (The Jackson Laboratory), *and Akap1^−/−^* mice were housed in covered cages, fed a standard rodent diet ad libitum, and kept on a 12 h light/12 h dark cycle. C57BL/6J mice were used as WT mice. *Akap1^−/−^* mice on a C57BL/6J background were generated in the laboratory of Dr. Stefan Strack, as previously reported (*10, 11*). Animals were assigned randomly to experimental and control groups. To investigate the effect of AAV2-AKAP1 in experimental mouse models of glaucoma, 5-month-old D2 and age-matched D2G mice, and 3-month-old C57BL/6J mice were used. All animal procedures were in accordance with the Association for Research in Vision and Ophthalmology Statement for the Use of Animals in Ophthalmic Vision Research and under protocols approved by the Institutional Animal Care and Use Committee at the University of California, San Diego.

### IOP Measurement

IOP elevation typically begins between 5 and 7 months of age, and by 9-10 months of age, ON axon damage due to elevated IOP has advanced dramatically (*19, 44, 65*). The IOP was assessed as previously described, and 65.3% (64/98) of glaucomatous DBA/2J mice showed confirmed IOP rise at 10 months. The highest IOP rise was 21.5 ± 4.5 mmHg in the right eye of 10-month-old D2 mice and 19.9 ± 3.7 mmHg in the left eye. Significant ON damage, including axon loss, was found in glaucomatous D2 mice aged 10 months, indicating the presence of acquired optic neuropathy. Each of the 10-month-old D2 mice in this study had a single IOP measurement to demonstrate a spontaneous IOP increase of more than 20 mmHg, with five measurements obtained and the average calculated automatically with a tonometer (Icare TONOVET). Furthermore, non-glaucomatous control C57BL/6 or D2G mice used in this study (*n* = 5 measurements) also had a single IOP measurement. Mice were anesthetized with Isoflurane before IOP measurement.

### Mouse model of MB-induced ocular hypertension

MB-induced glaucoma was induced using a modified protocol based on previously reported (*19, 28, 29, 87*). In brief, six-week-old C57BL/6J mice were intravitreally injected with either AAV2-Null or AAV2-AKAP1 as described above. At 3 weeks post-injection, mice were anesthetized by an IP injection of ketamine/xylazine. We topically applied 0.5% proparacaine hydrochloride, and the pupils were dilated with 1:1 mixture of 1% tropicamide and 2.5% phenylephrine, and then the cornea was gently punctured using a 30-gauge needle. Next, 2 µl of 1-µm-diameter polystyrene MB suspension (containing 3.0 × 10^7^ beads) followed by an air bubble, 2 µl of 6-µm-diameter polystyrene MB suspension (containing 6.3 × 10^6^ beads, Polysciences), and 1 µl of PBS containing 30% Healon was injected into the anterior chamber using a 32-gauge needle (Hamilton Company) connected to a syringe (Hamilton Company), which induced moderate elevation of IOP for ≥ 8 weeks. An equivalent volume of PBS was injected into the contralateral eyes to serve as a sham control. IOP was measured weekly or biweekly for 8 week using tonometer. At 8 weeks after MB injection, mice were tested for visual function and then sacrificed, and the retinal tissues were isolated.

### Mouse model of ONC

A modified protocol based on a previous report (*64*) was used to conduct ONC-induced ON injury. Three-month-old C57BL/6J mice were intravitreally injected with either AAV2-Null or AAV2-AKAP1. At 3 weeks after AAV injection, the mice were anesthetized using a mixture of ketamine and xylazine, and 1% proparacaine drops were also administered to their eyes. A limbal conjunctival peritomy was performed in the inferior eye region to allow access to the ON, which was exposed through a small window made by gentle blunt dissection between the surrounding muscle bundles and fatty tissue. During the procedure, extra care was taken to avoid damage to the muscles or the vortex veins. The ON was clamped with a pair of Dumont no. 5 self-closing tweezers for 3 sec at a site approximately 2 mm posterior to the globe. The contralateral eyes were used as a control with a sham operation. Depending on experimental design, either after 1 week or 3 weeks of ONC, the mice were tested for optokinetic response and visual functions, and retinal tissues were prepared.

### Primary RGCs isolation and purification

RGCs from postnatal 3-5 days of Sprague-Dawley rats were purified by immunopanning and cultured in serum-free defined growth medium, as previously reported (*44, 88*). Purified RGCs were seeded on 60 mm dishes, 24-well plates and 96-well plates in approximately 2 × 10^5^, 1 × 10^5^, and 1.5 × 10^4^, respectively, which were coated first with poly-d-lysine (70 kDa, 10 μg/ml; Sigma-Aldrich) and then with laminin (10 μg/ml; Sigma-Aldrich) in neurobasal medium. RGCs were cultured in serum-free defined growth medium containing BDNF (50 μg/ml; Sigma-Aldrich), CNTF (10 μg/ml; Sigma-Aldrich), insulin (5 μg/ml; Sigma-Aldrich), and forskolin (10 μg/ml; Sigma).

### PQ and elevated HP treatment in primary RGCs *in vitro*

Cultured RGCs were subjected to either PQ or elevated HP treatment to model oxidative and mechanical stress (*31, 34, 44*). For oxidative stress, cells were treated with 50 μM PQ (Sigma-Aldrich) diluted in complete growth medium for 24 hours. PQ-induced oxidative damage by generating intracellular ROS was evaluated using MitoSOX red staining. For elevated HP treatment, RGCs were exposed to elevated pressure for 24 h using a custom-built pressurized incubator, as previously described. Briefly, culture plates were placed in a sealed plexiglass pressure chamber connected to a certified 5% CO₂/95% air gas source (Airgas Inc.) via a low-pressure two-stage regulator (Gilmont Instruments). The pressure within the chamber was maintained at 15 mmHg for normal and 30 mmHg above atmospheric pressure throughout the treatment. The setup was housed in a standard tissue culture incubator at 37 °C with 5% CO₂ to ensure optimal cell culture conditions during pressurization. Cells were treated under these stress conditions 3 days post-transduction with AAV2-AKAP1 or control (Null). Following treatment, cells were subjected to downstream analyses including MTT and LDH assays, mitochondrial ROS detection, Western blot analysis of mitochondrial and apoptotic markers, and mitochondrial respiration profiling using Seahorse XF extracellular flux analysis to measure OCR.

### Elevated HP treatment in rMC-1 cells *in vitro*

rMC-1 cells, an immortalized rat retinal Müller glial cell line (Kerafast), were grown in Dulbecco’s Modified Eagle Medium (DMEM, Corning Inc.) supplemented with 5% fetal bovine serum and 1% penicillin/streptomycin solution at 37 °C in a humidified CO2 incubator. The cells were seeded into six-well plates at a density of 2 × 10^5^ cells/well for Western blot analysis or 96-well plates at 1.5 × 10^4^ cells/well for MTT and LDH assays and maintained for 24 h. Cells were treated to 15 mmHg or 30 mmHg for 24 h. A cell culture experiment was performed with 3 independent experiments.

### Production of recombinant AAV2-AKAP1 vector

AAV2-AKAP1 was generated using a method, as previously reported (*2, 19*). Murine AKAP1 (1-857aa) was fused with 6x-His at the C-terminus (AKAP1-His). AKAP1-His was cloned into pAAV-MCS vector (AAV2-M4 for Rep/Cap) (Agilent Technologies). AAV was titrated using quantitative PCR (qPCR) with primers for the inverted terminal repeats (ITRs) (TaKaRa Bio Inc).

### Transduction with recombinant AAV2 constructs *in vivo* and *in vitro*

Mice were anesthetized with an intraperitoneal (IP) injection of a cocktail of ketamine (100 mg/kg, Ketaset; Fort Dodge Animal Health) and xylazine (9 mg/kg, TranquiVed; VEDCO Inc.), and topical 1% proparacaine eye drops. To begin the procedure, the globe was punctured using the bevel of a 30-gauge needle, and then a 33-gauge Hamilton syringe was inserted through the puncture site, and a total of 1 µl of either AAV2-Null or AAV2-AKAP1 (1 × 10^13^ gc/ml) was injected into the vitreous humor of the eye. Injections were given slowly for over 1 min, and the needle was maintained in position for an additional 5 min to minimize vector loss through the injection tract (*19, 44*). For the *in vitro* study, primary RGCs were transduced with either AAV2-Null or AAV2-AKAKP1 for 3 days.

### Tissue Preparation

At the end of each mouse experiment, mice were anesthetized with an IP injection of ketamine/xylazine, as described above, and cervical dislocation. Following enucleation of eyeballs, the eyes were either fixed with 4% paraformaldehyde in PBS for 4 h at 4 °C, and the retinas were prepared as flattened whole-mounts for immunohistochemical analysis, or the retinas were isolated immediately and used for Western blot analysis and qRT-PCR. For SCs, the mouse brain was fixed via cardiac perfusion with 0.9% saline followed by 4% paraformaldehyde (Sigma-Aldrich) in phosphate-buffered saline (PBS, pH 7.4, Sigma-Aldrich). Then the brain sample was processed for cryosection. For EM studies, the eyes were fixed via cardiac perfusion with a solution of 2% paraformaldehyde and 2.5% glutaraldehyde in 0.15 M sodium cacodylate (pH 7.4) at 37°C, followed by placing them in a pre-cooled fixative of the same composition on ice for 1 h.

### Western blot analysis

The harvested retinas were homogenized for 1 min on ice with a RIPA lysis buffer [25 mM Tris-HCl, pH 7.6, 150 mM NaCl, 1% NP-40, 0.5% deoxycholic acid, 0.1% sodium dodecyl sulfate (SDS)], (Thermo Scientific), containing protease inhibitor cocktail (Thermo Scientific). The lysates were then centrifuged at 13,000 rpm for 15 min, and protein concentration in the supernatants was measured by BCA Protein assay (Thermo Scientific). Proteins (10-20 μg) were run on a 4-20% Mini-PROTEAN TGX-precast protein gel electrophoresis (Bio-Rad) and transferred to polyvinylidene difluoride (PVDF) membranes (GE Healthcare Bioscience). The membranes were blocked with TBS/0.1% Tween-20 (T-TBS) containing 5% skim-milk for 1 hour at room temperature and incubated with primary antibodies (Table S6) for overnight at 4 °C. Membranes were washed three times with T-TBS and then incubated with horseradish peroxidase-conjugated secondary antibodies (Cell Signaling Technology) for 1 h at room temperature. They were developed using an enhanced chemiluminescence substrate system (ThermoFisher Scientific). The images were captured using a ChemiDoc™ Imaging System (Biorad).

### Immunohistochemistry

Immunohistochemical staining was examined on 7 μm wax sections of full-thickness retina and ONH. Sections from wax blocks from each group were used for immunohistochemical analysis. Tissues were incubated in 1% BSA/PBS for 1 h at room temperature to prevent non-specific background, then incubated in primary antibodies (Table S6) for 16 h at 4°C. After several washes, the tissues were incubated with the secondary antibodies for 4 h at 4°C. The tissues were then washed with PBS and counterstained with the nucleic acid stain Hoechst 33342 (1 μg/ml) in PBS. Images were acquired with Keyence All-in-One Fluorescence microscopy (BZ-X810, Keyence Corp. of America), and the fluorescent integrated intensity of each target protein in pixel per area was measured using the ImageJ software (National Institutes of Health). All imaging parameters remained the same and were corrected with background subtraction.

### Whole-mount immunohistochemistry and RGC counting

Mouse retinas from enucleated eyes were dissected as flattened whole-mounts. Retinas were immersed in 30% sucrose (prepared in PBS) for 24 h at 4°C. The retinas were then blocked in a blocking solution containing 3% donkey serum, 1% BSA, 1% fish gelatin, and 0.1% Triton X-100 to prevent non-specific binding. Primary antibodies, specifically rabbit polyclonal anti-RBPMS antibody or mouse monoclonal anti-TUJ1 antibody, were added and incubated for 3 days at 4°C. After several wash steps, the retinas are incubated with secondary antibodies labeled with Alexa Fluor-488 conjugated donkey anti-rabbit IgG antibody or Alexa Fluor-568 conjugated donkey anti-mouse IgG antibody for 24 hours at 4°C and subsequently washed with PBS. Images were acquired with Keyence All-in-One Fluorescence microscopy (BZ-X810, Keyence Corp. of America). RBPMS or TUJ1-positive RGCs were counted in two zones by middle and/or peripheral retina (three-sixths and five-sixths of the retinal radius), and the scores were averaged using ImageJ.

### Spatial Transcriptomic Profiling

Retinas from 10-month-old WT and *Akap1^−/−^* mice were dissected, embedded in OCT, and flash-frozen in LN2 gas. Frozen tissues were stored at –80°C and cryo-sectioned at 10 µm, with sections mounted onto Curio Seeker 3×3 spatial transcriptomics slides (Takara Curio Bioscience) according to the manufacturer’s protocol. Spatial transcriptomic processing included tissue fixation, permeabilization, capture of mRNA on spatially barcoded beads, on-chip reverse transcription, and indexed cDNA library construction, followed by sequencing on an Illumina platform. Raw sequencing data were processed using the LatchBio Curio Seeker pipeline, which performed read demultiplexing, alignment, barcode assignment, and generation of spatially resolved expression matrices. Downstream analysis focused on DEG between WT and *Akap1^−/−^*retinas, including normalization, scaling, PCA, clustering, and statistical testing, with visualization of spatial expression patterns via the LatchBio Seeker platform.

### Transcriptomic Profiling

Total RNA was extracted from mouse retinas using TRIzol™ reagent (Thermo Fisher Scientific), and RNA integrity was assessed by Azenta NGS Laboratory using an Agilent TapeStation, with all samples showing RNA Integrity Number (RIN) values above 8.4. Libraries for whole transcriptomic sequencing were constructed by GENEWIZ (Azenta Life Sciences) using the Illumina platform with 2 × 150 base pair (bp) paired-end reads. Four biological replicates from WT mice and six from *Akap1⁻^/^⁻* mice were analyzed. RNA sequencing data were processed using the ROSALIND® platform (https://rosalind.bio/), a cloud-based bioinformatics system. Adapter sequences were trimmed using Cutadapt, and read quality was assessed using FastQC (Fast Quality Control). Reads were aligned to the Mus musculus reference genome (mm10) using Spliced Transcripts Alignment to a Reference (STAR) (*89*). Gene-level quantification was performed with HTSeq (High-Throughput Sequencing Python framework) (*90*). Normalization and differential expression analysis were conducted using the DESeq2 package (Differential Expression analysis based on the Negative Binomial Distribution) with Relative Log Expression (RLE) normalization (*91*). Read Distribution percentages, violin plots, identity heatmaps, and sample MDS plots were generated as part of the QC step using RSeQC (*92*). Genes were considered significantly differentially expressed if they exhibited a fold change ≥ 1.25 and a false discovery rate (FDR) < 0.05. Clustering for heatmap visualization was performed using the Partitioning Around Medoids (PAM) algorithm from the Flexible Procedures for Clustering (fpc) R package. GO term pruning and dependency correction were performed using the topGO R package. Functional enrichment analysis was carried out using ROSALIND’s integrated tools, referencing Gene Ontology (GO), Interpro (*93*), NCBI (*94*), Molecular Signatures Database (MSigDB) (*95*), REACTOME (*96*), and WikiPathways (*97*).

### pERG and pVEP analyses

pERG and pVEP were measured with the Celeris apparatus based on the specifications provided by the manufacturer (Diagnosys LLC), as previously reported (*19, 98, 99*). pERG responses were recorded using alternating, reversing, black and white vertical stimuli at 1 Hz (2 reversals per second), and 50 candela/m^2^ delivered by the pattern stimulator. Then, 200 traces were recorded per eye, and averaged waveforms were calculated in which amplitudes (µV) were measured from the P1 peak to the N2 trough. At the same time, pVEP responses were recorded. For all the recording, ground and reference needle electrodes were placed subcutaneously in the tail and snout, and the active electrode was placed subcutaneously in the midline of the head at the location of the visual cortex. Each eye was exposed to 100 flashes of 1 Hz, 0.05 cd s/m^2^ white light through the corneal stimulators and recorded for 300 ms with a sample frequency of 2000 Hz. Then, 200 traces were recorded per eye, and averaged waveforms were calculated in which amplitudes (µV) were measured from the P1 peak to the N1 trough. We performed at least 3 trials for each mouse and averaged over 100 sweeps per trial. The low– and high-filter frequency cutoffs for pVEP were set to 1.25 Hz and 100 Hz. The data were analyzed using the software Espion V6 (Diagnosys).

### Virtual OMR analysis

Spatial visual function was performed on a virtual optomotor system (OptoMotry; CerebralMechanics Inc.), as previously reported (*19, 98, 100*). Briefly, mice were placed on a freely accessible platform positioned at the center of a virtual environment formed by four monitors arranged in a square. These screens displayed rotating vertical black-and-white bars (sinusoidal gratings) at a speed of 12 degrees per second. A camera mounted above the setup recorded the mice, and a forehead-aligned cursor ensured the rotation remained centered on the animal’s visual field. Visual acuity was evaluated by identifying periods when the mouse remained physically still, exhibiting only head-tracking movements. The spatial frequency threshold, a measure of visual acuity, was determined automatically with accompanying OptoMotry software, which uses a stepwise paradigm based upon head-tracking movements at 100% contrast.

### CTB labeling

CTB is an established marker for active uptake and transport used successfully to assess the retinal projection to the SC in injury, as previously reported (*19*). Mice were anesthetized with an IP injection of a mixture of ketamine/xylazine and topical 1% proparacaine eye drops. A Hamilton syringe was used to inject 1 µL of Alexa Fluor 594-conjugated CTB into the vitreous humor. Injections were given slowly over 1 min, and the needle was maintained in position for an additional 5 min to minimize CTB loss through the injection tract. At 3 days after injection, the mice were fixed via cardiac perfusion with 4% paraformaldehyde (Ted Pella) following an IP injection of a mixture of ketamine/xylazine. After perfusion, the SC tissues were dissected and immersed in PBS containing 30% sucrose for 24 h at 4°C. The SC tissues were coronally sectioned at 30 μm using a Leica Cryostat (Wetzlar). The 10 representative sections were mounted on slides, and images were acquired with Keyence All-in-One Fluorescence microscopy (BZ-X810, Keyence Corp. of America). The area densities from the images were analyzed using ImageJ.

### TEM analysis

RGCs were grown on MatTek petri dishes coated with poly-D-lysine and processed, as previously described (*19, 44, 98*). For fixation, the growth medium was poured out, and the primary fixative, composed of 2% paraformaldehyde, 2.5% glutaraldehyde (Ted Pella) in 0.15 M sodium cacodylate (pH 7.4), was added at 37°C for 2 minutes. Then the samples were gradually cooled by placing them on ice for 20 min. The cells were washed for 3 × 3 min in ice-cold sodium cacodylate buffer and incubated in a mixture of 1% OsO4, 0.8% potassium ferrocyanide, 3 μM calcium chloride in sodium cacodylate buffer for 30 min on ice. Then, the cells were washed 3 times for 3 min with ice-cold double-distilled water and stained with 2% uranyl acetate for 30 min on ice. The cells were next incubated in increasing ethanol solutions: 20%, 50%, 70%, 90% on ice, followed by 3 x 100% at room temperature, each for 3 min. Subsequently, the cells were infiltrated with a mixture of 50% ethanol and 50% Durcupan ACM resin (Fluka) for 60 min with agitation, and then incubated for 3 × 2 h in 100% Durcupan with agitation and polymerized for 48 h at 60°C in an oven. After removing the glass cover slip, cutting out a block about 2 mm across, and gluing it on a dummy block, thin sections about 70 nm thick were cut using a Leica ultramicrotome. The sections were placed on 200-mesh uncoated thin-bar copper grids. A Tecnai Spirit (FEI) electron microscope operated at 80 kV was used to record images with a Gatan 2Kx2K CCD camera at 2.9 nm/pixel. For quantitative analysis, the mitochondrial profile lengths and profile areas were measured with ImageJ (National Institutes of Health). The number of mitochondria was normalized to the total area occupied by RGC somas in each image, which was also measured using ImageJ. The mitochondrial volume density, defined as the volume occupied by mitochondria divided by the volume occupied by the cytoplasm, was estimated using stereology as follows. A 112 × 112 square grid (112 × 112 chosen for ease of use with Photoshop) was overlaid on each image loaded in Photoshop (Adobe), and mitochondria and cytoplasm lying under intercepts were counted. The relative volume occupied by mitochondria was expressed as the ratio of intercepts coinciding with this organelle relative to the intercepts coinciding with cytoplasm.

### SBEM analysis

Retina tissues for SBEM analysis were prepared as previously described (*19, 44, 98*). SBEM was performed on Gemini scanning electron microscopy (Zeiss) equipped with a 3view2XP and OnPoint backscatter detector (Gatan), as previously reported(*19, 101*). The retina volumes were collected at 2.5 kV accelerating voltages, with a pixel dwell time of 0.5μs, and the raster size was 20k x 5k, with 3.5 nm pixels and 50 nm z step size. Following the volume collection, the histograms for the tissues throughout the volume stack were normalized to correct for drift in image intensity during acquisition. Digital micrograph files (.dm4) were normalized using Digital Micrograph and then converted to MRC format. The stacks were converted to eight-bit, and volumes were manually traced for reconstruction and analysis using IMOD software (http://bio3d.colorado.edu/imod/). Mitochondrial morphology metrics were measured by using morpheme, a set of Python and Amira scripts (https://github.com/CRBS/morphome). Using the 3D model generated by the IMOD software, distance measurements were conducted within the same SBEM-derived volume using MTK commands in the 3dmod software package. Each vesicle in the ribbon-attached pool was individually labeled, and the shortest distance from each vesicle to the synaptic ribbon was calculated at nanometer resolution. Similarly, we measured the number of mitochondria located within defined distance ranges (100–2000 nm) from the target synaptic ribbon.

### Machine-learning-based segmentation and Analysis

CDeep3M, a deep learning-based segmentation tool (*59, 60*), was used to predict and generate various models from serially sectioned and segmented SBEM datasets. An initial pretrained model was further refined through extensive training using large-scale retinal SBEM data, enhancing predictive confidence and improving model robustness. This iterative training process provided a solid foundation for 3D reconstruction. Multiple output layers, including enhanced SBEM images and predicted structures, were integrated to reconstruct three-dimensional representations of cellular membranes, mitochondria, and synaptic vesicles. To quantify target organelles from multilayer predictions generated by CDeep3M, we utilized VAST-lite to overlay individual layers and assemble the segmented organelles into three-dimensional structures (*19, 102*). For quantitative analysis, VAST-tool, in combination with MATLAB, was utilized to measure morphometric parameters, including synaptic bouton volume, vesicle number and volume, and mitochondrial surface area, volume, and length. Cellular structures and internal organelles were reconstructed from SBEM datasets using the IMOD software suite and Blender (https://docs.blender.org/manual/en/2.81/) (*19, 98, 103, 104*). Synaptic ribbon bodies within the active zones of bouton structures were manually traced in each SBEM section and assembled into three-dimensional models. Vesicles belonging to the ribbon-attached pool and the proximal free pool(*59*) were similarly annotated across serial sections and integrated into the corresponding synaptic ribbon reconstructions.

### 3D EM tomography analysis

Semi-thick sections of thickness about 350 nm were cut from the blocks of cells prepared for TEM with a Leica ultramicrotome and placed on 200-mesh uncoated thin-bar copper grids, as previously reported (*2, 19, 44*). 20-nm colloidal gold particles were deposited on each side of the grid to serve as fiducial cues. The specimens were irradiated for about 20 min to limit anisotropic specimen thinning during image collection at the magnification used to collect the tilt series before initiating a dual-axis tilt series. During data collection, the illumination was held to near parallel beam conditions and the beam intensity was kept constant. Tilt series were captured using SerialEM software (University of Colorado) on a Tecnai HiBase Titan (FEI) electron microscope operated at 300 kV. Tilt series were taken at 0.81 nm/pixel. Images were recorded with a Gatan 4Kx4K CCD camera. Each dual-axis tilt series first collected 121 images taken at 1 degree increments over a range of –60 to +60 degrees, followed by rotating the grid 90 degrees and collecting another 121 images with the same tilt increment. After collecting the orthogonal tilt series, to improve the signal-to-noise ratio, 2x binning was performed on each image by averaging a 2×2 x-y pixel box into 1 pixel using the newstack command in IMOD (University of Colorado). The IMOD package (https://en.wikipedia.org/wiki/IMOD) was used for tilt-series alignment, reconstruction, and volume segmentation. R-weighted back projection was used to generate the reconstructions. Volume segmentation of mitochondrial membranes was performed by tracing in each of the 1.62 nm-thick x-y planes in which the object appeared. The Drawing Tools and Interpolator plug-ins created stacks of contours. The traced contours were then surface-rendered by turning the contours into meshes to generate a 3D model. The surface-rendered volumes were visualized using 3DMOD. Measurements of mitochondrial outer, inner boundary, and cristae membrane surface areas and volumes were made within segmented volumes using IMODinfo. These were used to determine the crista density, defined as the ratio: sum of the cristae membrane surface areas divided by the mitochondrial volume (units: µm⁻¹). The number of crista junctions per crista was determined by placing a sphere with a diameter about the same as the crista junction diameter at each crista junction in the volume. The sum of the crista junctions divided by the sum of the cristae gave the ratio, crista junction number per crista, for each segmented mitochondrion. The crista junction opening size was measured along the crista long-axis using 3DMOD.

### Modeling the rate of ATP production from EM tomography volumes

We used a biophysical modeling scheme, which takes into account spatially accurate geometric representations of a crista, inner boundary membrane, and crista junction, to estimate the rate of mitochondrial ATP generation, as previously reported (*62, 63, 98, 105*). The model of the rate of ATP production depends linearly on the cristae membrane surface area with high curvature housing the ATP synthase dimers according to the modeling concepts of the Garcia group (*62, 105*). The rate of ATP production per mitochondrial volume was calculated from the high-curvature crista density. The rate of ATP production per mitochondrion was also modeled by measuring the volume of each mitochondrion.

### OCR analysis

RGCs were seeded into Seahorse XF96-well plates at 1.5 × 10^4^ cells per well. The cells were treated with either AAV2 Null or AAV2-AKAP1 and allowed to grow and transduce for 3 days. After 3 days of culture, the cells were treated with 25 μM paraquat for 6 h to induce oxidative stress. OCR was measured using an XF96 pro analyzer (Agilent) (*2, 19, 31, 34*). For OCR analysis, after measuring the basal respiration, oligomycin (2 μg/ml, Sigma-Aldrich), an inhibitor of ATP synthesis, carbonyl cyanide 4-(trifluoromethoxy) phenylhydrazone (FCCP; 1 μM, Sigma-Aldrich), the uncoupler, and rotenone (2 μM, Sigma-Aldrich), an inhibitor of mitochondrial complex I, were sequentially added to measure maximal respiration, ATP-linked respiration, and spare respiratory capacity.

### Quantitative PCR

Isolated retina tissues were homogenized in 1 mL of TRIzol reagent (Invitrogen) and subjected to RNA isolation and purification, as previously described (*19*). The extracted RNA dissolved in RNase-free water was used for the reverse transcription reaction to obtain cDNA. Using 1 μg RNA, the reaction was performed to synthesize cDNA using the High-Capacity RNA-to-cDNA kit (Thermo Fisher Scientific). The mRNA expression level of each gene was quantified by real-time PCR using the SYBR Green PCR Master Mix (Applied Biosystems) according to the manufacturer’s instructions. The following PCR primer pairs (Table S7) were designed using primer-BLAST on the NCBI website to detect each gene: primers specific to mouse. The reaction conditions were as follows: 95°C for 5 min, 40 cycles at 95°C for 30 sec, 60°C for 30 sec, and 72°C for 20 sec. The elongation condition was 75°C for 7 min. The fold changes in mRNA levels were determined relative to the GAPDH mRNA level.

### MTT and LDH assay

Mitochondrial metabolic activity was assessed using the MTT assay (3-[4,5-dimethylthiazol-2yl]-2,5-diphenyl tetrazolium bromide; Cell Proliferation Kit I, Roche Diagnostics), following the manufacturer’s protocol, as previously reported (*31, 34, 44*). Primary RGCs or rMC-1 cells were transduced with either AAV2-AKAP1 or control (Null) vector for 3 days, followed by exposure to oxidative stress (PQ, 50 μM) or elevated HP (30 mmHg) for 24 hours. After treatment, 10 μL of MTT stock solution was added to each well, including untreated controls. Cells were incubated for 3–4 hours at 37 °C in a humidified 5% CO₂ incubator. Subsequently, 100 μL of solubilization solution (DMSO) was added to dissolve the formazan crystals.

Absorbance was measured at 560 nm with background correction at 690 nm using a microplate reader (SpectraMax, Molecular Devices). Cell cytotoxicity was evaluated using a colorimetric LDH (lactate dehydrogenase) release assay kit (Millipore Sigma), based on the manufacturer’s instructions. Following the same treatment conditions described above, 50 μL of cell culture supernatant from each well was transferred to a fresh 96-well plate. An equal volume (50 μL) of the LDH reaction mixture was added to each sample and incubated in the dark at room temperature for 30 minutes. Absorbance was recorded at 490 nm with background correction at 680 nm using the same microplate reader. The LDH level corresponds to the extent of plasma membrane damage and was expressed as a percentage of total LDH release. Each dataset was obtained from multiple replicate wells within each experimental group (*n* = 3).

### Intracellular ROS detection

Intracellular ROS production was evaluated using MitoSOX Red (500 nM, Invitrogen) following the manufacturer’s protocol. Primary RGCs were transduced with either AAV2-AKAP1 or control (Null) vector for 3 days, followed by exposure to oxidative stress (PQ, 50 μM) for 24 hours. After treatment, RGC cells were stained with MitoSOX Red (500 nM, Invitrogen) and MitoTracker Green (100 nM, Invitrogen) for 30 minutes at 37°C in culture medium. After staining, cells were washed twice with PBS to remove excess dye, and fluorescence was visualized using a Keyence All-in-One Fluorescence microscopy (BZ-X810, Keyence Corp. of America). Fluorescence intensity was quantified using ImageJ software to evaluate ROS levels.

### Calcium imaging and laser irradiation

RGCs were incubated in 2 μM Fluo4-AM (Biotium) in medium for 35 minutes before experiments. Cells were placed back into RGC medium without Fluo4-AM after washing 3x with Hank’s Buffered Saline Solution. A Ti-Saphire Laser emitting 740nm (Spectra-Physics) was passed through a polarizer to control the power that reaches the sample. A shutter controlled the laser exposure, and the beam was then passed through beam steering optics that directed the beam to the cut site. The beam was telescoped to fill the back aperture of a 40x Zeiss Objective with a numerical aperture of 1.3. Figure S19A shows a schematic of our experimental setup. The power before the objective was measured to be 250-300 mW. Fluorescence imaging was carried out utilizing an LED Xcite (Excelitas) set at an intensity of 2%. Cells were placed on an Ibidi humidified stage heater with 5% CO2 during imaging. Images were captured with an EMCCD camera (QuantEM:512SC, Photometrics). A line perpendicular to the axon was irradiated with the laser to induce axonal injury. This line was drawn 7.5 μm away from the edge of the cell body (Fig. S19B). The mean pixel intensity at the cell body and 3 locations on the axon were quantified to monitor Ca^2+^ changes (Fig. S19C). To quantify calcium changes, the background was subtracted, and mean pixel values were normalized to pre-laser by dividing by pixel intensities before laser ablation. The percentage change can thus be calculated by subtracting 1 and multiplying by 100.

## Statistical analysis

All results were reported as means ± SEM. For comparison between two groups, statistical analysis was performed using a two-tailed unpaired Student’s t-test. For multiple group comparisons, we used one-way analysis of variance (ANOVA) and Tukey’s multiple comparisons test, using GraphPad Prism. A *P* value less than 0.05 was considered statistically significant. \**P* < 0.05, \*\**P* < 0.01, \*\*\**P* < 0.001, \*\*\*\**P* < 0.0001. The value of n per group is indicated within each figure legend.

## Acknowledgments

We thank Dr. Stefan Strack (University of Iowa) for providing the AKAP1 plasmid and *Akap1⁻^/^⁻* mice.

## Funding

National Institutes of Health grant R01 EY031697 (WKJ, GAP) National Institutes of Health grant R01 EY034116 (WKJ, SHC, KYK) National Institutes of Health grant P30 EY022589 (Vision core grant) Research to Prevent Blindness (New York, NY) University of California Gene Therapy Initiative (WKJ)

## Author contributions

Conceptualization: WKJ

Methodology: TB, KYK, ZS, GAP, MP, VG, YL, HC, SHC, WKJ

Investigation: TB, KYK, ZS, GAP, MP, VG, YL, HC, JYW, WKJ

Visualization: TB, KYK, ZS, GAP, WKJ

Funding acquisition: KYK, GAP, SHC, WKJ

Project administration: WKJ

Supervision: KYK, WKJ

Writing – original draft: TB, ZS, KYK, GAP, WKJ

Writing – review & editing: TB, KYK, GAP, MHE, RNW, WKJ

## Competing interests

Authors declare that they have no competing interests.

## Data and materials availability

All data are available in the main text or the supplementary materials.

## Notes

### Competing Interest Statement

The authors have declared no competing interest.

