## Supplemental Information for "AKAP1 regulates mitochondrial and synaptic homeostasis to enable neuroprotection and repair in retinal ganglion cell degeneration"

Graphic abstract

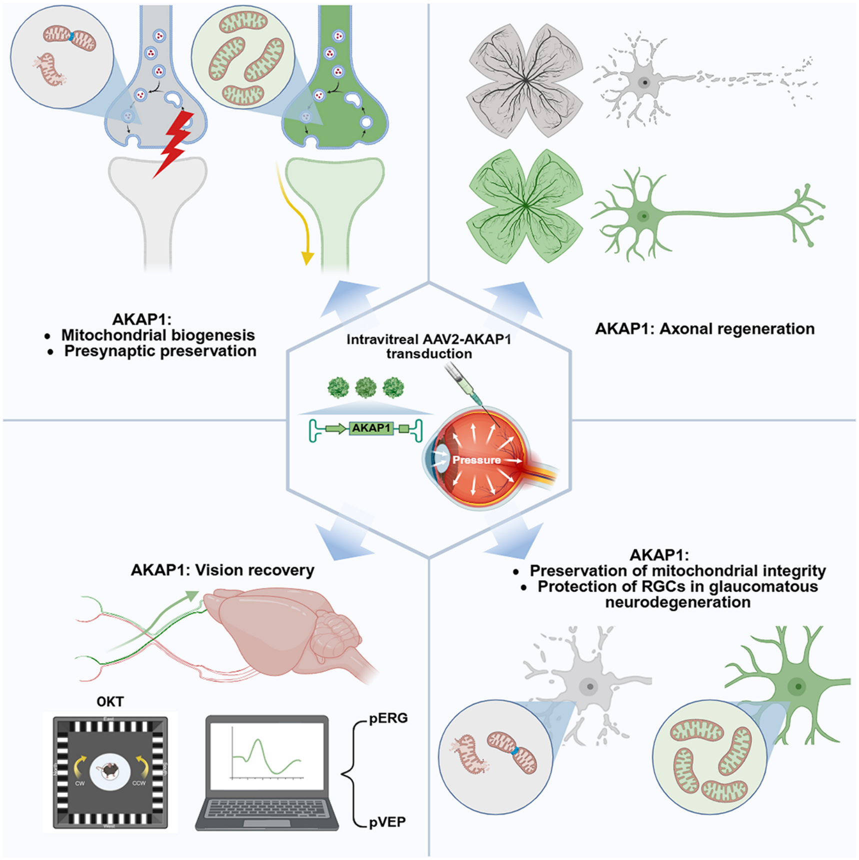

### Highlights

- AKAP1 is markedly downregulated in glaucomatous human and mouse retinas, implicating its deficiency in RGC vulnerability.
- AKAP1 expression enhances mitochondrial fusion and energy production, preserving mitochondrial integrity and function in RGCs.
- AKAP1 promotes mitochondrial biogenesis and maintains synaptic integrity, protecting retinal synapses from glaucomatous neurodegeneration.
- AKAP1 facilitates axonal regeneration, preserves the visual pathway, and sustains visual function.

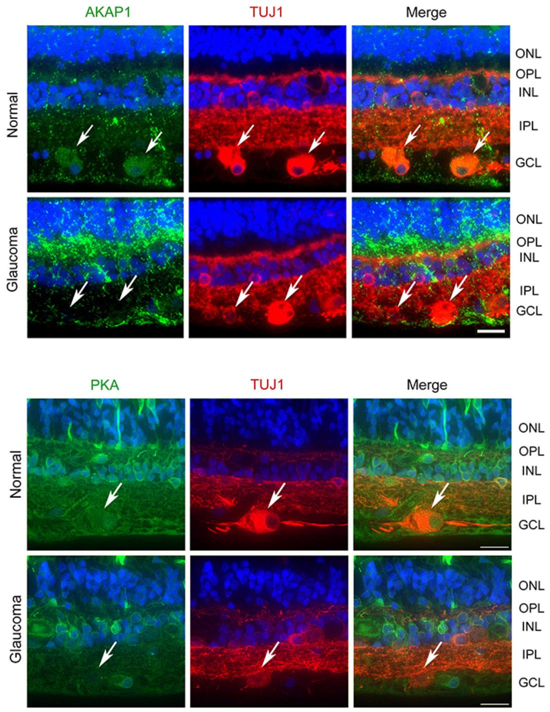

**Fig. S1** AKAP1 and PKA expression in glaucomatous human retina. Representative retinal image for AKAP1 and PKA (green) and TUJ1 (red) immunoreactivities. Arrows indicate AKAP1 immunoreactivity co-labeled with TUJ1 in RGC somas. Scale bars, 20 µm.

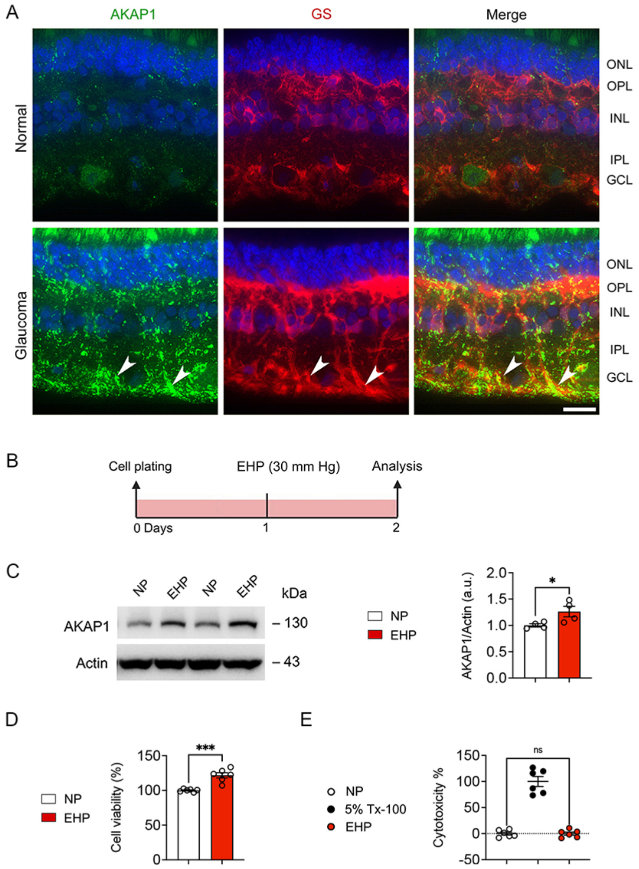

**Fig. S2** AKAP1 and GS expression in glaucomatous human retina. (A) Representative retinal images for AKAP1 (green) and GS (red) immunoreactivities. Arrows indicate AKAP1 immunoreactivity co-labeled with GS-positive Müller glial cells in the retina. (B) Experimental schematic and timeline, and data analysis in Müller glial cells (rMC-1) under elevated HP conditions *in vitro*. (C) Representative image of a western blot and densitometry graph for AKAP1 expression in Müller glial cells under elevated HP conditions *in vitro*. ($n$ = 4 biological replicates per group). (D and E) Quantitative analyses of Müller glial cell viability and death using MTT (D) and LDH release (E) assays under elevated HP conditions ($n$ = 6 biological replicates per group). Error bars represent SEM. Statistical analysis was performed using unpaired Student's $t$-test or one-way ANOVA and Tukey's multiple comparisons test. ***$P$ < 0.001. Scale bars, 20 µm. EHP, elevated hydrostatic pressure.

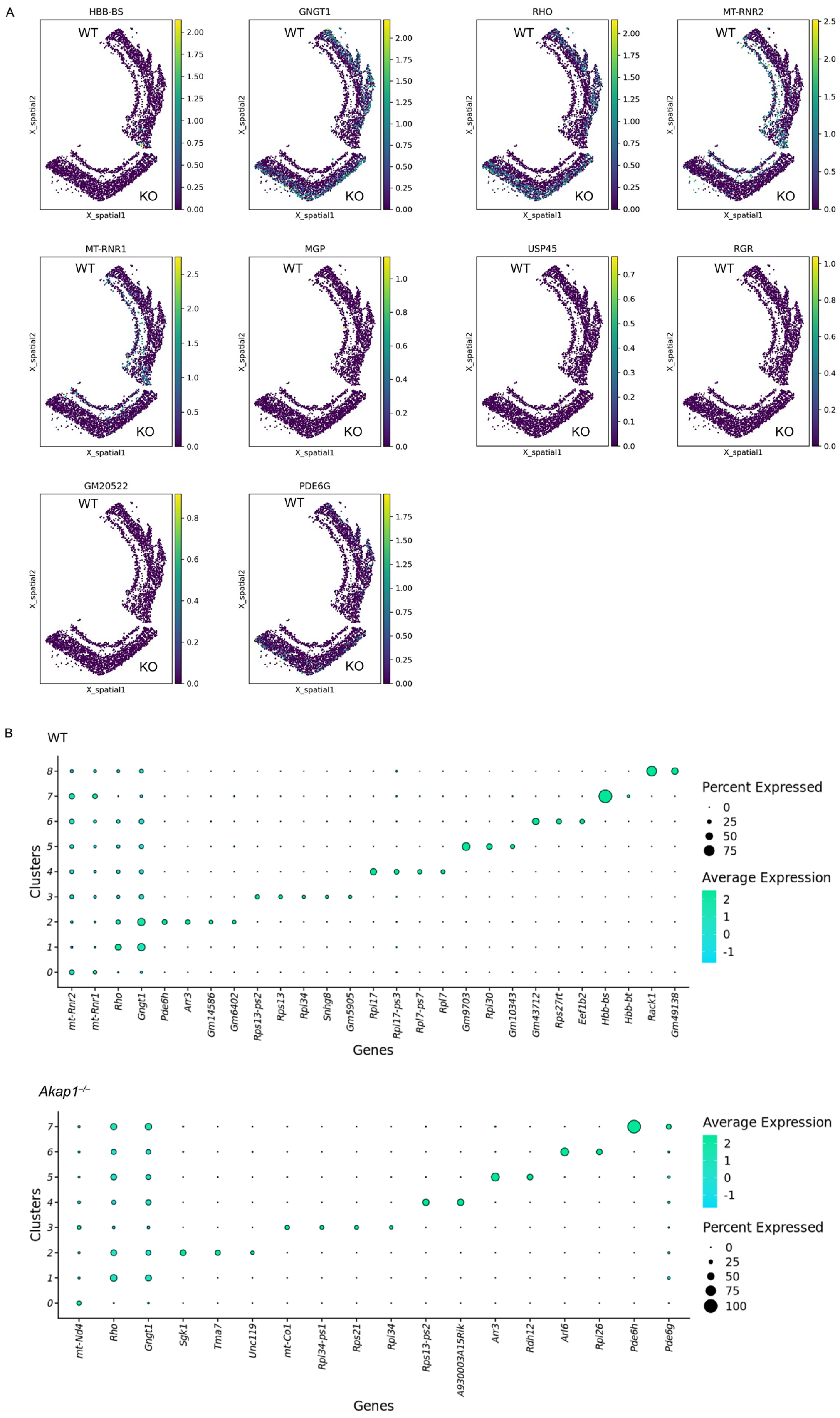
A
HBB-BS
GNGT1
RHO
MT-RNR2
MT-RNR1
MGP
USP45
RGR
GM20522
PDE6G
WT
KO
X_spatial1
X_spatial2
B
WT
Percent Expressed
0
25
50
75
Average Expression
2
1
0
-1
Clusters
Genes
mt-Rnr2
mt-Rnr1
Rho
Gngt1
Pde6h
Arr3
Gm14586
Gm6402
Rps13-ps2
Rps13
Rpl34
Snhg8
Gm5905
Rpl17
Rpl17-ps3
Rpl7-ps7
Rpl7
Gm9703
Rpl30
Gm10343
Gm43712
Rps27rt
Eef1b2
Hbb-bs
Hbb-bt
Rack1
Gm49138
Akap1−/−
Average Expression
2
1
0
-1
Percent Expressed
0
25
50
75
100
Clusters
Genes
mt-Nd4
Rho
Gngt1
Sgk1
Tma7
Unc119
mt-Co1
Rpl34-ps1
Rps21
Rpl34
Rps13-ps2
A930003A15Rik
Arr3
Rdh12
Arl6
Rpl26
Pde6h
Pde6g

**Fig. S3** Spatial transcriptomic profiling in WT and *Akap1*$^{-/-}$ retinas. (A) Spatial map of retinal sections showing the top spatially variable genes. (B) Dot plot of the top differentially expressed genes per cluster in WT and *Akap1*$^{-/-}$ retinas.

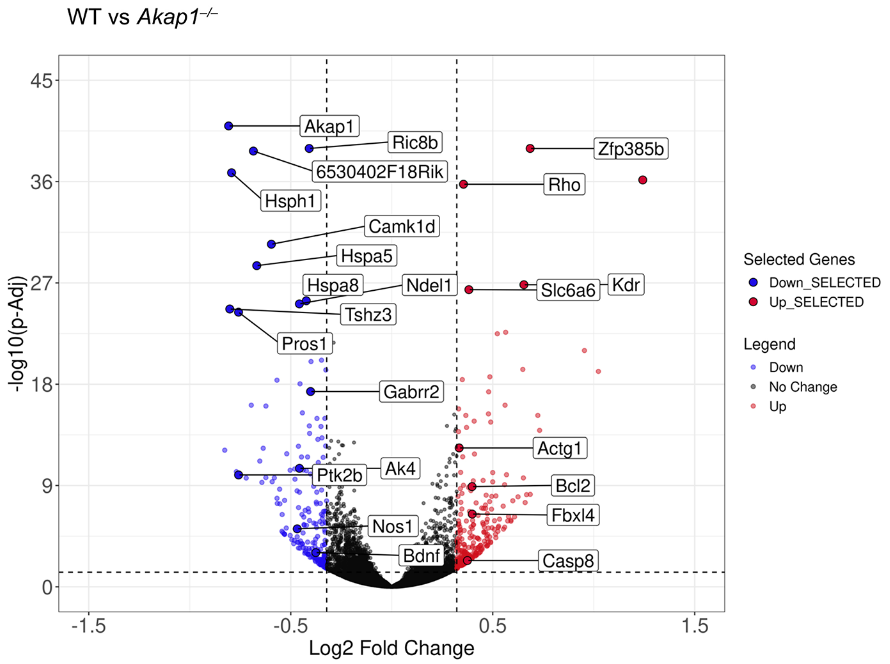

**Fig. S4** Transcriptomic profiling of retinas from WT and *Akap1*$^{-/-}$ mice. Volcano plot showing differentially expressed genes (DEGs) between WT and *Akap1*$^{-/-}$ retinas. A total of 244 genes were significantly upregulated and 196 genes were downregulated in the *Akap1*$^{-/-}$ group (fold change ≥ 1.25; adjusted $p < 0.05$).

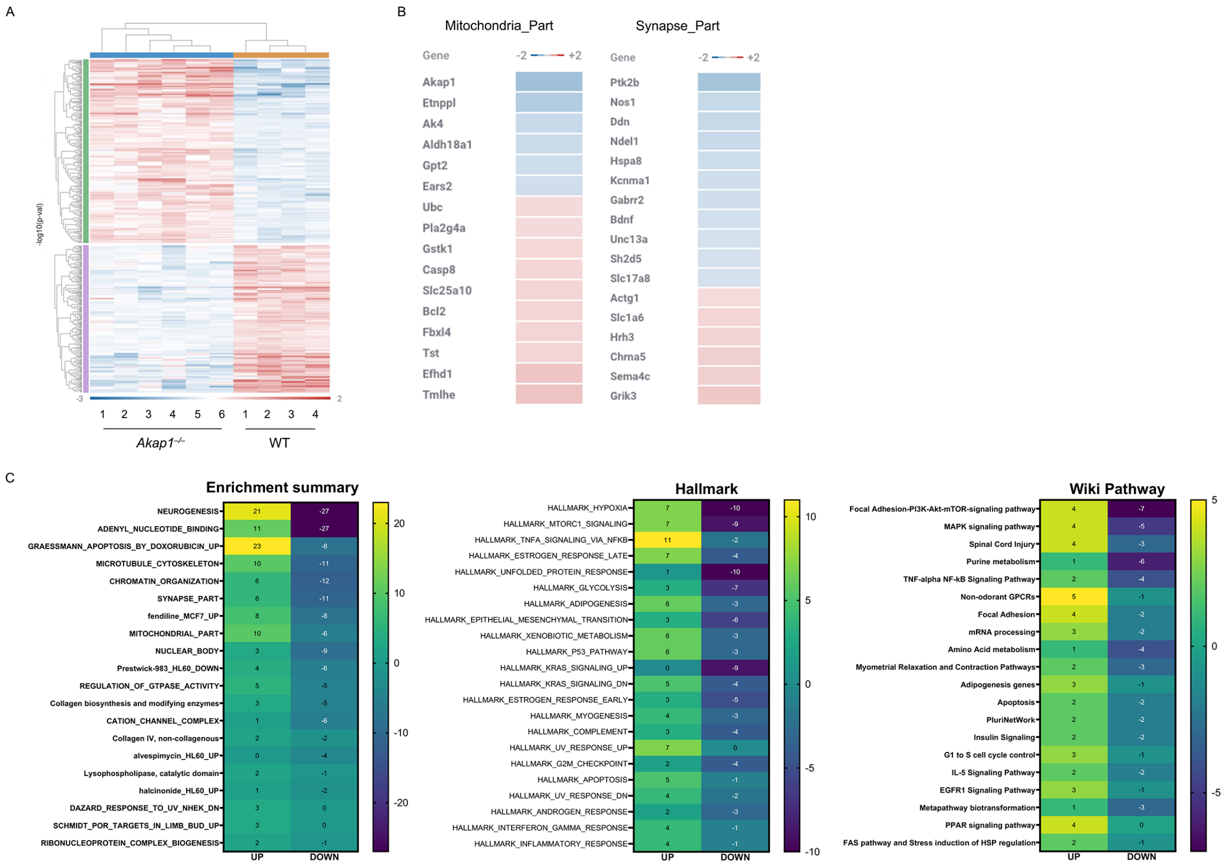

**Fig. S5** Transcriptomic profiling of retinas from WT and *Akap1*$^{-/-}$ mice. (A) Heatmap representing hierarchical clustering of DEGs, highlighting distinct gene expression patterns between WT and *Akap1*$^{-/-}$ retinas. (B) Gene ontology (GO) enrichment analysis of DEGs reveals enrichment of genes associated with mitochondrial components and synaptic structure, suggesting that AKAP1 influences mitochondrial function and neuronal signaling. (C) Functional enrichment summary, including Gene Ontology (GO), Hallmark gene sets, and WikiPathways, identifies pathways related to neurogenesis, synaptic signaling, and mitochondrial function, as significantly affected by AKAP1 deletion.

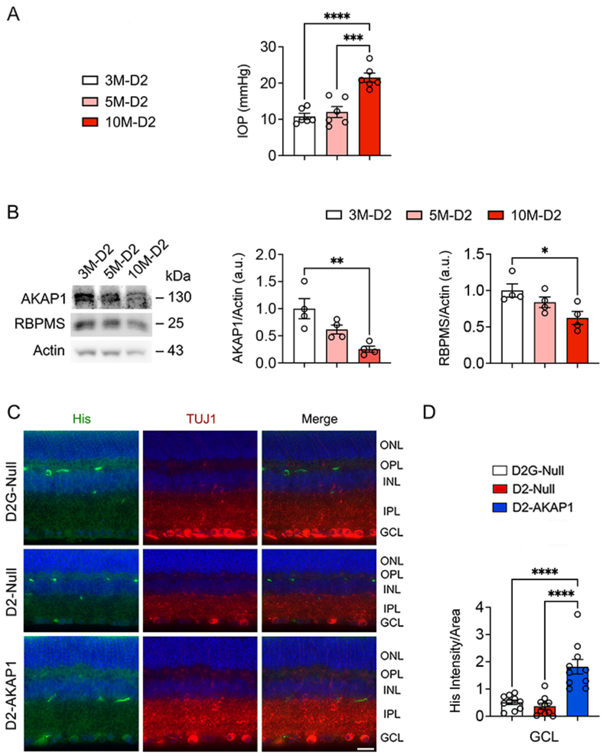

**Fig. S6** Decreased AKAP1 expression in non-treated glaucomatous D2 retina and His immunoreactivity in glaucomatous D2-AKAP1 retina. (A) IOP measurement in 3-, 5-, and 10-month-old non-treated D2 mice ($n$ = 6 mice per group). (B) Representative image of a western blot and densitometry graph for AKAP1 and RBPMS in the non-treated glaucomatous D2 retinas ($n$ = 4 retinas per group). (C) Representative retinal images for His (green) and TUJ1 (red) immunoreactivities in glaucomatous D2-AKAP1 retina. (D) Quantitative fluorescent intensity of His immunoreactivity in the GCL ($n$ = 10 retina sections from 3 mice per group). Error bars represent SEM. Statistical analysis was performed using one-way ANOVA and Tukey's multiple comparisons test. *$P < 0.05$, **$P < 0.01$, ***$P < 0.001$, and ****$P < 0.0001$. Scale bars, 20 µm.

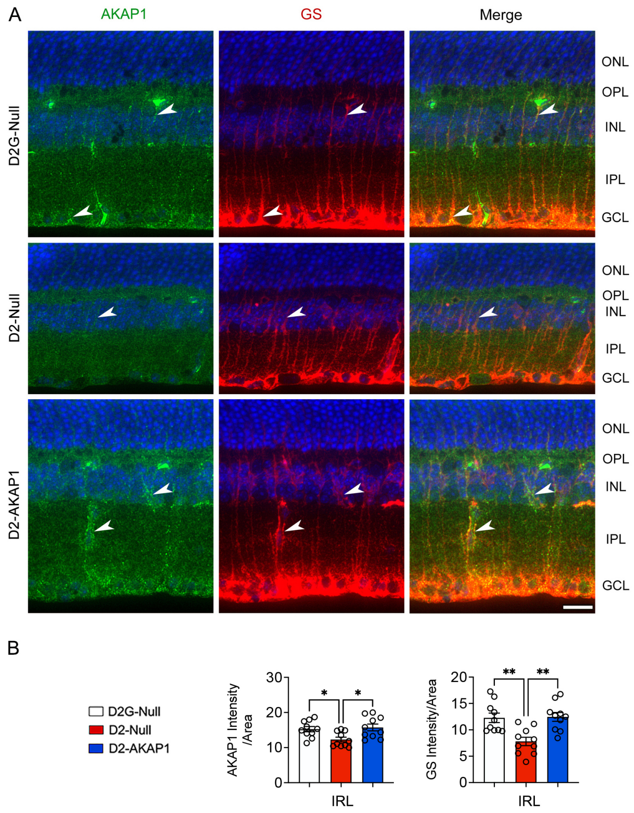

**Fig. S7** Increased AKAP1 expression in Müller glial cells of glaucomatous D2-AKAP1 retina. (A) Representative retinal images for AKAP1 (green) and GS (red) immunoreactivities from D2G-Null, D2-Null, and D2-AKAP1 retinas. Arrowheads indicate AKAP1 immunoreactivity co-labeled with GS-positive Müller glial cells. (B) Quantitative fluorescent intensity of AKAP1 and GC immunoreactivities in the inner retinal layer (IRL) ($n$ = 10 retina sections from 3 mice per group). Error bars represent SEM. Statistical analysis was performed using one-way ANOVA and Tukey's multiple comparisons test. *$P < 0.05$ and **$P < 0.01$. Scale bars, 20 μm.

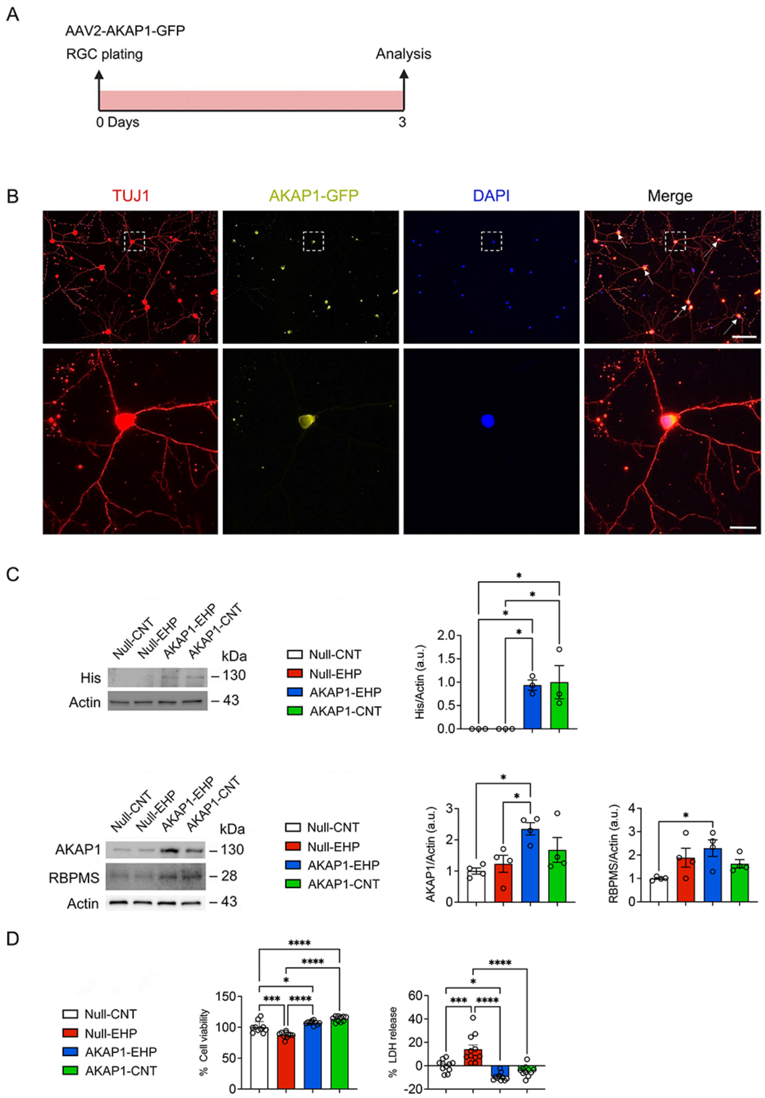

**Fig. S8** Amplifying AKAP1 expression promotes RGC survival under elevated HP conditions. (A) Experimental schematic and timeline of AAV2-AKAP1 or AAV2-AKAP1-GFP transduction in cultured primary RGCs and data analysis. (B) Representative images for AKAP1-GFP (yellow) expression in TUJ1(red)-labeled RGCs. (C) Western blot analysis for His, AKAP1, and RBPMS expression in cultured RGCs (*n* = 3 or 4 biological replicates per group). (D) Quantitative analyses of RGC viability and death using MTT and LDH release assays under elevated HP conditions (*n* = 12 biological replicates per group). Error bars represent SEM. Statistical analysis was performed using one-way ANOVA and Tukey's multiple comparisons test. *$P < 0.05$, ***$P < 0.001$, and ****$P < 0.0001$. Scale bars, 20 μm. EHP, elevated hydrostatic pressure.

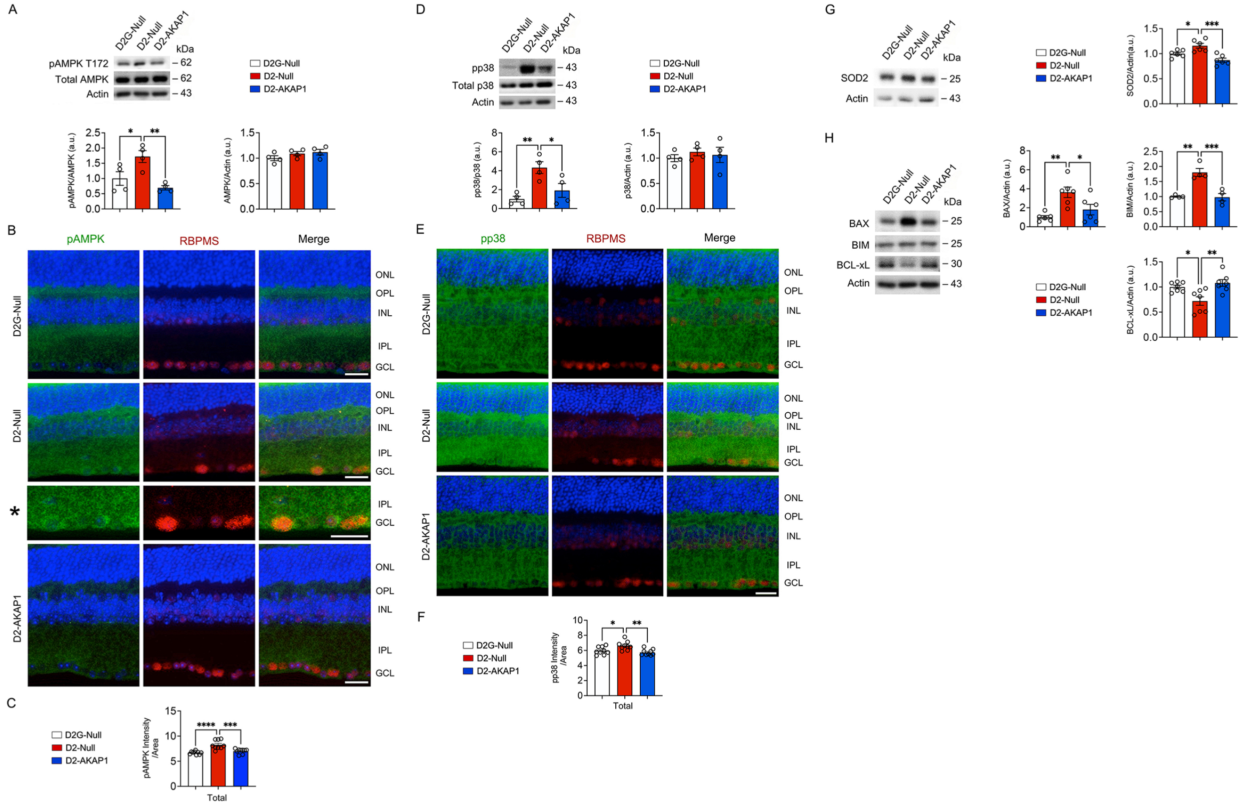

**Fig. S9** Restoring AKAP1 expression inhibits AMPK and P38 activation, oxidative stress, and apoptosis in glaucomatous D2 retina. (A) Western blot analysis for phospho-AMPK (pAMPK) and total AMPK expression in the retinas (*n* = 4 retinas per group). (B) Representative retinal images for pAMPK (green) and RBPMS (red) immunoreactivities. Asterisk (*) indicates high magnification for the representative image (pAMPK and RBPMS) for D2-Null retina. (C) Quantitative fluorescent intensity of pAMPK immunoreactivity in the total retinal layer (*n*=9 retinal sections from 3 mice per group). (D) Western blot analysis for phospho-p38 (pp38) and total p38 expression in the retinas (*n* = 4 retinas per group). (E) Representative retinal images for pp38 (green) and RBPMS (red) immunoreactivities. (F) Quantitative fluorescent intensity of pp38 immunoreactivity in the total retinal layer. (*n*=9 retinal sections from 3 mice per group). (G) Western blot analysis for SOD2 expression in the retinas (*n* = 6 retinas per group). (H) Western blot analysis for BAX, BIM, and BcL-xL expression in the retinas (*n* = 4 to 7 retinas per group). Error bars represent SEM. Statistical analysis was performed using one-way ANOVA and Tukey's multiple comparisons test. *$P < 0.05$, **$P < 0.01$, ***$P < 0.001$ and ****$P < 0.0001$. Scale bars, 20 μm.

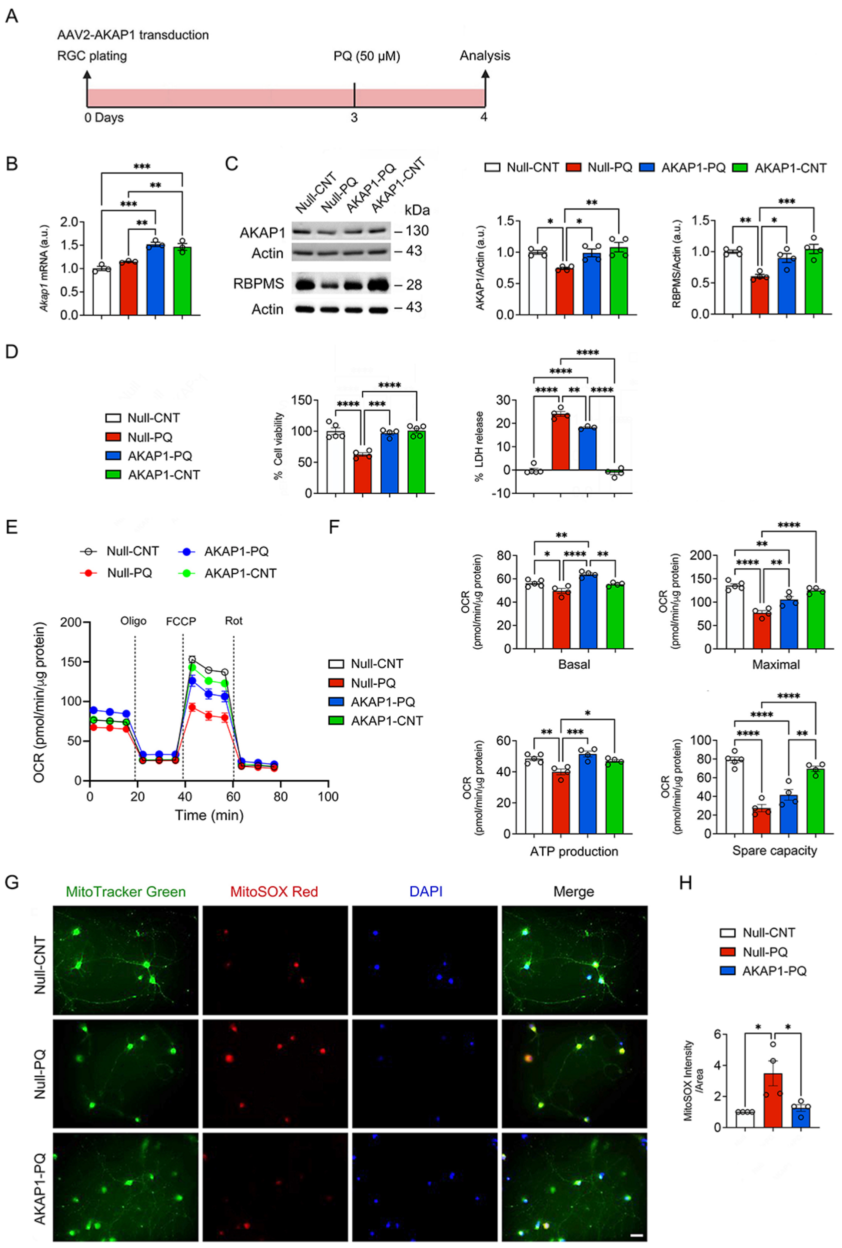

**Fig. S10** Amplifying AKAP1 expression protects RGCs against PQ-induced oxidative stress. (A) Experimental schematic and timeline of AAV2-AKAP1 transduction and PQ (50 μM) treatment in cultured primary RGCs and data analysis. (B) Quantitative PCR analysis for *Akap1* gene expression in cultured RGCs under oxidative stress conditions ($n$ = 3 biological replicates per group). (C) Western blot analysis for AKAP1 and RBPMS expression in cultured RGCs under oxidative stress conditions ($n$ = 4 biological replicates per group). (D) Quantitative analyses of RGC viability and death using MTT and LDH release assays under oxidative conditions ($n$ = 4 or 5 biological replicates per group). (E) OCR changes in AAV2-AKAP1-treated RGCs under PQ-induced oxidative stress conditions. (F) Quantitative analysis of OCR parameters: basal respiration, maximal respiration, ATP production, and spare capacity in cultured RGCs under oxidative stress conditions ($n = 5$ biological replicates per group). (G) Representative images for Mito-tracker green (green), MitoSOX red (red), and Hoechst (blue) staining in cultured RGCs under oxidative stress conditions. (H) Quantitative fluorescent intensity of MitoSOX (red) ($n = 4$ biological replicates per group). Error bars represent SEM. Statistical analysis was performed using one-way ANOVA and Tukey's multiple comparisons test. $^{*}P < 0.05$, $^{**}P < 0.01$, $^{***}P < 0.001$ and $^{****}P < 0.0001$. Scale bars, 20 μm.

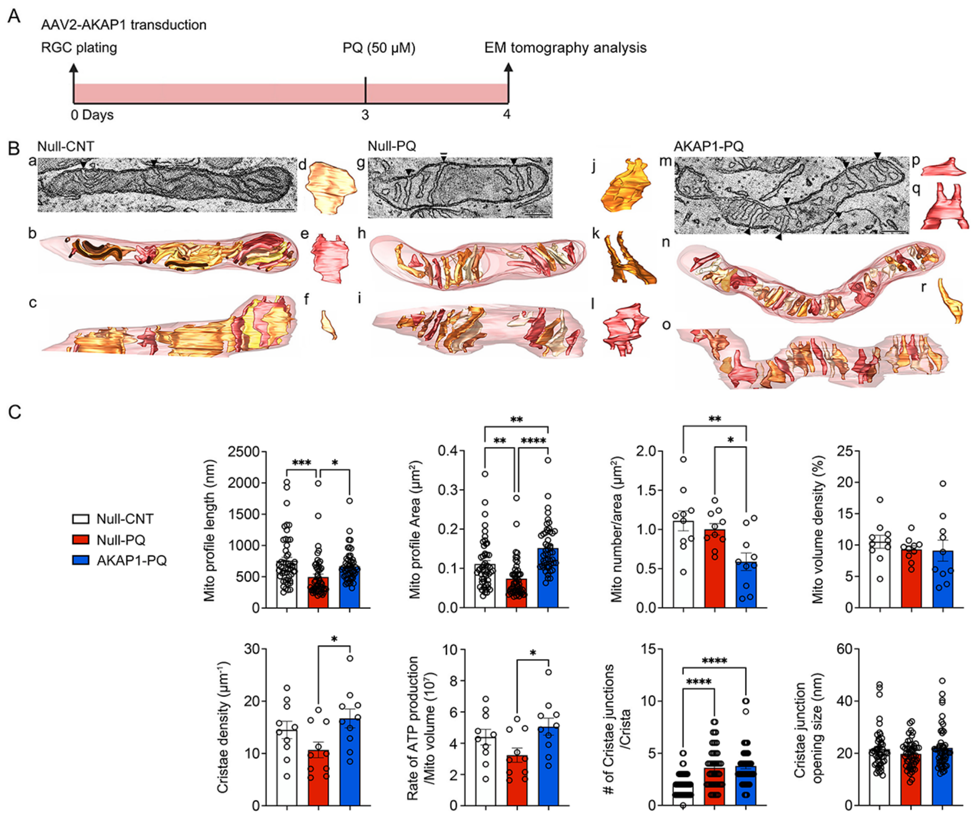

**Fig. S11** Amplifying AKAP1 expression preserves mitochondrial dynamics and promotes ATP production in RGCs against PQ-induced oxidative stress. (A) Experimental schematic and timeline of AAV2-AKAP1 transduction and PQ (50 μM) treatment in cultured primary RGCs and data analysis. (B) Representative TEM images of mitochondrial morphology with 3D-reconstruction and cristae junction (arrowhead) of Null-CNT (a-f), Null-PQ (g-l), and AKAP1-PQ (m-r). (C) Measurements for mitochondrial profile length (*n* = 20 mitochondria per group), mitochondrial profile area (n = 20 mitochondria per group), number per unit area (*n* = 10 mitochondria per group), volume density (*n* = 10 mitochondria per group), cristae density (n=10 mitochondria per group), rate of ATP generation per mitochondrial volume (*n* = 10 mitochondria per group), number of crista junction per crista (n = 10 mitochondria per group) and crista junction width (*n* = 20 mitochondria per group). Bars represent SEM. Statistical analysis was performed using one-way ANOVA and Tukey's multiple comparisons test. *$P < 0.05$, **$P < 0.01$, ***$P < 0.001$ and ****$P < 0.0001$. Scale bars, 500 nm.

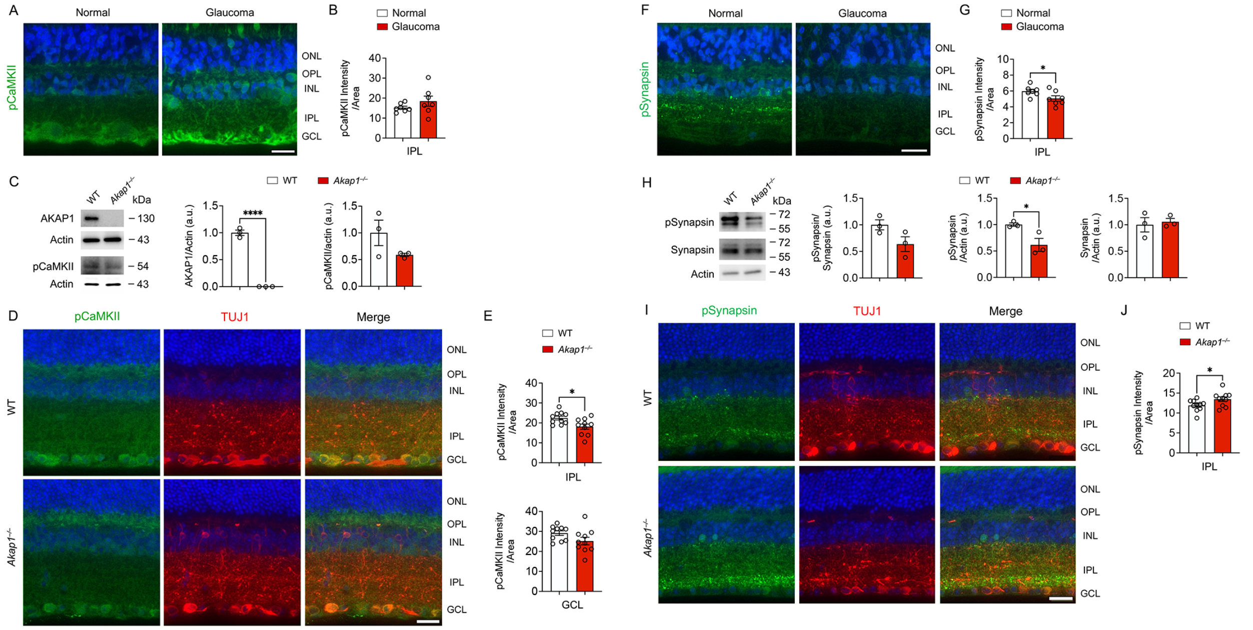

**Fig. S12** Phosphorylated CaMKII and Synapsin expression in glaucomatous human and *Akap1*$^{-/-}$ mouse retinas. (A) Representative retinal images for phospho-CaMKII (pCaMKII, green) immunoreactivity in normal and glaucomatous human retinas. (B) Quantitative fluorescent intensity of pCaMKII immunoreactivity in the IPL of the glaucomatous human retinas ($n$ = 7 retinal sections from 7 eyes from donors/group). (C) Western blot analysis for AKAP1 and pCaMKII expression in WT and *Akap1*$^{-/-}$ retinas ($n$ = 3 retinas per group). (D) Representative retinal images for pCaMKII immunoreactivity in WT and *Akap1*$^{-/-}$ retinas. (E) Quantitative fluorescent intensity of pCaMKII immunoreactivity in the IPL and GCL of the WT and *Akap1*$^{-/-}$ retinas ($n$=9 retinal sections from 3 mice per group). (F) Representative retinal images for phospho-Synapsin (pSynapsin, green) immunoreactivity in normal and glaucomatous human retinas. (G) Quantitative fluorescent intensity of pSynapsin immunoreactivity in the IPL of the normal and glaucomatous human retinas ($n$ = 7 retinal sections from 7 eyes from donors/group). (H) Western blot analysis for total synapsin and pSynapsin expression in WT and *Akap1*$^{-/-}$ retinas ($n$ = 3 retinas per group). (I) Representative retinal images for pSynapsin immunoreactivities in WT and *Akap1*$^{-/-}$ retinas. (J) Quantitative fluorescent intensity of pSynapsin immunoreactivity in the IPL of the WT and *Akap1*$^{-/-}$ retinas ($n$=9 retinal sections from 3 mice per group). Error bars represent SEM. Statistical analysis was performed using an unpaired Student's $t$-test. *$P$ < 0.05 and ****$P$ < 0.0001. Scale bars, 20 μm.

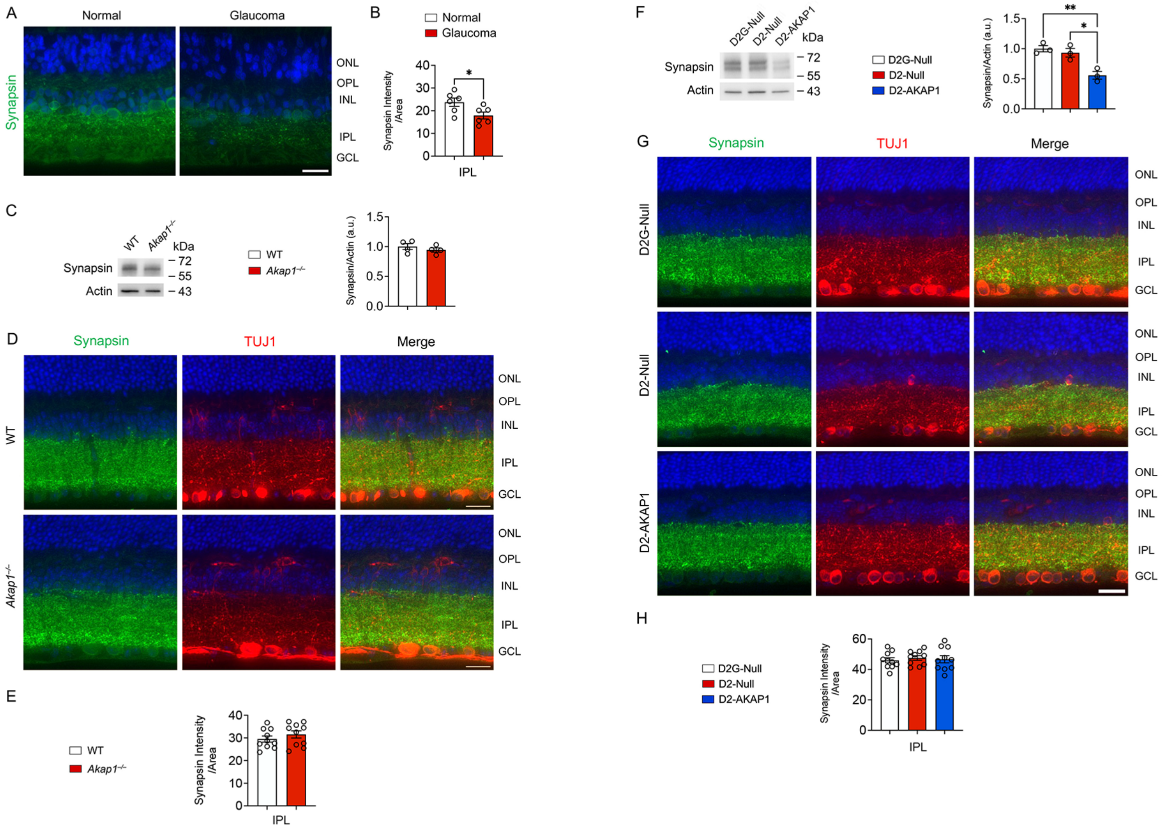

**Fig. S13** Synapsin expression in glaucomatous human and mouse retinas, as well as *Akap1*$^{-/-}$ mouse retinas. (A) Representative retinal images for total synapsin (green) immunoreactivity in normal and glaucomatous human retinas. (B) Quantitative fluorescent intensity of total synapsin immunoreactivity in the IPL of the glaucomatous human retinas ($n$ = 7 retinal sections from 7 eyes from donors/group). (C) Western blot analysis for total synapsin expression in WT and *Akap1*$^{-/-}$ retinas ($n$ = 4 retinas from 4 mice per group). (D) Representative retinal images for total synapsin immunoreactivity in WT and *Akap1*$^{-/-}$ retinas. (E) Quantitative fluorescent intensity of total synapsin immunoreactivity in the IPL of the WT and *Akap1*$^{-/-}$ retinas ($n$=9 retinal sections from 3 mice per group). (F) Western blot analysis for total synapsin expression in D2G and D2 retinas ($n$ = 3 retinas from 3 mice per group). (G) Representative retinal images for total synapsin immunoreactivity in D2G and D2 retinas. (H) Quantitative fluorescent intensity of total synapsin immunoreactivity in the IPL of D2G and D2 retinas ($n$ = 9 retinal sections from 3 mice per group). Error bars represent SEM. Statistical analysis was performed using an unpaired Student's $t$-test or one-way ANOVA and Tukey's multiple comparisons test. *$P < 0.05$, **$P < 0.01$ and ***$P < 0.001$. Scale bars, 50 μm.

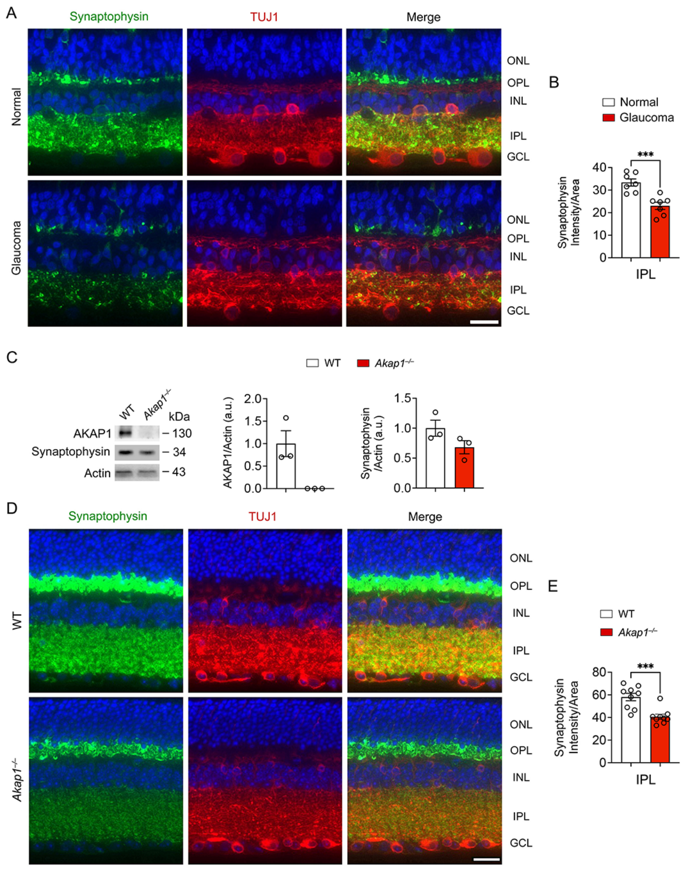

**Fig. S14** Synaptophysin expression in glaucomatous human and *Akap1*$^{-/-}$ mouse retinas. (A) Representative retinal images for synaptophysin (green) and TUJ1 (red) immunoreactivities in normal and glaucomatous human retinas. (B) Quantitative fluorescent intensity of synaptophysin immunoreactivity in the IPL of the normal and glaucomatous human retinas ($n$ = 7 retinal sections from 7 eyes from donors/group). (C) Western blot analysis for synaptophysin expression in WT and *Akap1*$^{-/-}$ retinas ($n$ = 3 retinas per group). (D) Representative retinal images for synaptophysin immunoreactivity in WT and *Akap1*$^{-/-}$ retinas. (E) Quantitative fluorescent intensity of synaptophysin immunoreactivity in the IPL of the WT and *Akap1*$^{-/-}$ retinas ($n$=9 retinal sections from 3 mice per group). Error bars represent SEM. Statistical analysis was performed using an unpaired Student's $t$-test. ***$P$ < 0.001. Scale bars, 20 μm.

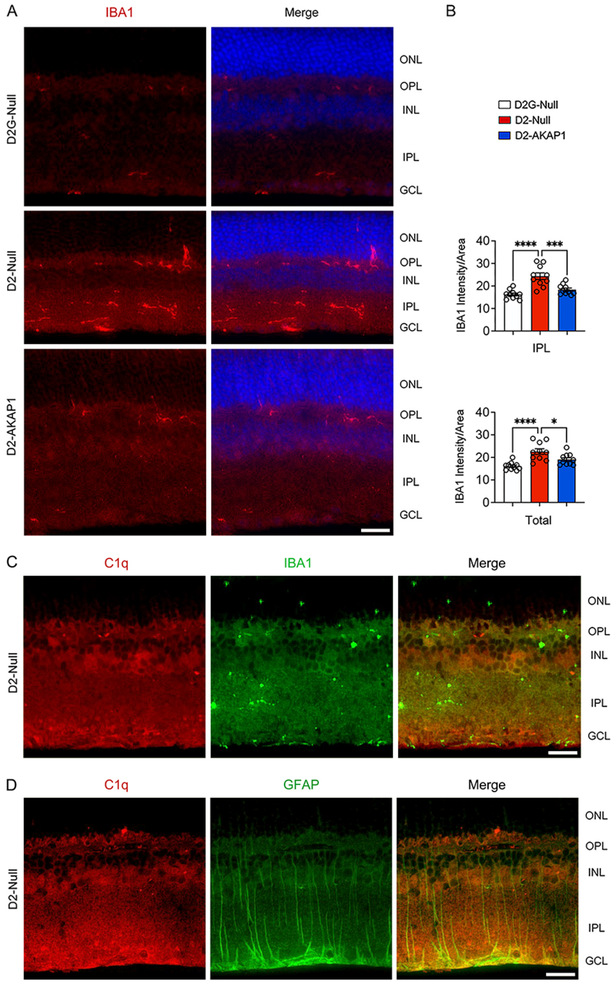

**Fig. S15** Restoring AKAP1 expression significantly decreases IBA1 protein expression in glaucomatous D2 retinas. (A) Representative retinal images for IBA1 (red) immunoreactivities in glaucomatous D2 retinas. (B) Quantitative fluorescent intensity of IBA1 immunoreactivity in the IPL and total layer of the normal and glaucomatous human retinas ($n$ = 10 retinal sections from 3 mice per group). (C) Representative retinal images for C1q (red) and IBA1 (green) immunoreactivities in glaucomatous D2 retinas. (D) Representative retinal images for C1q (red) and GFAP (green) immunoreactivities in glaucomatous D2 retinas. Error bars represent SEM. Statistical analysis was performed using one-way ANOVA and Tukey's multiple comparisons test. *$P < 0.05$, ***$P < 0.001$ and ****$P < 0.0001$. Scale bars, 20 μm.

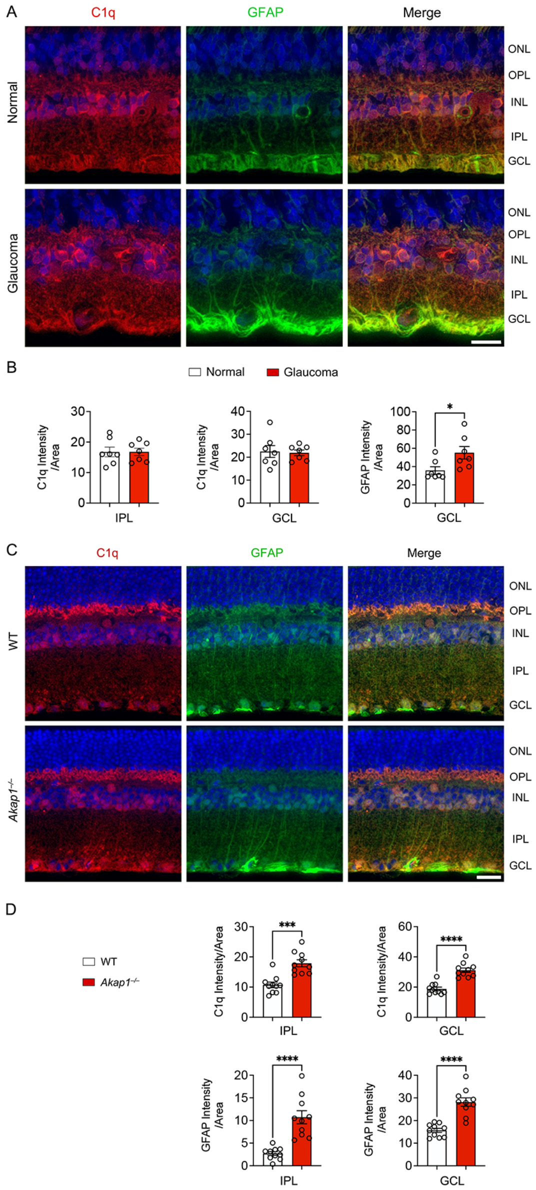

**Fig. S16** Increased C1q expression in glaucomatous human and *Akap1*$^{-/-}$ retinas. (A) Representative retinal images for C1q (red) and GFAP (green) immunoreactivity in normal and glaucomatous human retinas. (B) Quantitative fluorescent intensity of C1q immunoreactivity in the IPL and GCL of the glaucomatous human retinas ($n$ = 7 retinal sections from 7 eyes from donors/group). (C) Quantitative fluorescent intensity of C1q immunoreactivity in the IPL and GCL in WT and *Akap1*$^{-/-}$ retinas. (D) Quantitative fluorescent intensity of C1q and GFAP immunoreactivities in the IPL and GCL of the WT and *Akap1*$^{-/-}$ retinas ($n$ = 9 or 10 retinal sections from 3 mice per group). Bars represent SEM. Error bars represent SEM. Statistical analysis was performed using an unpaired Student's $t$-test. *$P < 0.05$, ***$P < 0.001$, and ****$P < 0.0001$. Scale bars, 20 μm.

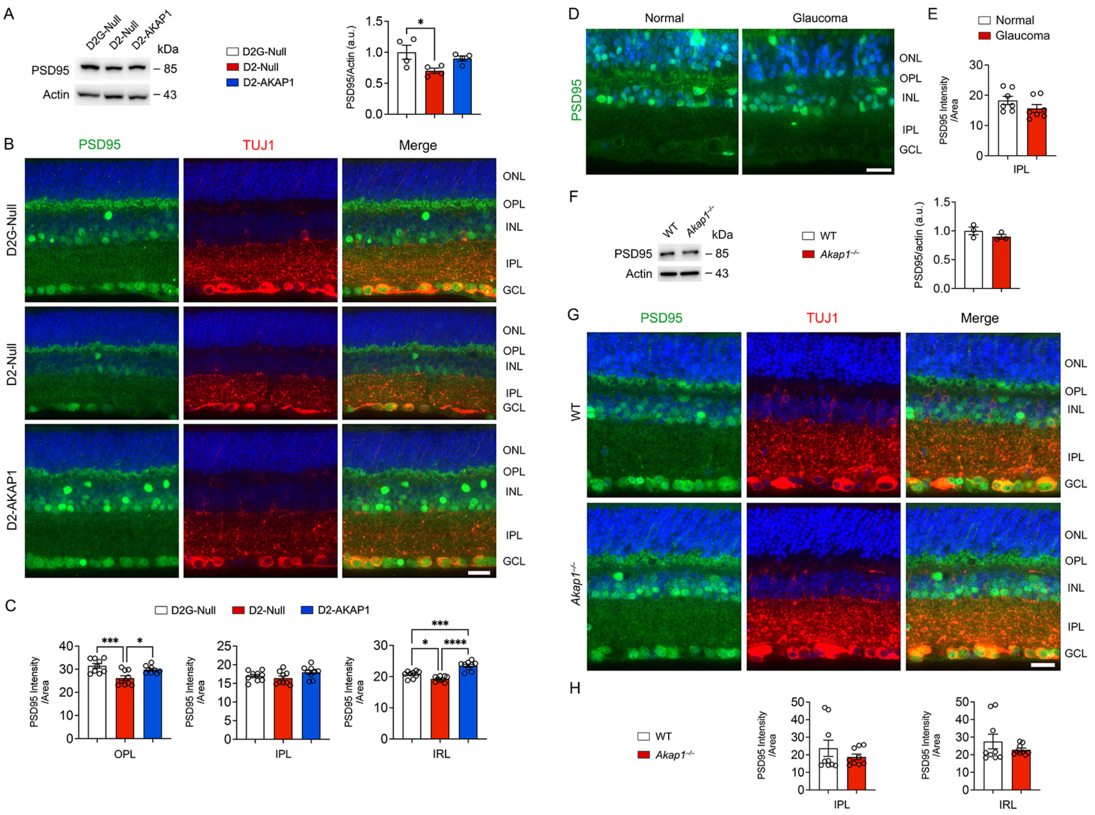

**Fig. S17** Restoring AKAP1 expression preserves PSD95 in glaucomatous D2 retina. (A) Western blot analysis for PSD95 expression in D2G and D2 retinas ($n$ = 4 retinas per group). (B) Representative retinal images for PSD95 immunoreactivity in D2G and D2 retinas. (C) Quantitative fluorescent intensity of PSD95 immunoreactivity in the OPL, IPL, and inner retinal layer of D2G and D2 retinas ($n$=9 retinal sections from 3 mice per group). (D) Representative retinal images for PSD95 (green) immunoreactivity in normal and glaucomatous human retinas. (E) Quantitative fluorescent intensity of PSD95 immunoreactivity in the IPL of the glaucomatous human retinas ($n$ = 7 retinal sections from 7 eyes from donors/group). (F) Western blot analysis for PSD95 expression in WT and *Akap1*$^{-/-}$ retinas ($n$ = 3 retinas per group). (G) Representative retinal images for PSD95 immunoreactivity in WT and *Akap1*$^{-/-}$ retinas. (H) Quantitative fluorescent intensity of PSD95 immunoreactivity in the IPL and inner retinal layer (IRL) of the WT and *Akap1*$^{-/-}$ retinas ($n$=9 retinal sections from 3 mice per group). Bars represent SEM. Statistical analysis was performed using unpaired Student's $t$-test or one-way ANOVA and Tukey's multiple comparisons test. *$P$ < 0.05, ***$P$ < 0.001, and ****$P$ < 0.0001. Scale bars, 50 µm.

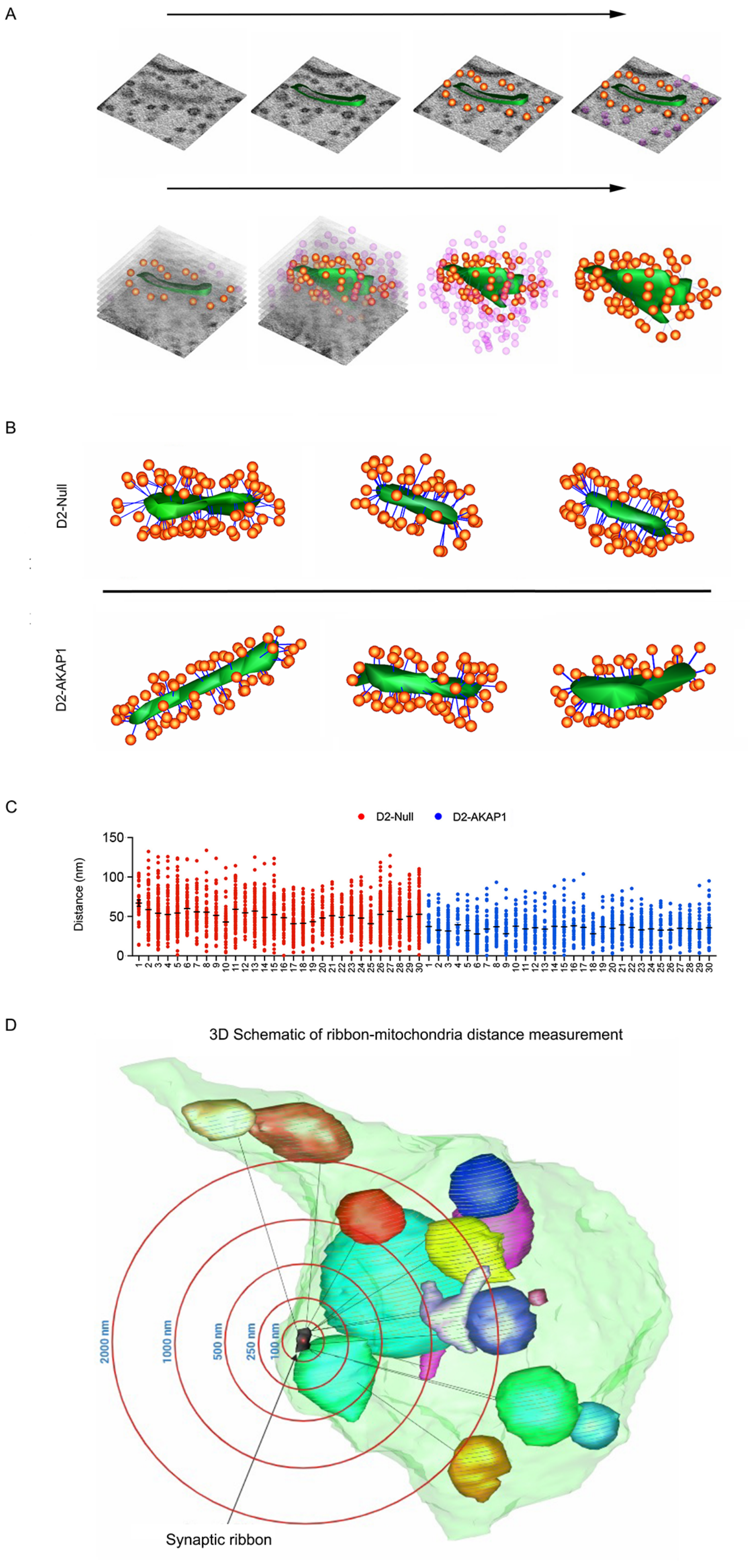

**Fig. S18** Restoring AKAP1 expression preserves presynaptic function by increasing the number of ribbon synapses and synaptic vesicles in glaucomatous D2 retina. (A) Representative SBEM images for synaptic bouton (gray), synaptic ribbon (green), and synaptic vesicles (orange dots) of D2-Null and D2-AKAP1 retina. (B) Representative SBEM images of synaptic ribbon (green) and synaptic vesicle (orange) in the synaptic bouton of D2-Null and D2-AKAP1 retina. (C) Quantitative analysis of synaptic vesicle distance from synaptic ribbons in D2-Null and D2-AKAP1 retinas. (D) 3D schematic for measurements of ribbon-mitochondria distance.

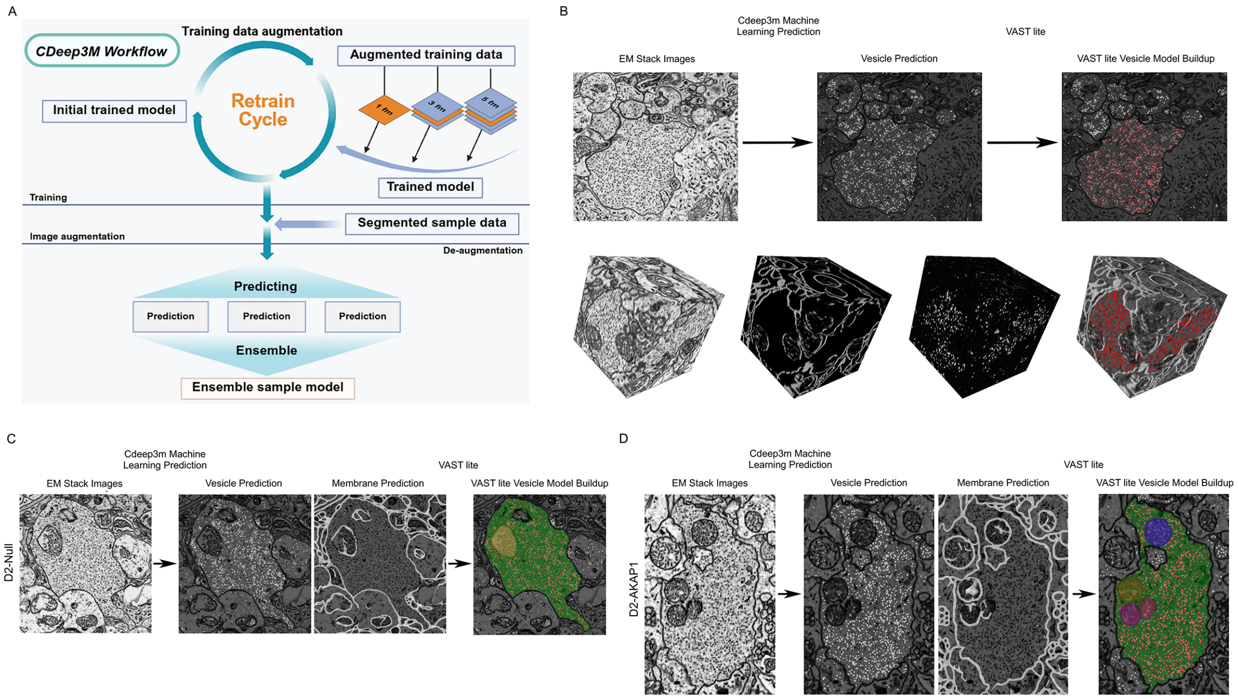

**Fig. S19** Workflow of Synaptic Vesicle Segmentation Using CDeep3M and VAST Lite. (A) Schematic summary for the AI-based segmentation tool CDeep3M workflow to facilitate deep learning–based image segmentation of 3D image stacks. (B) Top row: Representative 2D EM image from a stack (left) is used as input for machine learning-based segmentation using CDeep3M. The pipeline predicts synaptic vesicles (middle), which are then used as input for model building in VAST Lite (right). The overlay in the final image shows vesicles segmented in red. Bottom row: 3D renderings illustrate the pipeline progression. From left to right: raw EM image volume, membrane prediction, vesicle prediction, and automated vesicle segmentation. Red spheres denote segmented synaptic vesicles within the synaptic boutons. (C) D2-Null. 3D renderings illustrate the pipeline progression. From left to right: EM stack images, vesicle prediction, membrane prediction, and VAST Lite vesicle model buildup. The overlay in the final image shows vesicles segmented in red and mitochondria in several colors. (D) D2-AKAP1. 3D renderings illustrate the pipeline progression. From left to right: EM stack images, vesicle prediction, membrane prediction, and VAST Lite vesicle model buildup. The overlay in the final image shows vesicles segmented in red and mitochondria in several colors.

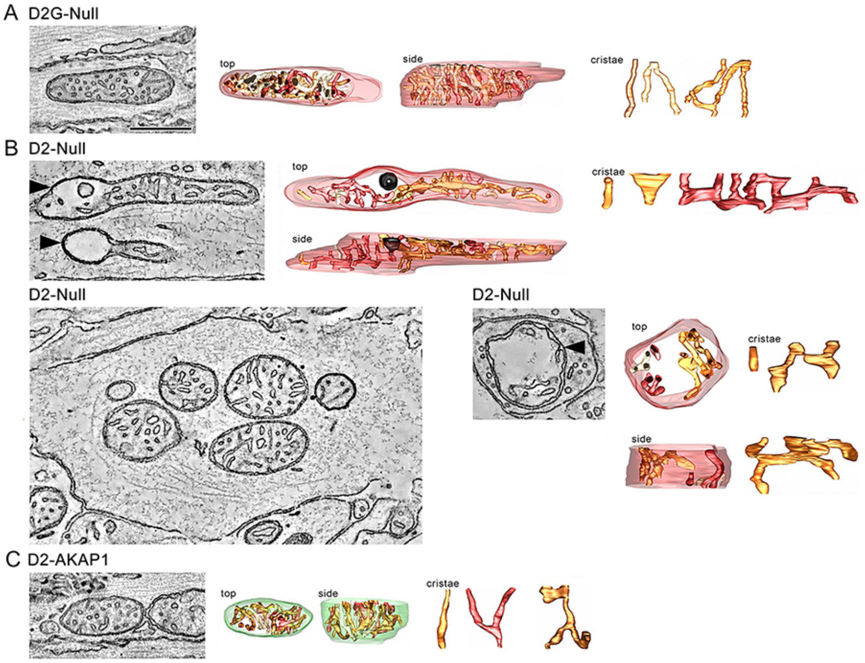

**Fig. S20** Restoring AKAP1 expression protects ONH axons by promoting mitochondrial fusion activity in the glaucomatous D2 mice. (A) Slice through the middle of an EM tomography volume of a D2G-Null ONH axon mitochondrion. Top and side views of the surface-rendered mitochondrial volume after membrane segmentation. All the cristae have a lamellar shape. (B) Slice through the middle of an EM tomography volume of D2-Null ONH axon mitochondria. Top and side views of the surface-rendered mitochondrial volume after membrane segmentation. All the cristae have a lamellar shape. (C) Slice through the middle of an EM tomography volume of D2-AKAP1 ONH axon mitochondria. Top and side views of the surface-rendered mitochondrial volume after membrane segmentation. All the cristae have a lamellar shape.

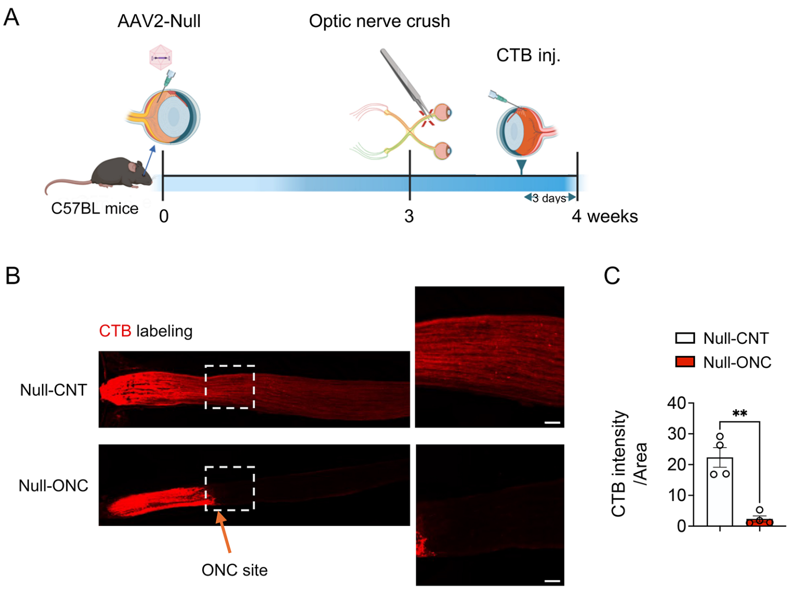

**Fig. S21** ONC modeling for AAV2 vector transduction. (A) Experimental schematic and timeline of AAV2 vector transduction and CTB labeling. (B) Representative images for CTB labeling in the optic nerve following ONC. (C) Quantitative analysis of CTB intensity in the ON ($n$ = 4 ONs from 4 mice per group). Error bars represent SEM. Statistical analysis was performed using an unpaired Student's $t$-test. $^{**}P < 0.01$. Scale bars, 50 μm.

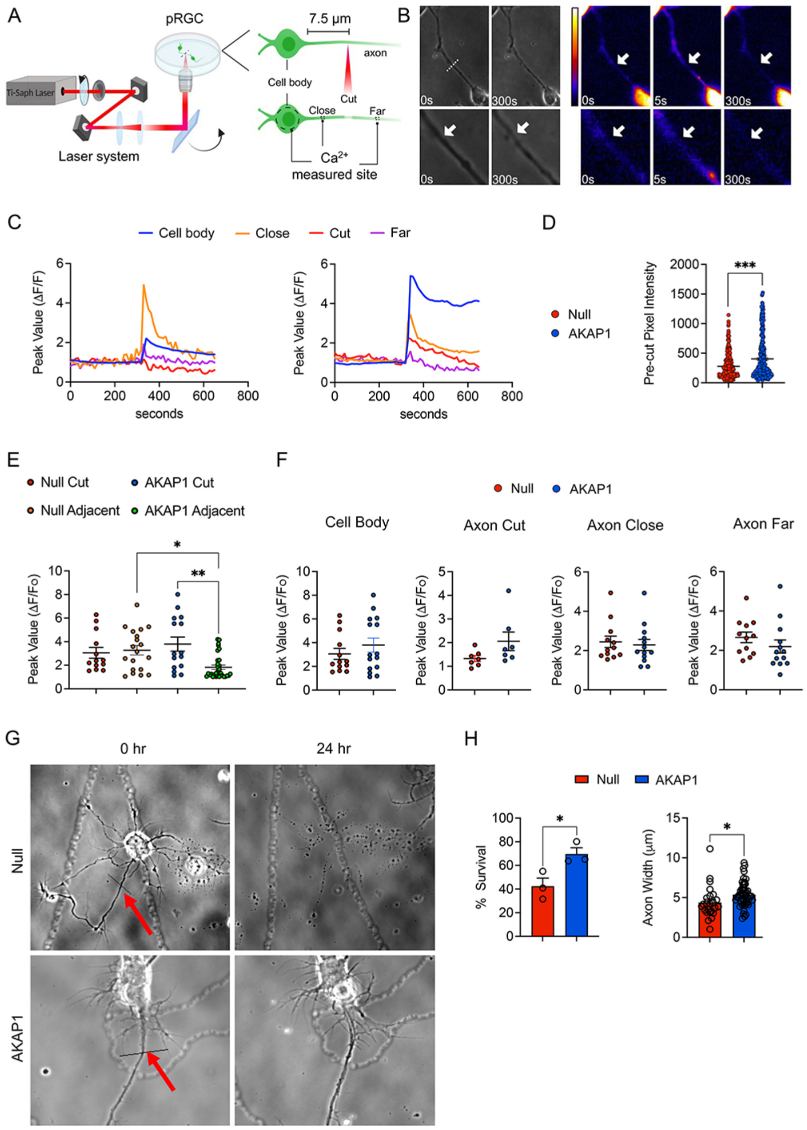

**Fig. S22** (A) Experimental schematic for an *in vitro* model of laser ablation. (B) Representative phase images. (C) Initial calcium levels within the cell body, and on the axon at varying distances from the cell body (close, cut, far). (D) Baseline calcium levels. (E) Calcium response. (F) Axonal calcium injury response by distribution. (G) Representative images for RGC survival. (H) Quantification of RGC survival following laser ablation. Bars represent SEM. Statistical analysis was performed using unpaired Student's *t*-test or one-way ANOVA and Tukey's multiple comparisons test. *$P < 0.05$, **$P < 0.01$, and ***$P < 0.001$.

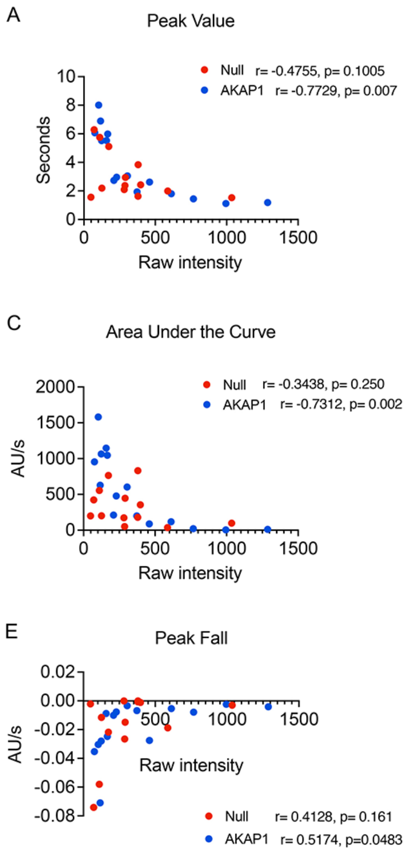

**Fig. S23** Correlative analysis of calcium signals in RGC and models. (A-E) Quantitative analysis of calcium signal intensity. Statistical analysis was performed using an unpaired Student's *t*-test.

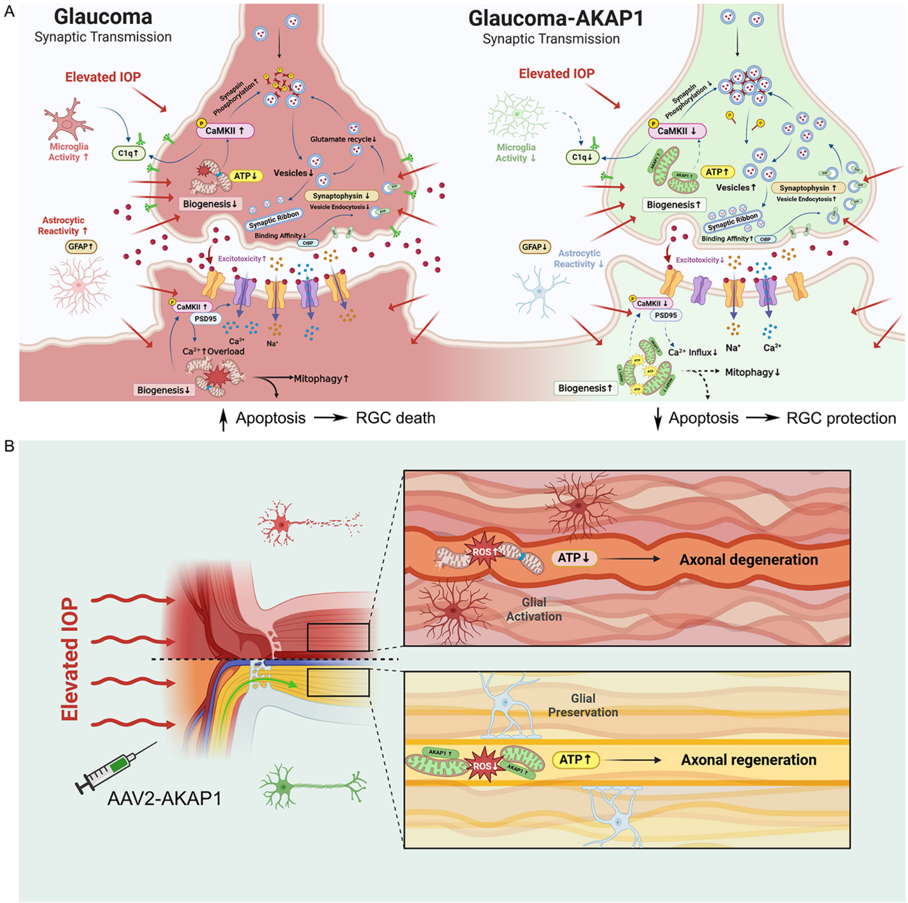

**Fig. S24** Schematic of the retinal synapse and ON axons in glaucomatous retina and ON by AAV2-AKAP1 transduction. (A) Schematic summary for structural and functional preservation of mitochondria and synapses in glaucomatous retina. (B) Schematic summary for structural and functional preservation of mitochondria and axonal regeneration in glaucomatous ONH.

### Supplemental Tables

**Table S1.** Effect of AAV2-AKAP1 on RBPMS-positive RGC survival in the middle and peripheral retina from 10-month-old glaucomatous DBA/2J mice, related to Figure 2K.

| Strain (DBA/2J) | Treatment | Age (Months) | Mean IOP (mmHg) at age of 10 months | RGC density per retina ($RGCs/mm^2$) | |
|---|---|---|---|---|---|
| | | | | Middle | Peripheral |
| D2G-Null | AAV2-Null | 10 | 10.7 ± 0.4 | 3427 ± 66 | 2853 ± 68 |
| D2-Null | AAV2-Null | 10 | 26.9 ± 0.6 | 1915 ± 197 | 1474 ± 138 |
| D2-AKAP1 | AAV2-AKAP1 | 10 | 26.5 ± 0.5 | 2883 ± 55 | 2360 ± 75 |

All results were reported as means ± SEM. *n* = 9-18 retina wholemounts from 5-15 mice per group.

**Table S2.** Effect of AAV2-AKAP1 on RBPMS-positive RGC survival in the middle/peripheral retina from glaucomatous mice induced by microbead (MB)-induced ocular hypertension, related to Figure 3C.

| Strain (C57) | Treatment | Age (Months) | Mean IOP (mmHg) at 8 weeks of MB | RGC density per retina ($RGCs/mm^2$) | |
|---|---|---|---|---|---|
| | | | | Middle | Peripheral |
| Null-CNT | AAV2-Null | 5 | 13 ± 0.4 | 3496 ± 70 | 2709 ± 72 |
| Null-MB | AAV2-Null | 5 | 26.2 ± 1 | 2805 ± 66 | 2318 ± 62 |
| AKAP1-MB | AAV2-AKAP1 | 5 | 26.4 ± 1.3 | 3503 ± 111 | 2849 ± 113 |
| AKAP1-CNT | AAV2-AKAP1 | 5 | 12.2 ± 0.5 | 3644 ± 54 | 2860 ± 41 |

All results were reported as means ± SEM. *n* = 7-10 retina wholemounts from 7-10 mice per group.

**Table S3.** Effect of AKAP1 deficiency on RBPMS-positive RGC survival in middle and peripheral retina from glaucomatous mice induced by microbead (MB)-induced ocular hypertension, related to Figure 3I.

| Strain (C57BL/6J and *Akap1*$^{-/-}$) | Age (Months) | Mean IOP (mmHg) at 8 weeks of MB | RGC density per retina ($RGCs/mm^2$) | |
|---|---|---|---|---|
| | | | Middle | Peripheral |
| WT-CNT | 5 | 13 ± 0.3 | 2861 ± 108 | 2569 ± 68 |
| WT-MB | 5 | 22.9 ± 0.5 | 2673 ± 70 | 2191 ± 57 |
| *Akap1*$^{-/-}$ -MB | 5 | 27.8 ± 1.3 | 2693 ± 84 | 2113 ± 68 |
| *Akap1*$^{-/-}$ -CNT | 5 | 13.1 ± 0.4 | 3122 ± 49 | 2673 ± 48 |

All results were reported as means ± SEM. *n* = 8 retina wholemounts from 8 mice per group.

**Table S4.** Effect of AAV2-AKAP1 on RBPMS-positive RGC survival in the middle and peripheral retina from a mouse induced by optic nerve crush injury, related to Figure 8D.

| Strain (C57BL/6J) | Age (Months) | RGC density per retina (RGCs/mm$^2$) | |
|---|---|---|---|
| | | Middle | Peripheral |
| Null-CNT | 4 | 3164 ± 107 | 2545 ± 105 |
| Null-ONC | 4 | 1361 ± 59 | 1092 ± 78 |
| AKAP1-ONC | 4 | 1639 ± 43 | 1425 ± 56 |
| AKAP1-CNT | 4 | 3364 ± 84 | 2785 ± 63 |

All results were reported as means ± SEM. *n* = 4-7 retina wholemounts from 4-7 mice per group.

**Table S5. Human tissue sample patient information**

| **SN** | **Patient Identification number** | **Age** | **Gender** | **Condition** |
|---|---|---|---|---|
| 1 | W4129-24-000586 | 81 | M | Normal |
| 2 | W4129-24-002012 | 83 | F | Normal |
| 3 | W4129-24-001150 | 66 | F | Normal |
| 4 | W4129-23-002157 | 75 | F | Normal |
| 5 | W4129-21-007052 | 84 | F | Glaucoma |
| 6 | W4129-23-002111 | 90 | F | Glaucoma |
| 7 | W4129-24-001609 | 77 | F | Glaucoma |
| 8 | W4129-24-002631 | 93 | F | Glaucoma |

**Table S6. Key Resource**

| **REAGENT OR RESOURCES** | **SOURCE** | **IDENTIFIER** |
|---|---|---|
| Antibodies | Supplier | Catalogue number |
| AKAP1 | Cell Signaling Technology | CST #5203 |
| AMPK | Cell Signaling Technology | CST #5831 |
| Phospho-AMPK | Cell Signaling Technology | CST #2535 |
| β-ACTIN | Millipore | MAB1501 |
| Active BAX | Santa Cruz Biotechnology | sc-23959 |
| BCL-xL | Santa Cruz Biotechnology | sc-8392 |
| CaN | BD Bioscience | #610259 |
| pCaMKII | Abcam | Ab5683 |
| CtBP | Santa Cruz | Sc-17759 |
| C1q | Quidel Corporation | A301 |
| Total DRP1 | BD Biosciences | #611113 |
| Phospho-DRP1 S616 | Cell Signaling Technology | CST #3455 |
| Phospho-DRP1 S637 | Cell Signaling Technology | CST #4867 |
| EAAT1 | Abcam | Ab181036 |
| GFAP | Invitrogen | 13-0300 |
| GSK3beta | Cell Signaling Technology | CST #12456 |
| Phospho-GSK3b S9 | Cell Signaling Technology | CST #5558 |
| Anti-6X His tag | Abcam | Ab245114 |

| | | |
|---|---|---|
| 6x-His Tag | Invitrogen | MB1-21315 |
| IBA1 | Wako Chemicals | 016-20001 |
| IL1b | Bio-techne | NB600-633 |
| LC3 | Cell Signaling Technology | CST #2775 |
| MFN1 | Abcam | ab57602 |
| MFN2 | Abcam | ab56889 |
| NF68 | Sigma | N5139 |
| OPA1 | BD Biosciences | #612607 |
| OXPHOS | Invitrogen | #458099 |
| p38 | Cell Signaling Technology | CST #8690 |
| Phospho-p38 | Cell Signaling Technology | CST #4511 |
| p62 | MBL International | PM045 |
| PGC-1alpha | Santa Cruz Biotechnology | sc-13067 |
| PKA | Santa Cruz Biotechnology | sc-28315 |
| PSD95 | Abcam | Ab18258 |
| IBA1 | Wako | 019-19741 |
| RBPMS | Novus Biologicals | NBP2-20112 |
| pSynapsin | Cell Signaling Technology | CST 2311S |
| Synapsin | EMD Millipore | AB1543 |
| Synaptophysin | GeneTex | GTX633821 |
| TFAM | GeneTex | GTX77852 |
| TUJ1 | BioLegend | 801202 |
| VDAC | Cell Signaling Technology | CST #4866S |
| Goat anti-rabbit HRP | Cell Signaling Technology | 7074 |
| Goat anti-mouse HRP | Cell Signaling Technology | 7076 |
| Alexa Fluor-488 conjugated donkey anti-mouse IgG antibody | Invitrogen | A-21203 |
| Alexa Fluor-568 conjugated donkey anti-mouse IgG antibody | Invitrogen | A-10037 |
| Alexa Fluor-488 conjugated donkey anti-rabbit IgG antibody | Invitrogen | A-21206 |
| Alexa Fluor-568 conjugated donkey anti-rabbit IgG antibody | Invitrogen | A-10042 |
| Alexa Fluor 594-conjugated CTB | ThermoFisher Scientific | C34777 |
| Bacterial and Virus Strains | | |
| AAV2-M4-AKAP1-His | | N/A |
| **Biological Samples** | | |
| Human retina tissue samples | San Diego Eye Bank | N/A |
| **Chemicals, Reagent and Commercial Kits** | **Supplier** | **Catalogue number** |
| Ketamine | Ketaset | |

| | | |
|---|---|---|
| MTT | Roche | 11465007001 |
| MitoTraker Red | Invitrogen | |
| Paraformaldehyde | Sigma-Aldrich | |
| Paraquat | Sigma-Aldrich | 36541-100mg |
| Proparacaine | | |
| Seahorse XF Cell Mito Stress Test Kit | Agilent | 103015-100 |
| SuperSignal Chemiluminescent | Thermo Scientific | 34580 |
| Topical phenylephrine | | |
| Tropicamide | | |
| Xylazine | TranquilVed | |
| **Experimental Models: Strains** | **Vendor** | |
| C57BL/6J mice | Jackson laboratory | Stock No: 000664 RRID:IMSR_JAX:000664 |
| DBA/2J mice | Jackson laboratory | Stock No: 000671 RRID:IMSR_JAX:000671 |
| DBA/2J-*Gpnmb*$^{+}$/SjJ mice | Jackson laboratory | Stock No: 007048 RRID:IMSR_JAX:007048 |
| *Akap1*$^{-/-}$ mice | | |
| Sprague-Dalwley Rat | Envigo (Harlan) | N/A |
| **Software and Algorithms** | **Suppliers** | **Site Link** |
| ImageJ | NIH, USA | https://imagej.nih.gov/ij/ |
| Adobe Photoshop | USA | https://photoshop.com |
| Prism 9 | GraphPad Inc. | https://www.graphpad.com/scientific-software/prism/ |

**Table S7. Primer list used in the experiment**

| SN | Primer list | Sequence (F) | Sequence (R) |
|---|---|---|---|
| 1 | AKAP1 | AAGCTATGACCCCACCACTG | CGCAACAGCTATCCACTGAA |
| 2 | DRP1 | AGGTGGCCTTAACACTATTGACA | AGACGCTTAATCTGACGTTTGAC |
| 3 | OPA1 | CATCTACCTTCCAGCTGCCC | TCTCCTCCTTCACAGCCTCC |
| 4 | MFN1 | GTGAAGTTCACAAGTGCAAA | GCTCGGGTGGAGAAACTGCT |
| 5 | MFN2 | CCCTCGACAGTGTTTCTCCC | CCAGGCCAGTAACCATGGAG |
| 6 | TFAM | CCAAAAAGACCTCGTTCAGC | ATGTCTCCGGATCGTTTCAC |
| 7 | PGC-1a | AAGGTCCCCAGGCAGTAGAT | GCGGTATTCATCCCTCTTGA |
| 8 | GFAP | GAAAGGTTGAATCGCTGGAG | GCCACTGCCTCGTATTGAGT |

| | | | |
|---|---|---|---|
| 9 | IBA1 | GGACAGACTGCCAGCCTAAG | GACGGCAGATCCTCATCATT |
| 10 | IL-1β | GGAGAACCAAGCAACGACAAA ATA | TGGGGAACTCTGCAGACTCAA AC |
| 11 | TNFα | CATCTTCTCAAAATTCGAGTGAC | TGGGAGTAGACAAGGTACAAC CC |
| 12 | Caspase 1 | AGGAATTCTGGAGCTTCAATCAG | TGG AAA TGT GCC ATC TTC TTT |
| 13 | NLRP3 | GCTCCAACCATTCTCTGACC | AAGTAAGGCCGGAATTCACC |
| 14 | GAPDH | AGAACATCATCCCTGCATCC | GTCCTCAGTGTAGCCCAAGA |

**Supplemental Videos**

**Video S1.** Null-NP, Surface-rendered mitochondrial volume after membrane segmentation, related to Figure 4M.

**Video S2.** Null-EHP, Surface-rendered mitochondrial volume after membrane segmentation, related to Figure 4M.

**Video S3.** Null-AKAP1, Surface-rendered mitochondrial volume after membrane segmentation, related to Figure 4M.

**Video S4.** D2-Null, Surface-rendered mitochondrial volume and synaptic bouton after membrane segmentation, related to Figure 5J.

**Video S5.** D2-AKAP1, Surface-rendered mitochondrial volume and synaptic bouton after membrane segmentation, related to Figure 5J.

**Video S6.** D2-Null, Surface-rendered ribbon synapses, synaptic vesicles, and spinules in the synaptic bouton after membrane segmentation, related to Figure 6F.

**Video S7**. D2-AKAP1, Surface-rendered ribbon synapses, synaptic vesicles, and spinules in the synaptic bouton after membrane segmentation, related to Figure 6G.

**Video S8.** D2-Null, Surface-rendered synaptic ribbon and vesicles in the synaptic bouton, related to Figure 6F.

**Video S9**. D2-AKAP1, Surface-rendered synaptic ribbon and vesicles in the synaptic bouton, related to Figure 6F.

**Video S10.** D2-Null, Surface-rendered mitochondria and synaptic vesicles using CDeep3M and Vast Lite, related to Figure 6L.

**Video S11.** D2-AKAP1, Surface-rendered mitochondria and synaptic vesicles using CDeep3M and Vast Lite, related to Figure 6L.

**Video S12.** D2G-Null, Surface-rendered mitochondrial volume after membrane segmentation, related to Figure 8E.

**Video S13.** D2-Null, Surface-rendered mitochondrial volume after membrane segmentation, related to Figure 8F.

**Video S14.** D2-Null, Surface-rendered mitochondrial volume after membrane segmentation, related to Figure 8F.

**Video S15.** D2-AKAP1, Surface-rendered mitochondrial volume after membrane segmentation, related to Figure 8G.
